# CCQM-P199b: Interlaboratory comparability study of SARS-CoV-2 RNA copy number quantification

**DOI:** 10.1101/2024.03.27.584106

**Authors:** Alison S. Devonshire, Eloise J. Busby, Gerwyn M. Jones, Denise M. O’Sullivan, Ana Fernandez-Gonzalez, Laura Hernandez-Hernandez, Xinhua Dai, Lianhua Dong, Chunyan Niu, Jie Xie, Xia Wang, Xiaoting Qiao, Xiang Fang, Clare Morris, Neil Almond, Megan H. Cleveland, Peter M. Vallone, Esther Castro Galván, Melina Pérez Urquiza, Mercedes Guadalupe Herrera López, Arifa S. Khan, Sandra M. Fuentes, John Emerson Leguizamon Guerrero, Sergio Luis Davila Gonzalez, Andres Felipe León Torres, Aurea V Folgueras-Flatschart, Marcelo Neves de Medeiros, Antonio Marcos Saraiva, Roberto Becht Flatschart, Carla Divieto, Mattia Pegoraro, Massimo Zucco, Laura Revel, Marco Mazzara, Philippe Corbisier, Gerhard Buttinger, Inchul Yang, Young-Kyung Bae, Alexandra Bogožalec Košir, Mojca Milavec, Malcolm Hawkins, A. Pia Sanzone, Phattarapornn Morris, Sasithon Temisak, David Lynch, Jacob McLaughlin, Michael Forbes-Smith, Felicity Hall, Daniel Burke, Sachie Shibayama, Shin-ichiro Fujii, Megumi Kato, Samreen Falak, Rainer Macdonald, Andreas Kummrow, Andrey Komissarov, Kseniya Komissarova, Sema Akyurek, Muslum Akgoz, Sumeyra Nur Sanal Demirci, Maxim Vonsky, Andrey Runov, Elena Kulyabina, Denis Rebrikov, Jim F. Huggett

## Abstract

Nucleic acid amplification tests including reverse transcription quantitative PCR (RT-qPCR) are used to detect RNA from Severe Acute Respiratory Syndrome Coronavirus 2 (SARS-CoV-2), the causative agent of the Coronavirus disease 2019 (COVID-19) pandemic. Standardized measurements of RNA can facilitate comparable performance of laboratory tests in the absence of existing reference measurement systems early on in a pandemic. Interlaboratory study CCQM P199b “SARS-CoV-2 RNA copy number quantification” was designed to test the fitness-for-purpose of developed candidate reference measurement procedures (RMPs) for SARS-CoV-2 genomic targets in purified RNA materials, and was conducted under the auspices of the Consultative Committee for Amount of Substance: Metrology in Chemistry and Biology (CCQM) to evaluate the measurement comparability of national metrology institutes (NMIs) and designated institutes (DIs), thereby supporting international standardization.

Twenty-one laboratories participated in CCQM P199b and were requested to report the RNA copy number concentration, expressed in number of copies per microliter, of the SARS-CoV-2 nucleocapsid (*N*) gene partial region (NC_045512.2: 28274-29239) and envelope (*E*) gene (NC_045512.2: 26245-26472) (optional measurements) in samples consisting of *in vitro* transcribed RNA or purified RNA from lentiviral constructs. Materials were provided in two categories: lower concentration (≈ (10^1^-10^4^) /μL in aqueous solution containing human RNA background) and high concentration (≈ 10^9^ /μL in aqueous solution without any other RNA background).

For the measurement of *N* gene concentration in the lower concentration study materials, the majority of laboratories (*n* = 17) used one-step reverse transcription-digital PCR (RT-dPCR), with three laboratories applying two-step RT-dPCR and one laboratory RT-qPCR. Sixteen laboratories submitted results for *E* gene concentration. Reproducibility (% CV or equivalent) for RT-dPCR ranged from 19 % to 31 %. Measurements of the high concentration study material by orthogonal methods (isotope dilution-mass spectrometry and single molecule flow cytometry) and a gravimetrically linked lower concentration material were in a good agreement, suggesting a lack of overall bias in RT-dPCR measurements. However methodological factors such as primer and probe (assay) sequences, RT-dPCR reagents and dPCR partition volume were found to be potential sources of interlaboratory variation which need to be controlled when applying this technique.

This study demonstrates that the accuracy of RT-dPCR is fit-for-purpose as a RMP for viral RNA target quantification in purified RNA materials and highlights where metrological approaches such as the use of *in vitro* transcribed controls, orthogonal methods and measurement uncertainty evaluation can support standardization of molecular methods.

## INTRODUCTION

RNA is the analyte targeted by nucleic acid amplification tests for Severe Acute Respiratory Syndrome Coronavirus 2 (SARS-CoV-2), the causative agent of the Coronavirus disease 2019 (COVID-19) pandemic. RNA concentration measured in respiratory specimens can vary by over nine orders of magnitude [1, 2]. Reverse transcription-quantitative PCR (RT-qPCR) provided the main diagnostic method for the management of COVID-19 as the pandemic spread in early 2020. The use of RT-qPCR varied considerably with both in-house/laboratory developed tests widely applied [3] and a range of commercially available *in vitro* diagnostic solutions being deployed. However considerable variation was observed in analytical performance [4], calling for standards to demonstrate conformity of an *in vitro* diagnostic solutions to stipulated limit of detection targets [5–8]. While RNA abundances were not generally used to guide treatment, quantitative thresholds were proposed to better stratify patients in terms of clinical relevance [9, 10]. However despite being a quantitative approach, SARS-CoV-2 diagnostic RT-qPCR results are often reported in arbitrary quantification cycle (C_q_) values (also termed cycle threshold or C_t_), which prevent meaningful comparison between laboratories, assays and instruments and over time (internal QC) [11].

In response to these standardisation challenges, reference materials (RMs) were developed, which include materials based on DNA or RNA constructs, for example, NIST RGTM 10169 [12], or whole virus preparations such as the first World Health Organization (WHO) International Standard for SARS-CoV-2 RNA [13]. Reverse transcription-digital PCR (RT-dPCR) was implemented as a candidate reference measurement procedure (RMP) and employed during the COVID-19 pandemic in External Quality Assurance schemes and for value assignment of Reference Materials [14].

In April 2020, the Consultative Committee for Amount of Substance: Metrology in Chemistry and Biology (CCQM) approved the Pilot study CCQM-P199b “SARS-CoV-2 RNA copy number quantification”. CCQM-P199b was designed to assess participants’ capabilities for targeted RNA copy number concentration and viral gene quantification. CCQM-P199b included multiple measurands (the SARS-CoV-2 nucleocapsid (*N*) and envelope (*E*) gene targets), while also assessing NMI/DI capacity to respond quickly to implement reference measurement procedures (RMPs) in support of rapidly deployed IVDs. CCQM-P199b was fast-tracked (material shipments organized between May and July 2020, with results submitted October 2020) to provide NMIs with the tools to be able to respond to national needs in the global response to the COVID-19 pandemic.

CCQM-P199b sought to address the comparability of higher order methods for targeted RNA quantification relevant to the diagnostic range (≈10^1^ to ≈10^5^ /μL purified RNA equivalent to 10^2^ /mL to > 10^6^ /mL of respiratory specimen) and to establish routes for SI-traceability of counting-based approaches (RT-dPCR) through testing trueness by comparison with orthogonal methods such as isotope dilution mass spectrometry (ID-MS) and single molecule flow cytometry (SMFC) which do not require reverse transcription or amplification, enzymatic steps which may lead to biases in RT-dPCR [15]. To address the metrological state of the art, provide evidence for NMIs/DIs for value assignment of calibration materials and support analytical standardization of diagnostic methods, *in vitro* synthesised RNA templates and purified RNA extracts were chosen as the Study Materials for analysis.

The following sections of this report document the Measurands, Study Materials, participants, timeline, results, discussion of sources of interlaboratory variation and consensus RVs of CCQM-P199b. The Appendices (Supplementary materials) reproduce the official communication materials and summaries of information about the results provided by the participants.

## MEASURANDS

The measurands (quantity intended to be measured [16]) of CCQM-P199b are RNA copy number concentration of two SARS-CoV-2 genes / gene regions which are commonly targeted by molecular diagnostic tests: the *N* and *E* genes, which are towards 3’ end of the viral genome (Figure 1). The analyte is synthetic or purified RNA template. The matrix is buffered solution, with a complex (human total RNA) background (Study Materials 1-3).

- **Measurand 1:** RNA copy number concentration expressed in number of copies per µL of the *N* gene partial sequence (MN908974.3: 28274-29239);
- **Measurand 2:** RNA copy number concentration expressed in number of copies per µL of the *E* full gene sequence (MN908974.3: 26245-26472) (optional).

**Figure 1:**
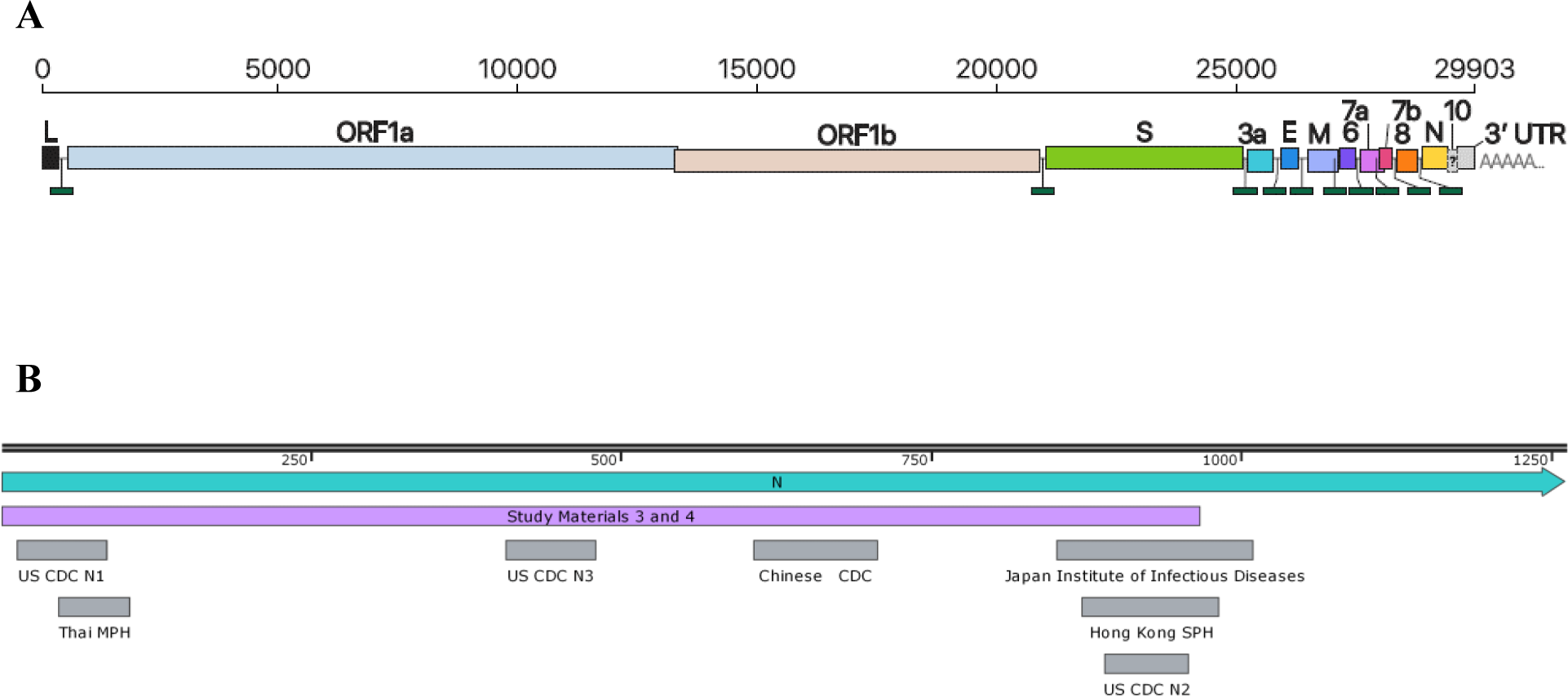
Schematic of the viral genome. (A) The structure of the SARS-CoV-2 genome [17]. (B) Relative locations of public health laboratory-developed assays (grey) to the *N* gene, showing Study Material 3 and 4 template region (purple).

Sequence information is provided in Appendix A.

## STUDY MATERIALS

### Background

Four Study Materials were designed and prepared by three of the coordinating laboratories: Study Material 1 (NIM China), Study Material 2 (NIBSC/NML), Study Materials 3 and 4 (NML). In addition to NML, NIST also evaluated Study Material 4 by RT-dPCR. Coordinating laboratory methodology is provided in Appendix B. All materials were synthetic or purified RNA, non-infectious and required bio-safety level 1 containment. Sequence information is summarised in Table 1 and RNA sequence information for each material is provided in Appendix A. It was expected that all study participants analyse Study Material 1, 2 and 3, whereas analysis of Study Materials 4 was optional. Study participants were provided with four units of each Study Material.

**Table 1:**
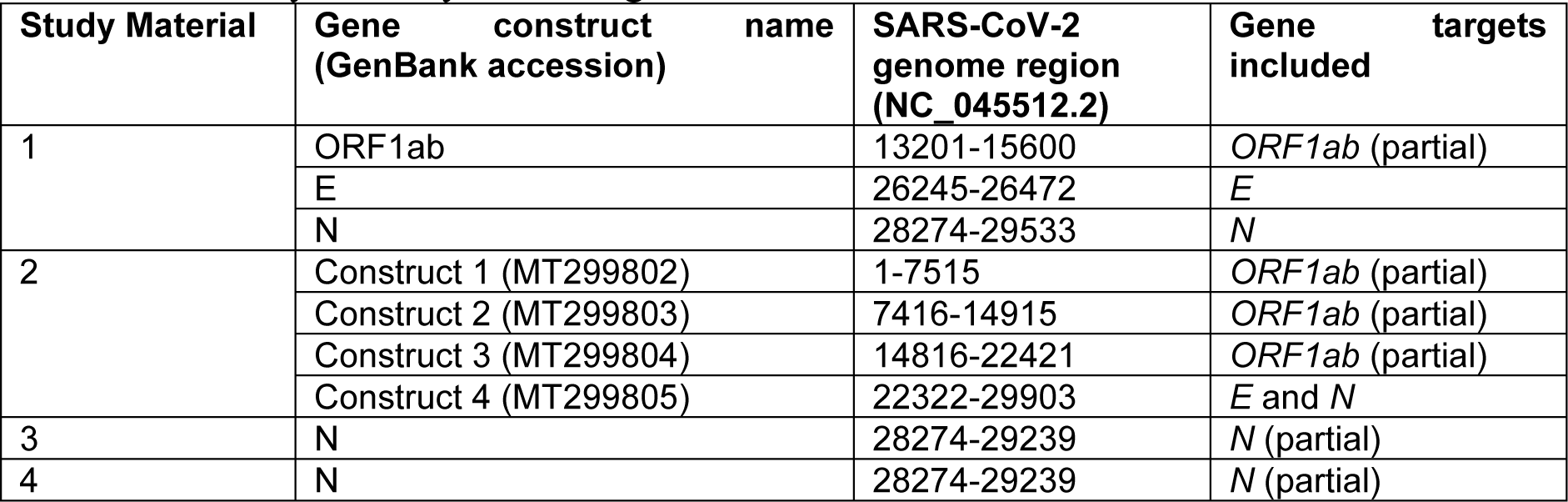
Summary of Study Material genomic information.

### Preparation of Study Materials

**Study Material 1** was composed of *in vitro* transcribed SARS-CoV-2 RNA segments of *ORF1ab*, complete *E* gene and complete *N* gene, at an approximate concentration of ≈ 10^3^ /μL in ≈ 5 ng/μL 293T human cell line total RNA (purchased from the National Infrastructure of Cell Line Resource and cultured by NIMC) in buffered solution (1 mM^1^ sodium citrate, pH 6.5 (RNA Storage Solution Thermo Fisher Scientific P/N AM7001)). *In vitro* transcription reaction was performed using MEGAscript T7 Transcription Kit (AM1334, Thermo Fisher Scientific, USA). RNA transcripts were purified with MEGAclearKit (Thermo Fisher scientific, USA). Further information on Study Material preparation is provided in Appendix B. A total of 200 units, each containing 100 µL, were prepared.

**Study Material 2** was prepared from NIBSC Research Reagent for SARS-CoV-2 RNA (NIBSC code 19/304; https://www.nibsc.org/documents/ifu/19-304.pdf)) which consists of four lentiviral constructs spanning the SARS-CoV-2 genome (for further information, see Appendix C). RNA was purified from two pooled vials each containing 0.5 mL of NIBSC Research Reagent for SARS-CoV-2 using the QIAamp UltraSens Virus (Qiagen) at NIBSC. A single vial of approximately 250 µL of RNA was thawed at the NML and a 10 µL aliquot removed for initial RT-dPCR analysis, following which 213 µL of the RNA stock was used for preparation of Study Material 2. Study Material 2 contained an approximate RNA copy concentration of ≈ 10^1^ /μL for both Measurands in ≈ 5 ng/μL FirstChoice Human T-Cell Leukemia (Jurkat) Total RNA (Ambion P/N AM7858) in RNA Storage Solution (as above). Initial measurement of a 20-fold volumetric dilution of Human T-Cell Leukaemia RNA concentration was performed using a Qubit RNA BR kit, and the solution was subsequently gravimetrically diluted 10-fold in RNA Storage Solution. Each unit of material contains 100 μL sample. A total of 206 units were prepared.

**Study Material 3** was composed of a single *in vitro* transcribed SARS-CoV-2 RNA molecule of the partial *N* gene at a similar RNA copy number concentration to Study Material 1, in ≈ 2.5 ng/μL human Jurkat cell line total RNA (as above) in RNA Storage Solution (as above) and was prepared by gravimetric dilution of Study Material 4 using a Mettler Toledo XP205 balance to 5 decimal places. Following cleaning of the balance, linearity was tested using a set of laboratory standard weights covering the range 0.1 g to 200 g. Standard uncertainty of measurement for the balance was ± 0.000159 g (based on the calibration certificate). Further details can be found in Appendix B (Table B-6). A total of 310 units, each containing 100 µL, were prepared.

**Study Material 4** contained the same *in vitro* transcribed SARS-CoV-2 RNA molecule of the *N* gene as Study Material 3 at an approximate concentration of (10^9^ to 10^10^) /μL (≈ 0.5 to 5 ng/μL) in RNA Storage Solution (as above) in a volume of 100 μL per unit. A total of 58 units were prepared. Cellular RNA extracts were not added to this material as it was designed to be suitable for analysis using chemical methods and SMFC, where background nucleic acids will or may interfere with measurement.

### Homogeneity Assessment of Study Materials

The homogeneity of all Study Materials was evaluated by RT-dPCR analysis (Bio-Rad QX200). The detailed methods used by coordinator laboratories is described in Appendix B. The homogeneity of Study Materials 1 to 3 was assessed by performing replicate measurements (sub-samplings) of 10 units. For Study Material 1 homogeneity evaluation, triplicate measurements of the *E* and *N* genes were performed by NIM. For Study Materials 2 and 3 homogeneity studies, eight replicate measurements of the *N* gene were performed by the NML at LGC using the ‘N2’ assay developed by United States Centres for Disease Control and Prevention (US CDC) [18].

Due to the smaller number of units produced for Material 4, homogeneity was evaluated in ≈ 10 % of units in accordance with International Organization for Standardization (ISO) Guide 35 in which six units of Study Material 4 were assessed by performing duplicate gravimetric dilutions prepared using a Mettler Toledo XP205 balance) with triplicate RT-dPCR measurements of each dilution. The mass concentration of *N* IVT in Study Material 4 was also analysed by fluorimetric assay (Qubit RNA HS Assay, Invitrogen/Thermo Fisher Scientific) with 6 units (*n* = 3 assays).

#### Analysis and results of homogeneity studies

Study Material 1 homogeneity data were analysed by one-way ANOVA and between-unit standard deviation (*s*_bb_) calculated. Homogeneity data for Study Materials 2, 3 and 4 were analysed using R version 3.6.1 running inside RStudio version 1.2.5001, using mixed effects models with maximum likelihood estimation. Models with mean value and dPCR plate row and column as fixed effects, and between unit variation (and dilution, Material 4) as (nested) random effect(s) were evaluated, with the most appropriate model chosen based on Akaike Information Criterion (AIC). For Materials 2 and 4, the best model was with mean value as the only fixed effect. For Material 3, dPCR plate column was included as additional fixed effect. Between-unit standard deviations are reported in Table 3 and were concluded to be acceptable.

**Table 2:**
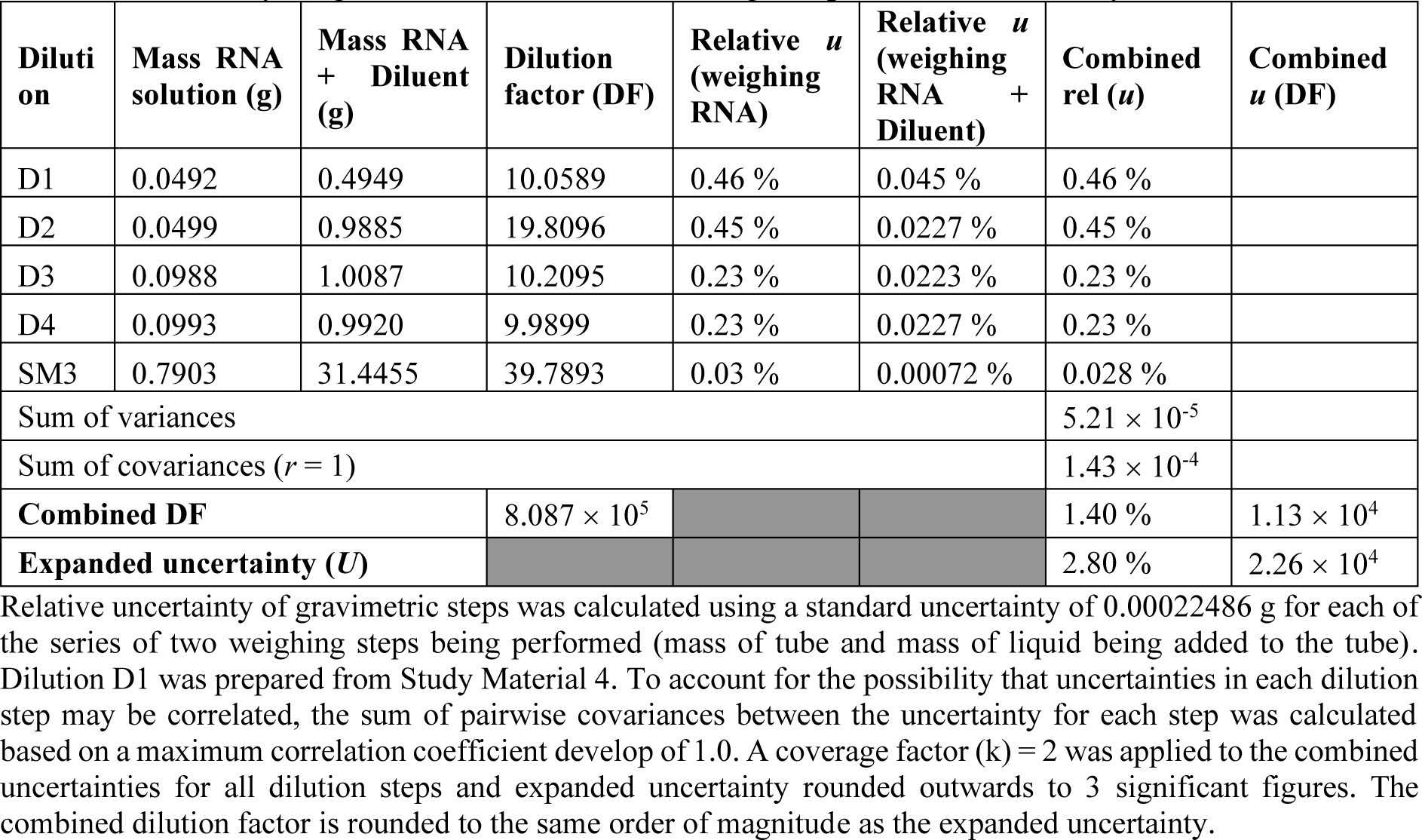
Summary of gravimetric dilutions linking the production of Study Materials 3-4.

| Dilution | Mass RNA solution (g) | Mass RNA + Diluent (g) | Dilution factor (DF) | Relative $u$ (weighing RNA) | Relative $u$ (weighing RNA + Diluent) | Combined rel ($u$) | Combined $u$ (DF) |
| --- | --- | --- | --- | --- | --- | --- | --- |
| D1 | 0.0492 | 0.4949 | 10.0589 | 0.46 % | 0.045 % | 0.46 % |  |
| D2 | 0.0499 | 0.9885 | 19.8096 | 0.45 % | 0.0227 % | 0.45 % |  |
| D3 | 0.0988 | 1.0087 | 10.2095 | 0.23 % | 0.0223 % | 0.23 % |  |
| D4 | 0.0993 | 0.9920 | 9.9899 | 0.23 % | 0.0227 % | 0.23 % |  |
| SM3 | 0.7903 | 31.4455 | 39.7893 | 0.03 % | 0.00072 % | 0.028 % |  |
| Sum of variances | | | | | | $5.21 \times 10^{-5}$ | |
| Sum of covariances ($r = 1$) | | | | | | $1.43 \times 10^{-4}$ | |
| **Combined DF** | | | $8.087 \times 10^5$ | | | 1.40 % | $1.13 \times 10^4$ |
| **Expanded uncertainty ($U$)** | | | | | | 2.80 % | $2.26 \times 10^4$ |

Relative uncertainty of gravimetric steps was calculated using a standard uncertainty of 0.00022486 g for each of the series of two weighing steps being performed (mass of tube and mass of liquid being added to the tube). Dilution D1 was prepared from Study Material 4. To account for the possibility that uncertainties in each dilution step may be correlated, the sum of pairwise covariances between the uncertainty for each step was calculated based on a maximum correlation coefficient develop of 1.0. A coverage factor ( $k$ ) = 2 was applied to the combined uncertainties for all dilution steps and expanded uncertainty rounded outwards to 3 significant figures. The combined dilution factor is rounded to the same order of magnitude as the expanded uncertainty.

**Table 3:** Study Material homogeneity results.

| Study Material (gene target measured) | Relative $s_{bb}$ |
| --- | --- |
| 1 (*N*) | 0.9 % |
| 1 (*E*) | 0.9 % |
| 2 (*N*) | 2.7 % |
| 3 (*N*) | 2.0 % |
| 4 (*N*) | 4.5 % |

### Stability Assessment of Study Material

#### Design of short-term stability studies

A short-term stability (STS) study was performed by incubation of Study Materials 1 to 3 on dry ice, at 4 °C and 27 °C for 3 days and 7 days (and 14 days, Study Material 1 only) and compared to reference temperature (−80 °C) (*n* = 3 units per condition). Due to the limitation in unit number, two units of Study Material 4 were placed on dry ice, at 4 °C or 27 °C for 7 days and compared to the reference temperature.

Short-term stability for all four Study Materials was assessed by RT-dPCR (*n* = 3). For Study Material 1 (NIMC), a duplex assay to *ORF1ab*/*E* was used. For Study Materials 2-4 (NML), the US CDC ‘N2’ assay was used. For Study Material 4, duplicate gravimetric dilution series were performed using an Ohaus E10640 balance to 4 decimal places with triplicate RT-dPCR measurements. Study Material 4 stability was also evaluated using the Total RNA Pico Kit with the Agilent 2100 Bioanalyzer to check for effects on the expected molecular size of 974 bp.

#### Results of short-term stability studies

Study Material 1 short-term stability study data were analysed using *T*-test with Bonferroni correction for multiple testing. Each storage temperature was compared with the reference temperature (−80 °C) with time points and units as a fixed effect. The magnitude and significance of the effects of storage temperature and duration are shown in Table 4. Study Material 1 showed a non-significant reduction in concentration at the 3 day timepoint at dry ice temperature, however this was interpreted as a random effect as no difference was found at 7 days. Concentration increased at the 7 day timepoint under 4 °C (*p* = *N.S*.), which was possibly caused by evaporation. Material 1 showed significant reductions in concentration following 3-day and 7-day incubation at 27 °C.

**Table 4:** Results of Short-Term Stability studies.

| Material | Storage condition | Time (days) | Relative change in concentration (value/value reference temperature) | *t*-value | Significant (*p*-value) * |
| --- | --- | --- | --- | --- | --- |
| 1 | 27 °C | 3 | -12 % |  | Yes (1.25E-4) |
|  | 27 °C | 7 | -10.4 % |  | Yes (3.48E-4) |
|  | 4 °C | 3 | 3 % |  | *N.S.* (0.15) |
|  | 4 °C | 7 | 8.6 % |  | *N.S.* (0.02) |
|  | Dry ice | 3 | -7.9 % |  | *N.S.* (0.02) |
|  | Dry ice | 7 | -2.9 % |  | *N.S.* (0.17) |
| 2 | 27 °C | 3 | -7 % | -0.85 | *N.S.* |
|  | 27 °C | 7 | -8 % | -1.00 | *N.S.* |
| | 4 °C | 3 | -15 % | -3.15 | Yes ($t > 3$) |
|  | 4 °C | 7 | 0 % | 0.06 | *N.S.* |
|  | Dry ice | 3 | -1 % | -0.12 | *N.S.* |
|  | Dry ice | 7 | -5 % | -0.52 | *N.S.* |
| 3 | 27 °C | 3 | -1 % | -1.31 | *N.S.* |
| | 27 °C | 7 | -7 % | -5.93 | Yes ($t > 3$) |
|  | 4 °C | 3 | 0 % | -0.55 | *N.S.* |
|  | 4 °C | 7 | 0 % | 0.04 | *N.S.* |
|  | Dry ice | 3 | 0 % | -0.27 | *N.S.* |
|  | Dry ice | 7 | 1 % | 0.51 | *N.S.* |
| 4 | 27 °C | 7 | -22 % | -8.08 | Yes ($t > 3$) |
|  | 4 °C | 18** | 2 % | 0.73 | *N.S.* |
|  | Dry ice | 7 | -1 % | -0.47 | *N.S.* |

\*Threshold for significance: $p = 0.0083$ ( $\alpha = 0.05$ , $n$ (number of tests) = 6)
\*\*Due to an error, Material 4 units stored at 4 °C were incubated for 18 days rather than 7 days.

For Study Materials 2, 3 and 4, short-term stability study data were analysed using mixed effects models with maximum likelihood estimation. Each storage temperature was compared with the reference temperature (−80 °C) with time points a fixed effects and unit (and dilution, Material 4) as random effect(s). The magnitude and significance of the effects of storage temperature and duration are shown in Table 4. Study Material 2 showed a reduction in concentration at the 3 day timepoint at 4 °C, however this was interpreted as a random effect as no difference was found at 7 days. Study Materials 3 and 4 showed significant reductions in concentration following 7 days incubation at 27 °C. Therefore, it was concluded that dry ice and 4 °C were suitable for shipment whilst ambient temperature was not. Study Material 4 stability was also evaluated by analysis of undiluted material by capillary gel electrophoresis (Figure 2), with no change in size of transcript observed.

**Figure 2:**
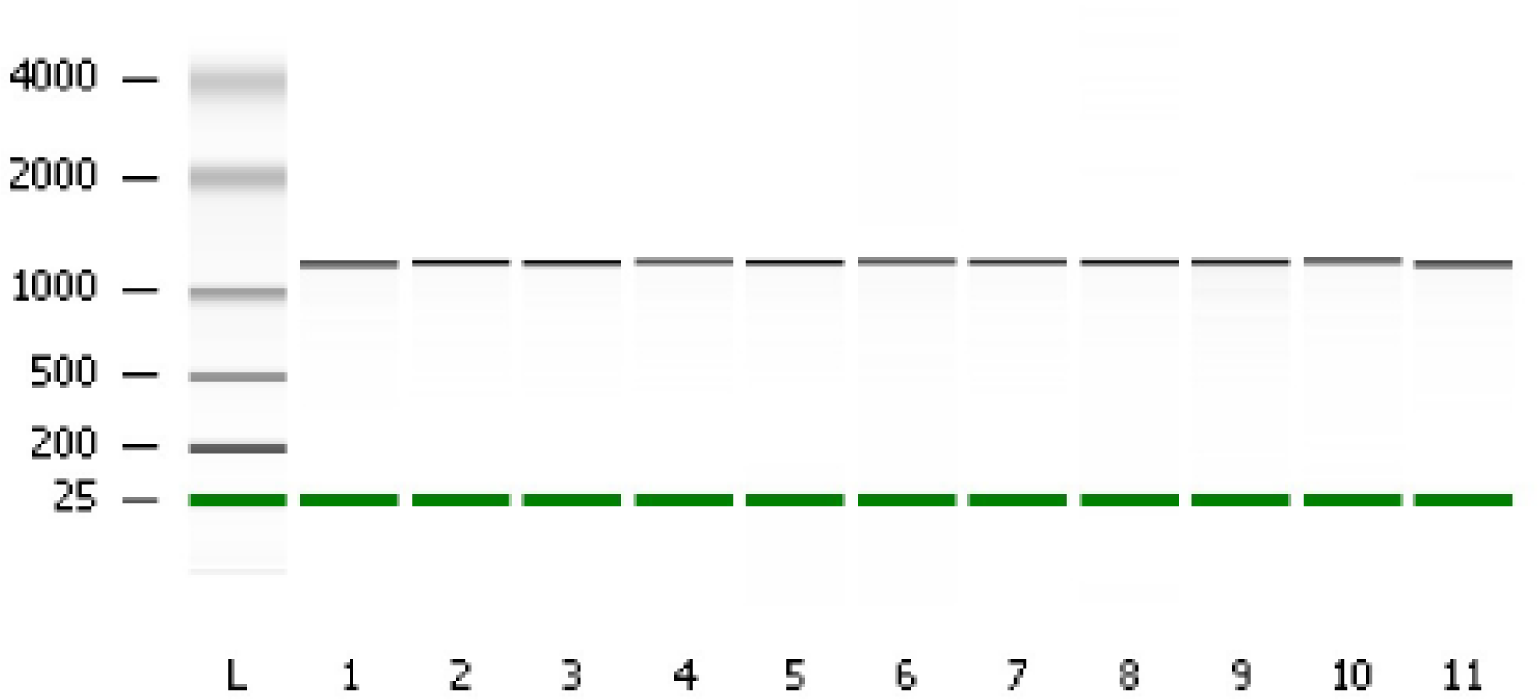
Study Material 4 short term stability: Size analysis using an Agilent 2100 Bioanalyzer. Lane (L): 1-2, Dry ice; 3-4, 4 °C; 5-6, 27 °C; 7-11, −80 °C.

#### Design of long-term stability studies

Long-term stability (LTS) of the Study Materials was monitored over the duration of study participation and changes in copy number concentration evaluated by RT-dPCR. Only units stored at the reference temperature of −80 °C were analysed as this is the storage temperature for RNA samples which was recommended to participants.

Study Material 1 stability was evaluated by NIMC at 0 months, 4 months and 9 months post-production (March, July, December 2020) (*n* = 3 units per timepoint) after preparation, with measurements of both *N* and *E* genes performed (RT-dPCR *n* = 3) (see Appendix B).

Study Material 2, 3 and 4 stability was evaluated by NML for ≈ 6 months (November 2020) after the date of the homogeneity and short-term stability studies (May 2020), with four units of each material being assessed. Triplicate RT-dPCR measurements of each unit were performed with the CDC N2 assay. A single gravimetric dilution series was performed with each unit of Study Material 4 (as described for the short-term stability study of SM4, Appendix B).

#### Results of long-term stability studies

The results of long-term stability studies (LTS) are shown in Figure 3.

**Figure 3:**
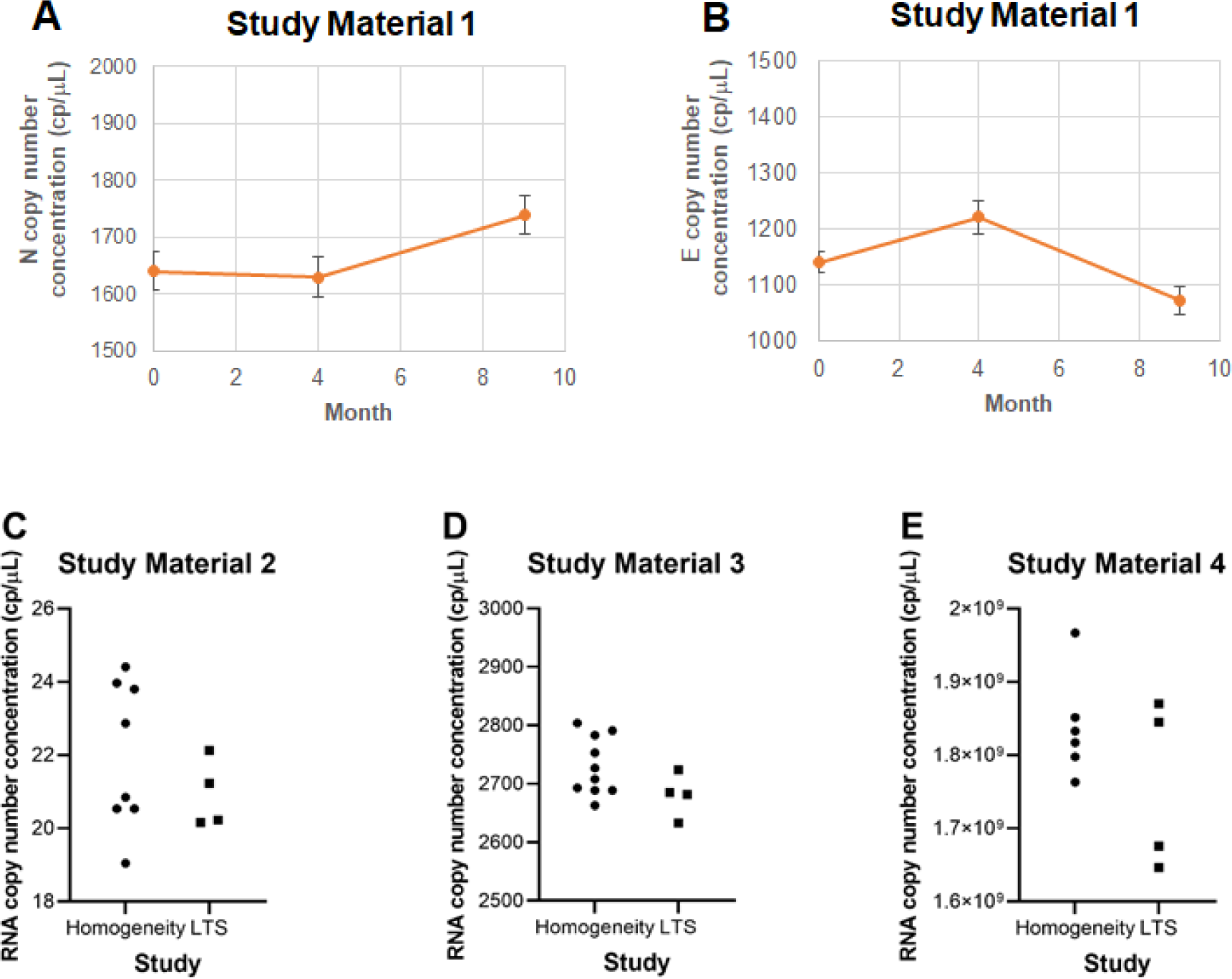
Results of the long-term stability assessment studies. (A) Study Material 1 (Measurand 1) (B) Study Material 1 (Measurand 2); Error bars show between-unit SD. Measurand 1 results for (C) Study Material 2; (D) Study Material 3 and (E) Study Material 4. Dots represent mean value per unit.

LTS results for Study Material 1 were analyzed with linear regressions by plotting time points (*x*, months) and RNA copy concentration (*y*, cp/μL). The slope of the regression lines was tested for statistical significance (95 % confidence level). No obvious trend was observed for both E and N gene after storage of 9 months and the observed values were within the range of the coordinator’s assigned values (Table 5).

**Table 5:** Coordinator’s assigned values and uncertainties (Study Material 1).

| MATERIAL | MEASURAND (UNIT) |
| --- | --- |
| **STUDY MATERIAL 1** | **Measurand 1 (N gene) (/μL)** |
| Value (x) | 1640 |
| Standard uncertainty (u) | 57 |
| Coverage factor (k) | 2 |
| Expanded uncertainty (U) | 113 |
| Relative expanded uncertainty (Rel U) | 7 % |
|  | **Measurand 2 (E gene) (/μL)** |
| Value (x) | 1142 |
| Standard uncertainty (u) | 76 |
| Coverage factor (k) | 2 |
| Expanded uncertainty (U) | 152 |
| Relative expanded uncertainty (Rel U) | 13 % |

LTS results for Study Materials 2, 3 and 4 were analyzed using the model:

*y* ≈ Time / Unit,
where Unit (a random effect) is nested within time point (a fixed effect), and *y* is either λ (Study Materials 2 and 3) or RNA copy concentration (Study Material 4). A between-time point mean square *MS_t_* which is larger than the between-unit mean square *MS_u_* indicates a non-zero time effect (which may or may not be significant). For Study Material 2, *MS_t_* < *MS_u_*, and this shows that there is no significant evidence for a difference in mean between the two time points. For Study Materials 3 and 4 *MS_t_* > *MS_u_*, therefore results were then analyzed using a mixed-effects model with restricted maximum likelihood, with timepoint as fixed factor and unit as random factor. This confirmed that there was no significant effect of storage for either Study Material 3 or 4 (*p* = 0.20 and *p* = 0.21 respectively). The standard error associated with time in the mixed-effects model was included as an allowance for stability in the coordinator’s value assignment of these materials.

### Coordinators’ value assignment of Study Materials

#### Study Material 1

The coordinator’s assigned values for Study Material 1 are shown in Table 5 with contributions to the uncertainty in the assigned values shown in Table 6. Uncertainty due to long term stability was calculated according to ISO Guide 35.

**Table 6:** Uncertainty contributions to coordinator’s assigned values (Study Material 1).

| Factor | Measurand 1 (N) | Measurand 2 (E) |
| --- | --- | --- |
| Method precision (%) | 1.77 | 1.84 |
| Homogeneity (%) | 0.85 | 0.88 |
| Partition volume (%) | 0.80 | 0.80 |
| Stability (%) | 2.7 | 6.3 |

Contributions are shown as relative standard uncertainties.

#### Study Materials 2, 3 and 4

The coordinator’s assigned values for Measurand 1 in Study Material 2, 3 and 4 are shown in Table 7 based on measurements performed with the US CDC N2 gene assay. Measurand 2 was not measured in Study Material 2 as it was expected that the ratio of the *N* and *E* genes was approximately equal. Contributions to the uncertainty in the assigned values are shown in Table 8. For Study Material 2, no difference in variation between units was observed in the long-term stability study compared to the homogeneity study, therefore no allowance was included for stability. For Study Materials 3 and 4, an increase in variance was associated with time in the long-term stability study, therefore an allowance is included based on the LTS (Table 8). Method precision is based on within-unit variation in the homogeneity study (repeatability conditions).

**Table 7:**
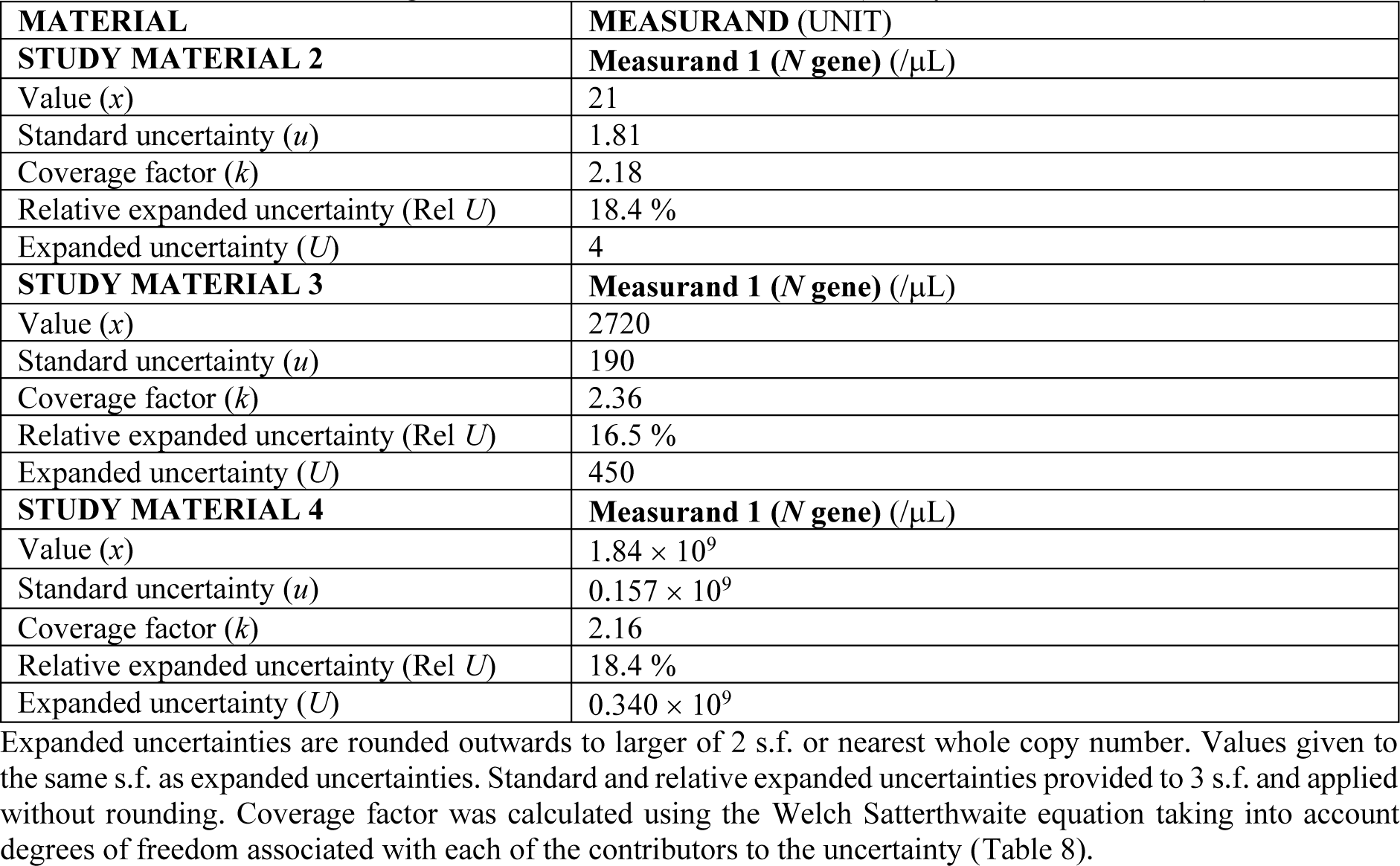
Coordinator’s assigned values and uncertainties (Study Materials 2, 3, 4)

**Table 8:** Uncertainty contributions to coordinator’s assigned values (Study Materials 2, 3, 4)

| **Factor** | **Study Material 2** | **Study Material 3** | **Study Material 4** |
| --- | --- | --- | --- |
| Method precision (%) | 4.63 % | 0.840 % | 0.650 % |
| Homogeneity (%) | 2.68 % | 1.94 % | 4.43 % |
| Partition volume (%) | 5.19 % | 5.19 % | 5.19 % |
| Stability (%) | No allowance | 1.19 % | 3.12 % |
| Assay (%) | 4.00 % | 4.00 % | 4.00 % |

Contributions are shown as relative standard uncertainties to 3 s.f.

## SAMPLE DISTRIBUTION

The majority of Study Materials were shipped on dry ice, however due to import or freight carrier restrictions, some participants received materials shipped on ice packs. Laboratory 12 did not receive Study Material 1 as it was not possible to arrange direct shipment from China.

## TIMELINE

Table 11 shows the timeline for CCQM-P199b.

**Table 9:**
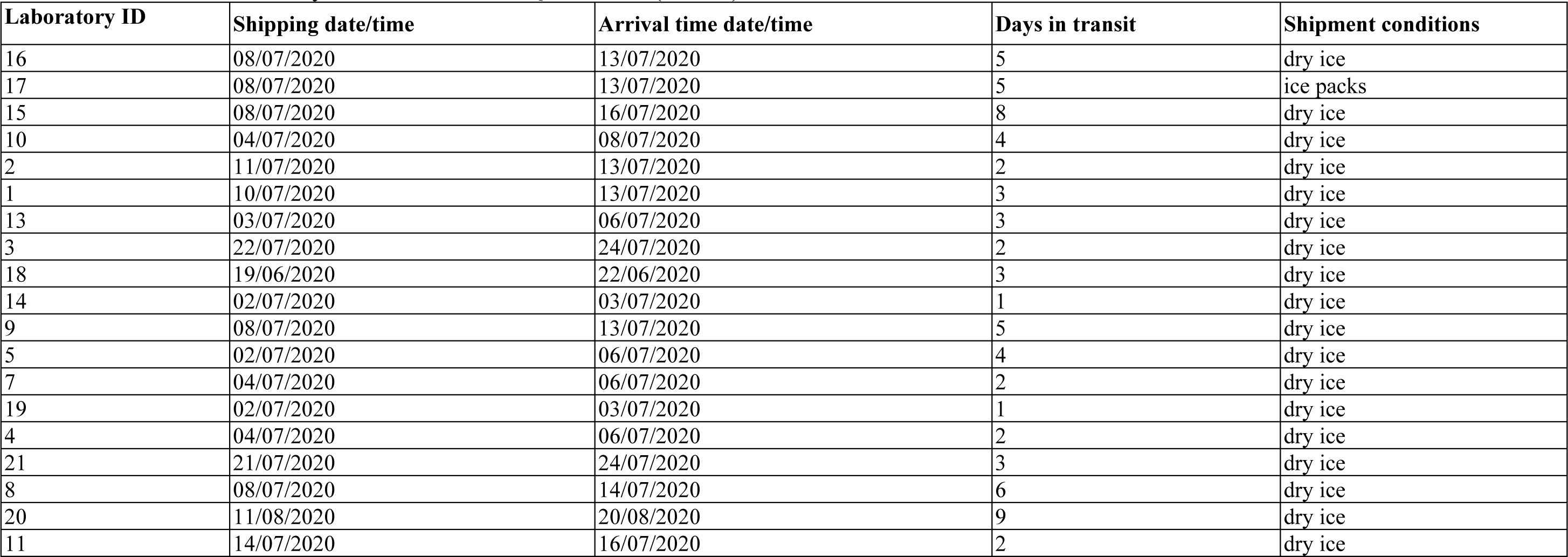
Distribution of Study Material 1 for CCQM-P199b (NIMC).

**Table 10:** Distribution of Study Materials 2-4 for CCQM-P199b (NML).

| Laboratory ID | Study Material 4 | Shipped Date (UTC) | Arrival Date (UTC) | Days in transit | Shipment conditions | Comments |
| --- | --- | --- | --- | --- | --- | --- |
| 16 |  | 19/06/2020, 17:51 | 02/07/2020 | 13 | dry ice |  |
| 17 |  | 12/06/2020, 9:32 | 15/06/2020, 17:44 | 3.3 | dry ice |  |
| 10 |  | 02/06/2020, 15:22 | 03/06/2020, 9:00 | 0.7 | dry ice |  |
| 12 |  | 17/07/2020 | 05/08/2020 | 19 | dry ice | Arrival time not recorded |
| 15 | Y | 29/06/2020 | 01/07/2020, 13:00 | 2 | dry ice |  |
| 2 |  | 02/06/2020, 15:22 | 03/06/2020, 12:33 | 0.8 | dry ice |  |
| 1 |  | 09/06/2020, 14:33 | 10/06/2020, 13:00 | 0.8 | dry ice |  |
| 13 | Y | 19/06/2020 | 22/06/2020, 8:10 | 3 | dry ice |  |
| 18 |  | 02/06/2020, 15:22 | 03/06/2020, 10:15 | 0.8 | dry ice |  |
| 3 |  | 02/06/2020, 15:22 | 03/06/2020, 10:10 | 0.8 | dry ice |  |
| 14 |  | 12/06/2020 | 17/06/2020 | 5 | dry ice | Arrival time not recorded |
| 6 | Y | 03/07/2020 | 07/07/2020 | 4 | dry ice | Form 2 not received |
| 9 |  | 12/06/2020, 09:32 | 15/06/2020, 16:40 | 3.2 | dry ice |  |
| 5 | Y | 12/06/2020 | 15/06/2020 | 3 | dry ice | Form 2 not received |
| 7 | Y | 19/06/2020 | 23/06/2020 | 4 | dry ice | Arrival time not recorded |
| 4 | Y | 02/06/2020, 15:22 | 03/06/2020, 10:00 | 0.8 | dry ice |  |
| 21 |  | 29/06/2020, 9:04 | 09/07/2020, 16:25 | 10.3 | ice packs |  |
| 8 | Y | 01/06/2020, 9:31 | 05/06/2020, 12:49 | 4.1 | dry ice |  |
| 20 |  | 29/06/2020, 9:04 | 03/07/2020, 15:45 | 5.2 | ice packs |  |
| 11 |  | 29/06/2020, 9:04 | 07/07/2020, 10:00 | 8 | ice packs |  |

**Table 11:** Timeline for CCQM-P199b.

| Date | Action |
| --- | --- |
| 05 May 2020 | Call for participation (Protocol v1.0 circulated) |
| 06 May 2020 | Study proposal and authorization at NAWG meeting |
| 12 May 2020 | Registration deadline for participants |
| 22 May to 11 August 2020 | Study Material 1 distribution (NIMC) |
| 01 June to 31 July 2020 | Study Material 2 to 4 distribution (NML) |
| 05 August 2020 | Protocol v1.1 circulated (Results of homogeneity and stability studies) |
| 30 September 2020 | Result submission deadline (participants informed 05 August 2020) |
| 14 October 2020 | Extension to Result submission deadline (participants informed 14 September 2020) |
| 20 November 2020 | Anonymised Study Results sent to participants for review |
| 24-25 November 2020 | Presentation of Study Results at CCQM NAWG meeting (Study Material 4 results blinded) |
| 10 March 2021 | Draft A Report circulated to Participants |
| 30 March 2022 | Draft B Report circulated to Participants |
| November 2022 | Final Report circulated to NAWG |

## RESULTS

### Reporting of Results

Participants were requested to report an average value and expanded uncertainty for RNA copy number concentration result for each of the Measurands and Materials.

In addition to the quantitative results (Form 3), participants were instructed to describe their analytical methods, experimental design and approach to uncertainty estimation (Form 4). As the majority of laboratories used RT-dPCR, dPCR experimental parameters were recorded in Form 5 based on the Minimum Information for Publication of Digital PCR Experiments (dMIQE) guidelines [19].

CCQM-P199b results were submitted by 21 of the 22 institutions that received samples. Laboratory 8 withdrew their result for Study Material 2 due to inconsistencies between the values for the two measurands (which were expected to be approximately equal based on both genes being on the same lentiviral construct, as described in Study Protocol). Laboratory 12 did not receive Study Material 1. Seven laboratories received Study Material 4 which was intended for analysis with orthogonal (non RT-dPCR) methods; Laboratory 4 and Laboratory 8 did not report a result for Study Material 4.

Laboratory 10 resubmitted their results following unblinding of the Study Results (19 November 2020) due to noticing that a dilution factor had not correctly been applied. This was before discussion of the results at CCQM-Nucleic Acid Working Group (NAWG) meeting, therefore both original and amended results were presented by the coordinator (Appendix F).

Following discussion of the study results (January 2021), Laboratory 8 provided supplementary follow-up data for Study Materials 1 to 4 using one-step RT-dPCR. Laboratory 20 also provided supplementary data from two-step RT-dPCR analysis which was performed at the time of study participation (Appendix K).

### Calibration Materials Used by Participants

For the analysis of Study Materials 1 to 3, participants did not apply calibration materials, with the exception of Laboratory 21 which used RT-qPCR calibrated to commercial RM (Table 12).

**Table 12:**
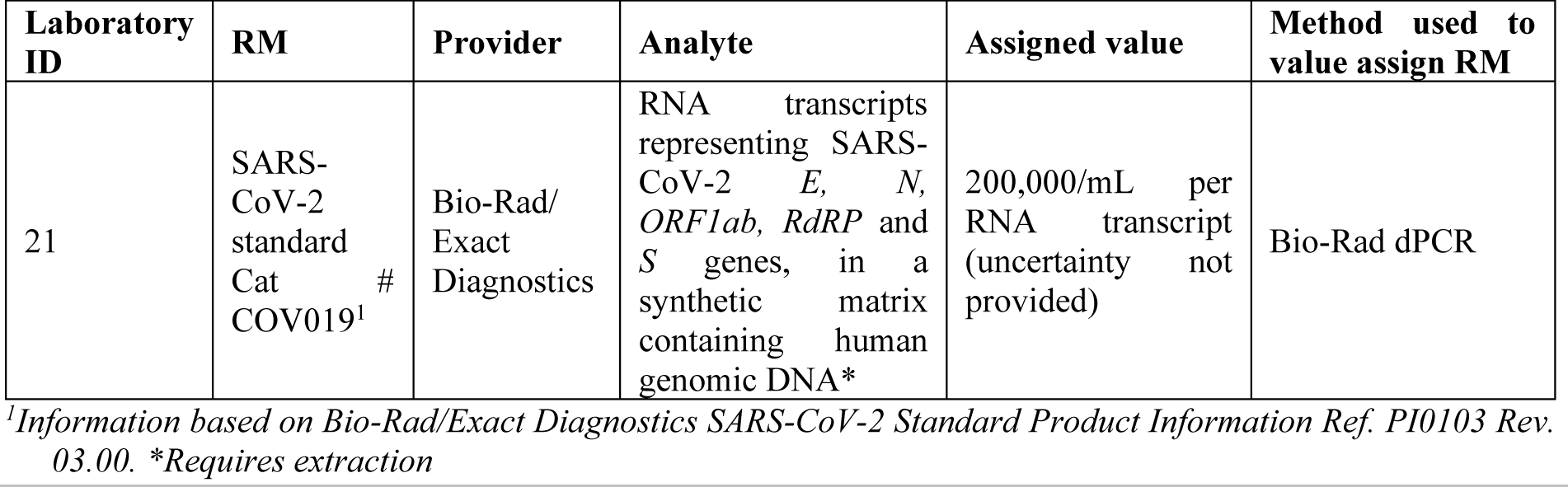
Calibration Materials used for measurement of Study Material 1-3.

For the analysis of Study Material 4, laboratories performing ID-MS applied in-house calibration materials which were characterised in terms of purity. Table 13 lists the materials, their assigned molar concentration or purity, the method used, and how the participant had demonstrated their competence in the use of the method(s).

**Table 13:** Calibration Materials used for measurement of Study Material 4.

| Laboratory ID | CRM | Provider | Measurand | Certified value and uncertainty<sup>a</sup> [units] | Method used to value assign CRM; basis for (traceability) |
| --- | --- | --- | --- | --- | --- |
| 5 | Calibration solutions prepared and certified at Laboratory 5 | Lab. 5 | Amount of substance concentration of nucleotide monophosphate solutions | AMP-1-B<br>751.4 nmol/g<br>±1.4 nmol/g<sup>b</sup><br>UMP-1-B<br>601.9 nmol/g<sup>b</sup><br>GMP-1-B<br>586.5 nmol/g<sup>b</sup><br>CMP-1-B<br>620.5 nmol/g<sup>b</sup> | Quantitative NMR (traceable to NMIA CRM QNMR002) |
| 6 | GBW(E) 100154 Purity of Adenosine-5' monophosphoric acid | NIMC | Purity (mass fraction) of nucleotide monophosphoric acid, | AMP: 98.9 % ± 0.7 % (g/g)<br>CMP: 99.3 % ± 0.6 % (g/g) | HPLC and HR-ICP-MS |
| | GBW(E) 100067<br>Purity of Cytidine-5'monophosphoric acid<br>GBW(E) 100068<br>Purity of Guanosine-5'monophosphate disodium salt<br>GBW(E) 100069<br>Purity of Uridine-5'monophosphate disodium salt | | Purity (mass fraction) of nucleotide monophosphate disodium salt | GMP: 98.8 %± 0.6 % (g/g)<br>UMP: 99.4 %± 0.4 % (g/g) | |
| 7 | In-house NMP standard solutions | Lab. 7 | Amount of substance concentration of nucleotide monophosphate | AMP: 1083.5 nmol/g±31.4 nmol/g<br>CMP: 1179.2 nmol/g±31.9 nmol/g<br>GMP: 1052.8 nmol/g±24.2 nmol/g<br>UMP: 1155.7 nmol/g±25.7 nmol/g | <sup>1</sup>H NMR using potassium hydrogen phthalate (traceable to NMIJ CRM 3001-b as internal standard). |

*a* Stated as Value ± $U_{95}$ (Value)
*b* The certified value expanded uncertainty used $k = 2$

### Methods Used by Participants

For the analysis of Study Materials 1, 2 and 3, the majority of laboratories (19/21) used RT-droplet dPCR (Bio-Rad QX100/QX200 systems), with 17 of those laboratories using the One-Step RT-ddPCR Advanced Kit for Probes (Bio-Rad) and two laboratories employing a two-step RT-dPCR approach (Laboratory 8 and Laboratory 11). One laboratory (Laboratory 7) used the microfluidic chip-based QS3D dPCR system (Thermo Fisher Scientific). One laboratory (Laboratory 21) used one-step RT-qPCR, calibrated using a commercial SARS-CoV-2 standard (Table 12 above).

For the analysis of Study Material 4, five laboratories reported results. Laboratory 5 and Laboratory 6 each reported two results sets using RT-dPCR and ID-MS. Laboratory 7 measured Study Material 4 using ID-MS. Laboratory 13 measured this material using SMFC with specific oligonucleotide probes. Laboratory 15 was not able to measure Study Material 4 using orthogonal methods therefore submitted a result using RT-dPCR.

Further information on the analytical techniques and dPCR methodological parameters are summarized in Appendix H. The participants’ approaches to estimating uncertainty are provided in Appendix I.

### Participant Results

Participant results for CCQM-P199b are detailed in Tables 14 to 17 and presented graphically in Figures 4 to 7.

**Figure 4:**
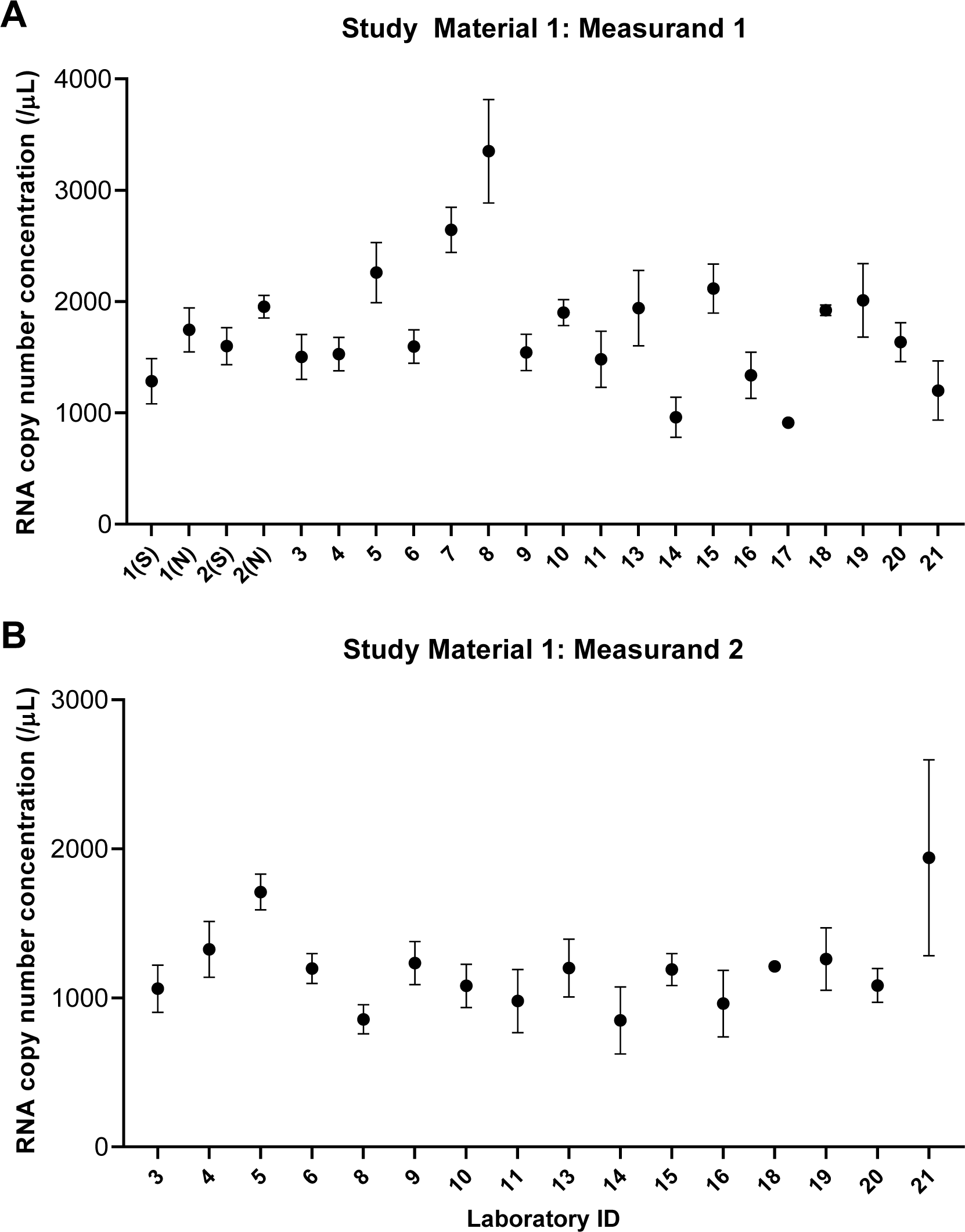
CCQM-P199b participants’ measurement results for Study Material 1. Dots represent the reported values, *x*; bars their 95 % expanded uncertainties, *U*(*x*). N, nominated result. S, supplementary result.

**Figure 5:**
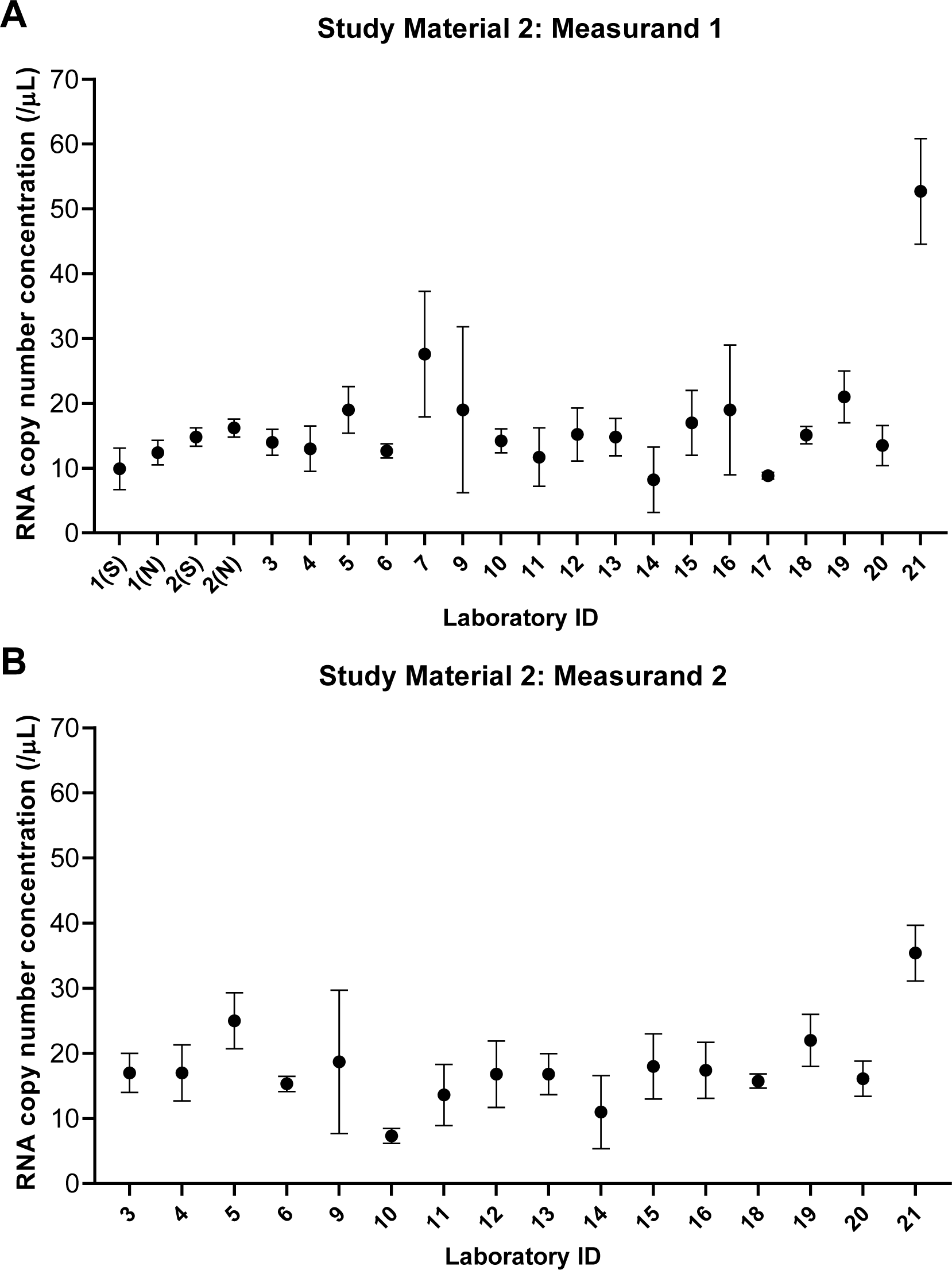
CCQM-P199b participants’ measurement results for Study Material 2. Dots represent the reported values, *x*; bars their 95 % expanded uncertainties, *U*(*x*). N, nominated result. S, supplementary result.

**Figure 6:**
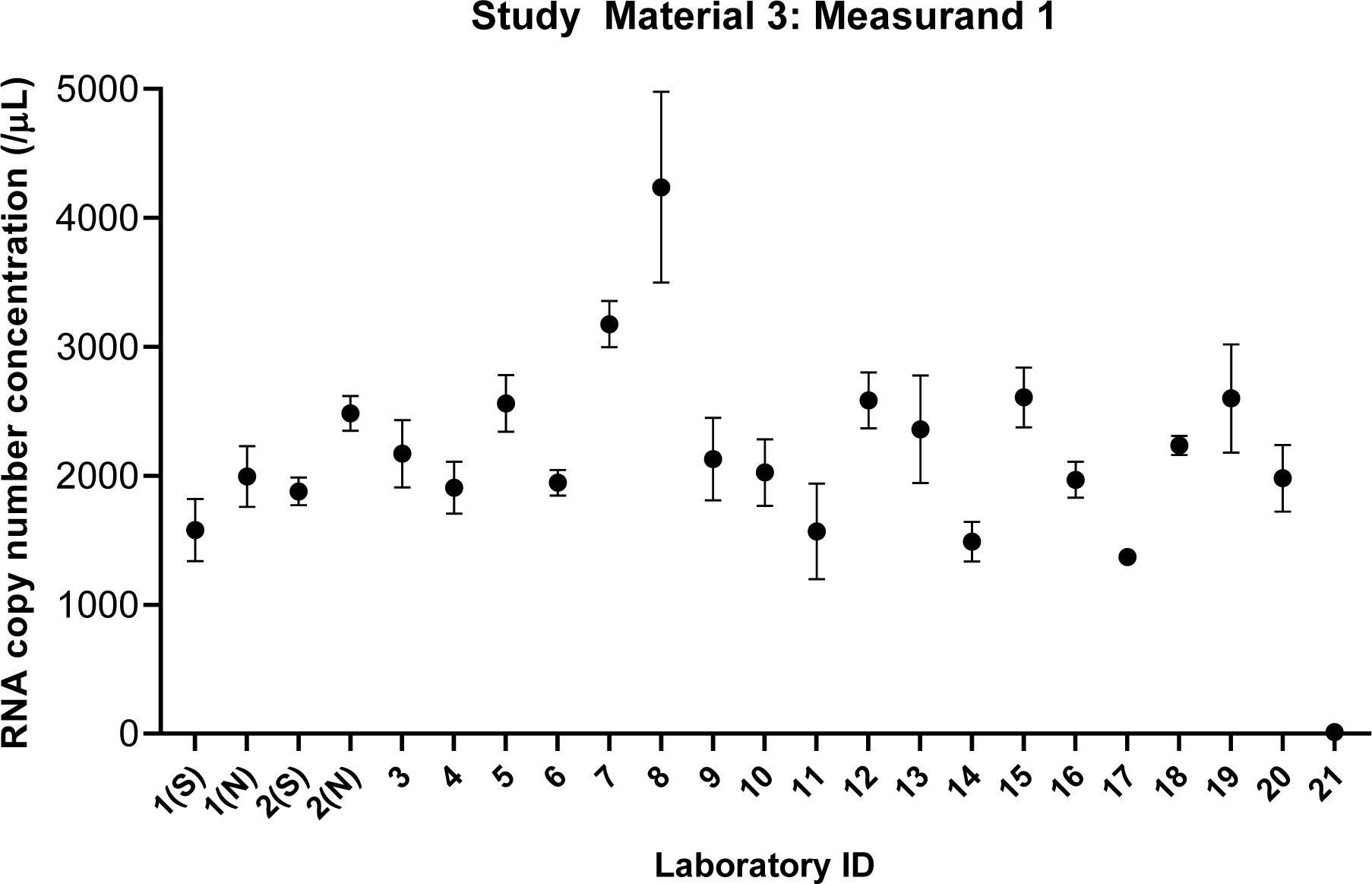
CCQM-P199b participants’ measurement results for Study Material 3. Legend as Figures 4-5.

**Figure 7:**
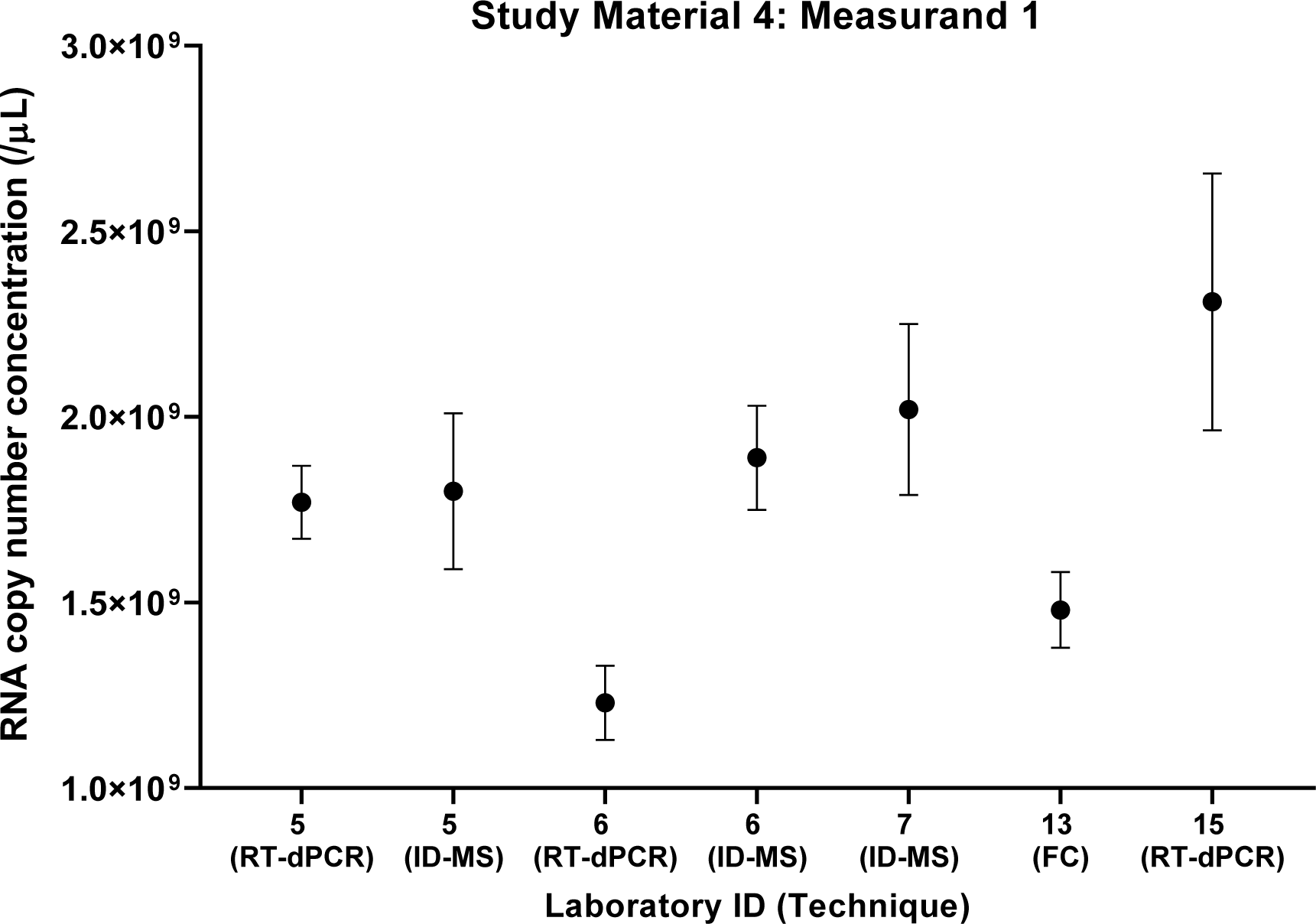
CCQM-P199b participants’ measurement results for Study Material 4. Dots represent the reported values, *x*; bars their 95 % expanded uncertainties, *U*(*x*). Techniques used by participants are given in brackets: ID-MS, isotope dilution mass spectrometry; FC, single molecule flow cytometry.

**Table 14:**
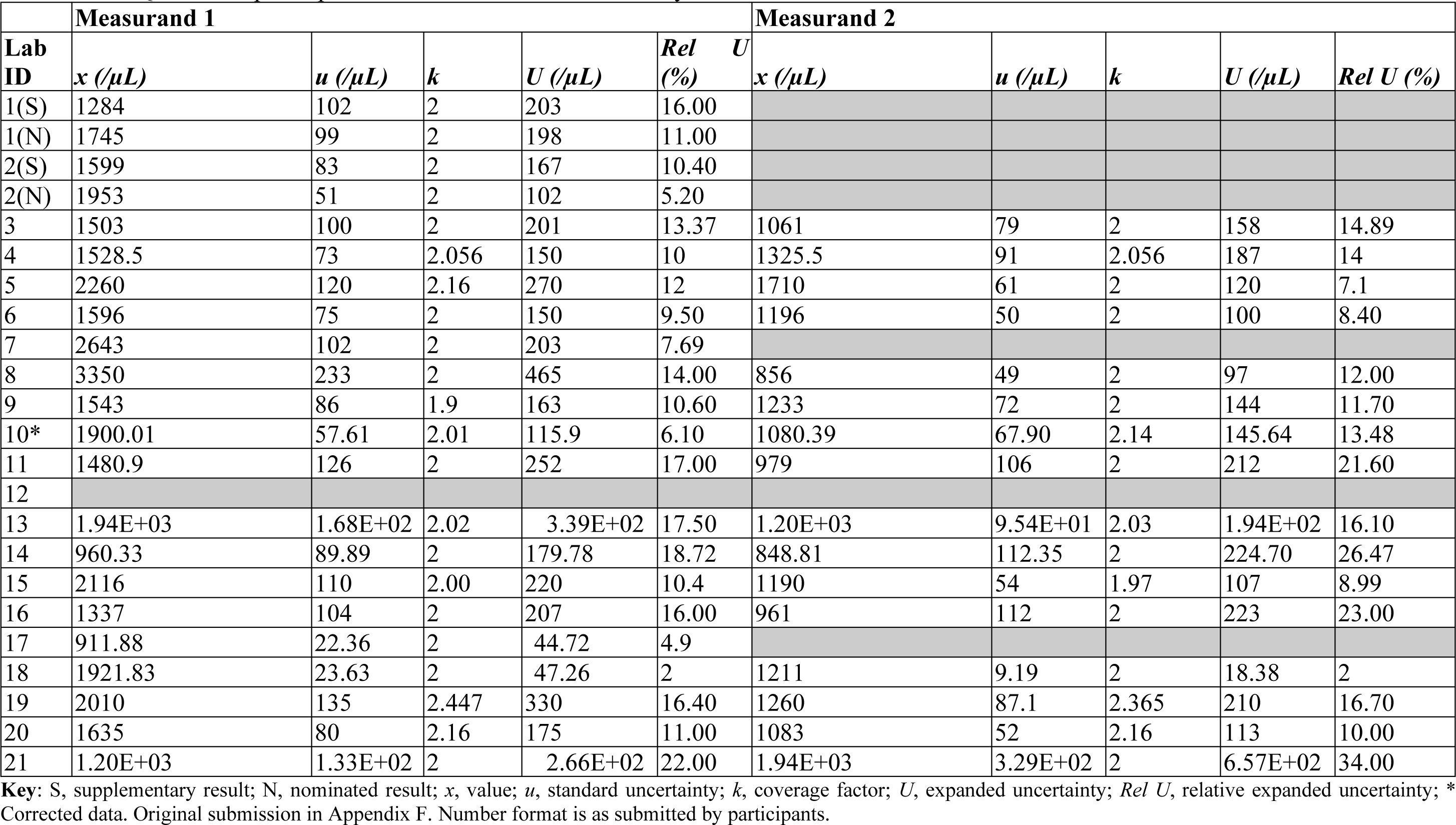
CCQM-P199b participants’ measurement results for Study Material 1.

**Table 15:**
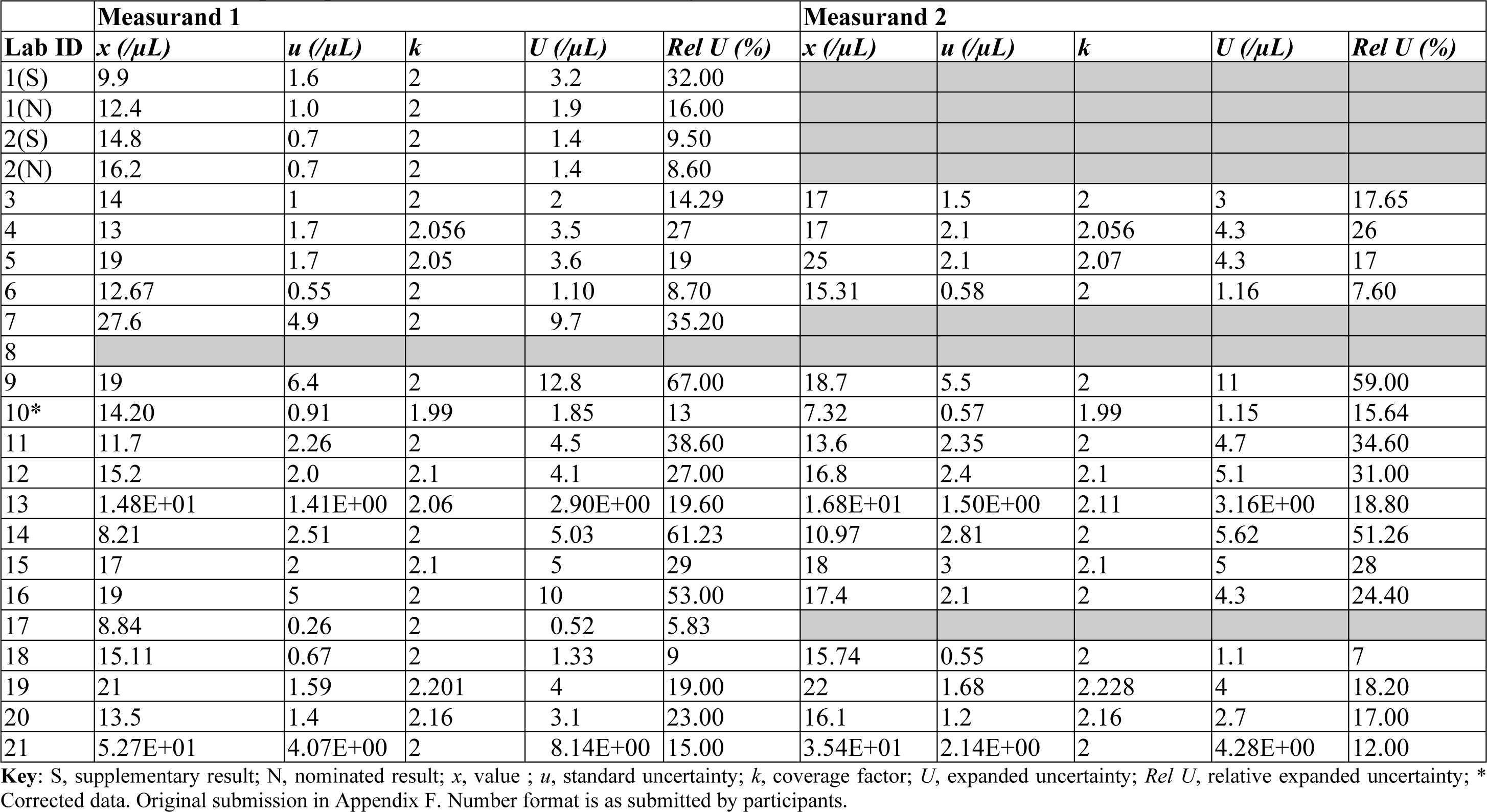
CCQM-P199b participants’ measurement results for Study Material 2.

**Table 16:**
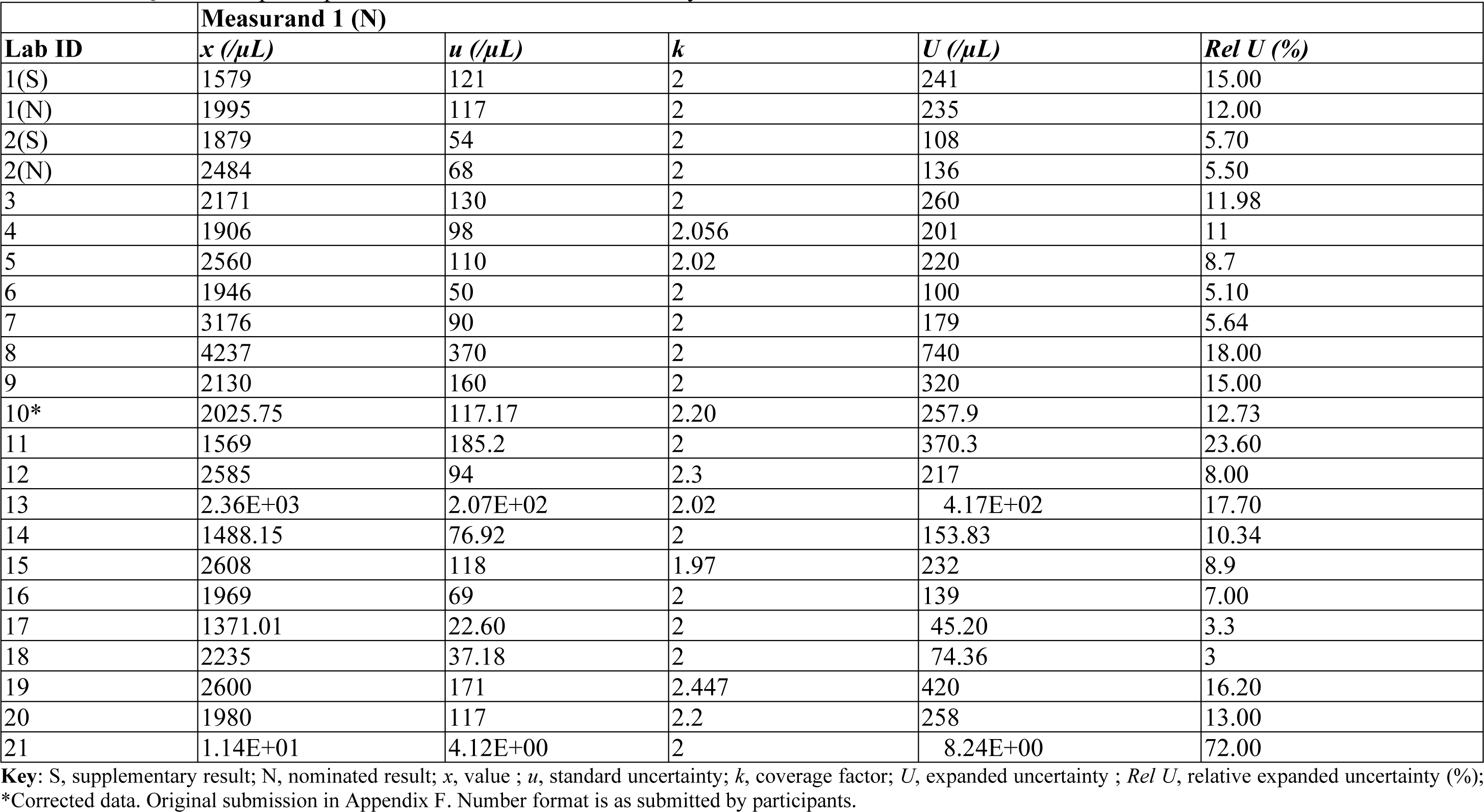
CCQM-P199b participants’ measurement results for Study Material 3.

**Table 17:** CCQM-P199b participants’ measurement results for Study Material 4.

| Lab ID | Technique | Measurand 1 |  |  |  |  |
| --- | --- | --- | --- | --- | --- | --- |
| | | $x$ (/μL) | $u$ (/μL) | $k$ | $U$ (/μL) | $Rel\ U$ (%) |
| 1 |  |  |  |  |  |  |
| 2 |  |  |  |  |  |  |
| 3 |  |  |  |  |  |  |
| 4 |  |  |  |  |  |  |
| 5A | RT-dPCR | 1.77E+09 | 4.8E+07 | 2.03 | 9.8E+07 | 5.5 |
| 5B | ID-MS | 1.80E+09 | 9.3E+07 | 2.2 | 2.1E+08 | 11 |
| 6A | RT-dPCR | 1.23E+09 | 5E+07 | 2 | 1.0E+08 | 8.40 |
| 6B | ID-MS | 1.89E+09 | 7E+07 | 2 | 1.4E+08 | 7.4 |
| 7 | ID-MS | 2.02E+09 | 1.15E+08 | 2 | 2.30E+08 | 11.4 |
| 8 |  |  |  |  |  |  |
| 9 |  |  |  |  |  |  |
| 10 |  |  |  |  |  |  |
| 11 |  |  |  |  |  |  |
| 12 |  |  |  |  |  |  |
| 13 | SMFC | 1.48E+09 | 4.40E+07 | 2.3 | 1.02E+08 | 6.90 |
| 14 |  |  |  |  |  |  |
| 15 | RT-dPCR | 2.31E+09 | 1.35E+08 | 2.57 | 3.46E+08 | 15 |
| 16 |  |  |  |  |  |  |
| 17 |  |  |  |  |  |  |
| 18 |  |  |  |  |  |  |
| 19 |  |  |  |  |  |  |
| 20 |  |  |  |  |  |  |
| 21 |  |  |  |  |  |  |

**Key:** For laboratories submitting > 1 result: A, Result 1; B, Result 2; $x$ , value (/μL); $u$ , standard uncertainty (/μL); $k$ , coverage factor; $U$ , expanded uncertainty (/μL); $Rel\ U$ , relative expanded uncertainty. Number format is as submitted by participants.

All 21 laboratories measured Study Material 3 (Measurand 1). Twenty laboratories reported values for Measurand 1 in Study Materials 1 and 2. Sixteen laboratories reported values for optional Measurand 2 in Study Materials 1 and 2. Two laboratories (Laboratories 1 and 2) reported two results for Study Materials 1, 2 and 3 (Measurand 1), with one result being nominated as their main result and one result being designated as supplementary. Five laboratories reported results for Study Material 4, with two laboratories (Laboratories 5 and 6) reporting two sets of results (both treated as nominated results as completely different methods were used).

Laboratory 21 reported a low value for Study Material 3 (Measurand 1) which was attributable to the assay used not being able to amplify the Study Material 3 template. Laboratory 21’s RT-qPCR assay for the N gene used was based on the Hong Kong University developed assay (amplicon position reference NC_045512.2:29145-29254), with the reverse primer being situated outside of the template region (Table 1).

### Interlaboratory reproducibility and consistency

Table 18 provides a summary of the reproducibility of the study results according to Study Material, Measurand and technique expressed as the adjusted median absolute deviation from the median (MADe) or standard deviation (SD) or their values relative to the average (median or mean). For Study Materials 1 and 2, all nominated results are shown as well as only data for laboratories performing RT-dPCR. For Study Material 3, the RT-qPCR result (laboratory ID 21) was excluded due to the *N* gene assay used not being able to detect the Study Material 3 partial *N* gene construct. For Study Materials 1 and 2, the reproducibility was better for Measurand 2 (*E* gene) compared to Measurand 1 (*N* gene). This is likely to be associated with a wider variety of assays and techniques being used for Measurand 1 compared to Measurand 2. The reproducibility of ID-MS results for the measurement of Study Material 4 was good, with an interlaboratory % CV of 5.8 % compared to RT-dPCR, which varied between 19 % and 31 % (range for all Study Materials and Measurands).

**Table 18:** Summary of descriptive statistics for nominated results.

| Study Material (Measurand) Data | Median (/μL) | MADe (/μL) | Relative MADe | Normal distrib. (Y/N)** | Mean (/μL) | SD (/μL) | % CV |
| --- | --- | --- | --- | --- | --- | --- | --- |
| 1 (1) all | 1690 | 380 | 23 % | Y | 1777 | 562 | 32 % |
| 1 (1) RT-dPCR | 1745 | 359 | 21 % | Y | 1807 | 560 | 31 % |
| 1 (2) all | 1193 | 181 | 15 % | N | 1196 | 286 | 24 % |
| 1 (2) RT-dPCR | 1190 | 163 | 14 % | Y | 1146 | 213 | 19 % |
| 2 (1) all | 15.0 | 3.6 | 24 % | N | 17.3 | 9.4 | 55 % |
| 2 (1) RT-dPCR | 14.8 | 3.3 | 22 % | Y | 15.4 | 4.5 | 29 % |
| 2 (2) all | 16.9 | 2.0 | 12 % | N | 17.7 | 6.2 | 35 % |
| 2 (2) RT-dPCR | 16.8 | 1.8 | 11 % | Y | 16.5 | 4.1 | 25 % |
| 3 (1) RT-dPCR | 2150 | 429 | 20 % | N | 2270 | 634 | 28 % |
| 4 (1) all | 1.800 ×10<sup>9</sup> | 3.26 ×10<sup>8</sup> | 18 % | Y | 1.786 ×10<sup>9</sup> | 3.52×10<sup>8</sup> | 20 % |
| 4 (1) RT-dPCR | *n too small* |  |  | N/A | 1.770 ×10<sup>9</sup> | 5.40×10<sup>8</sup> | 31 % |
| 4 (1) ID-MS | *n too small* |  |  | N/A | 1.903 ×10<sup>9</sup> | 1.11×10<sup>8</sup> | 5.8 % |

MADe and SD are given to 3 significant figures (s.f.) for Materials 1, 3 and 4, with median or mean to the same order of magnitude.
For Study Material 2, MADe and SD are given to 2 s.f. and median or mean to the same number of decimal places. Relative variation (% MADe or % CV) are given to 2 s.f. for all Materials/Measurands.
\*\*Normality testing shows the result of Shapiro-Wilk test (alpha = 0.05).
For non-normally distributed datasets, mean, SD and % CV shown for information only (grey).
N/A: normality not assessed for subsets of Study Material 4 data as number of laboratories too small (*n* = 3).

The results of pairwise comparison of individual laboratories’ results and uncertainties was performed (Appendix J). The majority of results were consistent between laboratories. In some cases, underestimation in the reported uncertainties (for example where only precision terms where included in uncertainty budgets) led to statistically significant differences in values with other laboratories. As observed in the reproducibility metrics, Measurand 2 for Study Material 1 was measured more consistently between laboratories than Measurand 1. Study Material 2 results (both Measurands) showed fewer inconsistencies between laboratories, presumably due to the low concentration of the material leading to less precise measurements and higher reported laboratory uncertainties. The Study Material 3 dataset showed outlying results at both ends of the reported concentration range (Laboratories 14 and 17 at the lower end and Laboratories 7 and 8 at the higher end of the range). In the Study Material 4 dataset, the Laboratory 6 RT-dPCR and Laboratory 13 SMFC results were significantly lower than the other five results (*p* < 0.001 for majority of pairwise comparisons), whilst the Laboratory 15 RT-dPCR was higher than the majority of results (*p* < 0.05), with the exception of the Laboratory 7 ID-MS result. Factors which may have led to differences between participants’ results are explored further in the Discussion of Results section.

### Comparison between Study Material 3 and 4 results

As Study Material 3 was prepared by gravimetric dilution of Study Material 4, the results from analysis of the two materials were compared to inform sources of bias between the techniques used for analysis (predominantly RT-dPCR for Study Material 3, and ID-MS, SMFC and RT-dPCR for Study Material 4). The comparison also enables evaluation of the ability of RT-dPCR to analyse highly concentrated materials which require serial dilution. The results of the five laboratories who measured Study Material 4 were extrapolated to the Study Material 3 concentration range (Table 19) by division by the dilution factor of 8.087 × 10^5^ linking the two study materials (Table 2). In addition, the submitted standard measurement uncertainties were combined with additional uncertainties for Study Material 4 homogeneity (relative *u* = 2.2 % for the analysis of 4 units) and the dilution uncertainty (relative *u* = 1.40 %, Table 2). The average values for Study Material 4 extrapolated results was very similar to the median of the Study Material 3 results (Table 18).

**Table 19:**
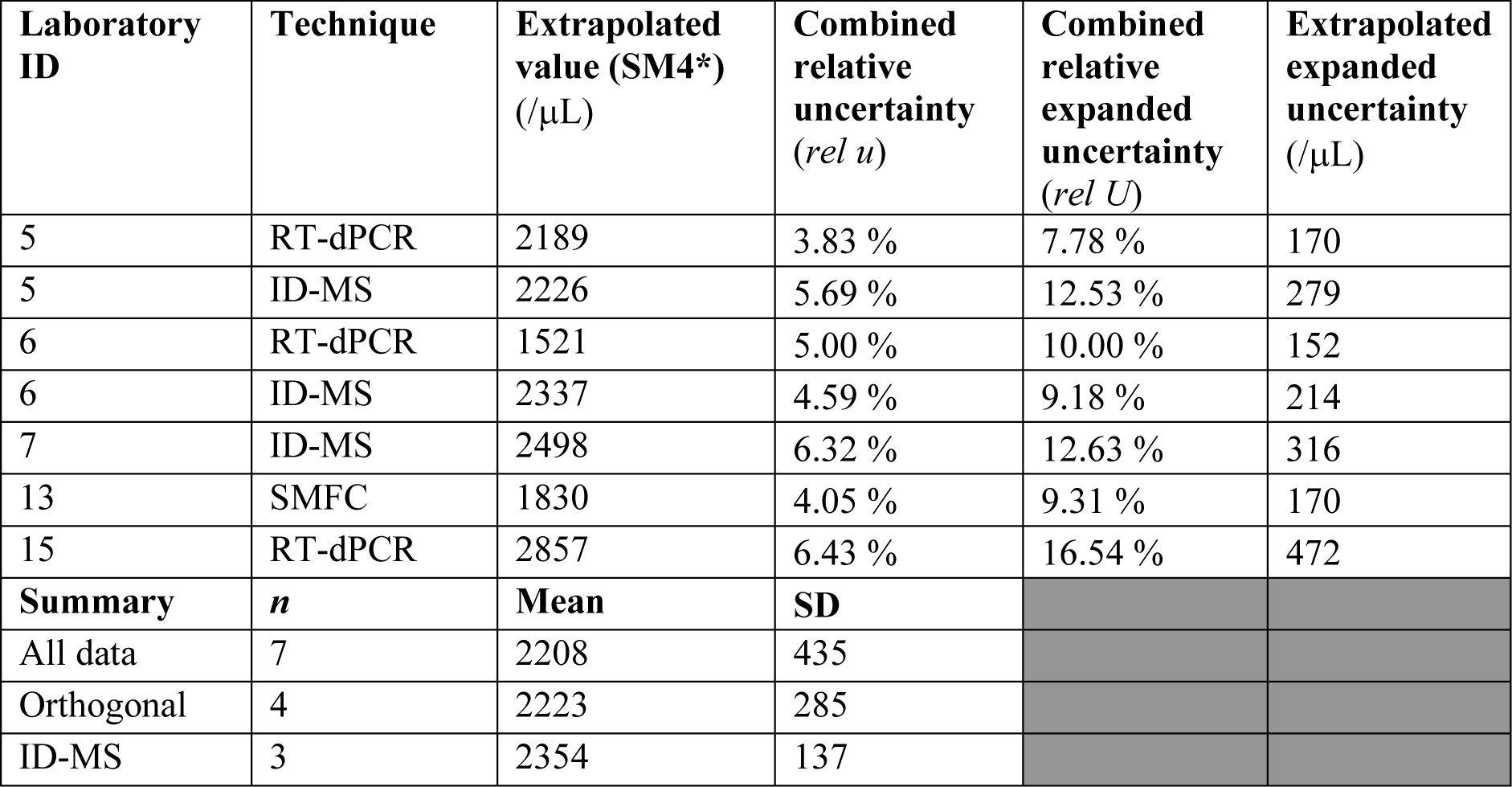
Study Material 4 participant results extrapolated to Study Material 3 concentration range.

Comparison of nominated results for Study Material 3 (*n* = 20) and all extrapolated Study Material 4 (*n* = 7) (Figure 8A) by Welch’s *t*-test or non-parametric Wilcoxon rank sum test show no significant difference in mean values (*p* = 0.78 and *p* = 0.98 respectively). The results of the five laboratories who analysed both study materials are compared in Figure 8B. Generally the results within laboratory were consistent with each other however Laboratory 6’s RT-dPCR-based result for Study Material 4 was lower than the equivalent for Study Material 3.

**Figure 8:**
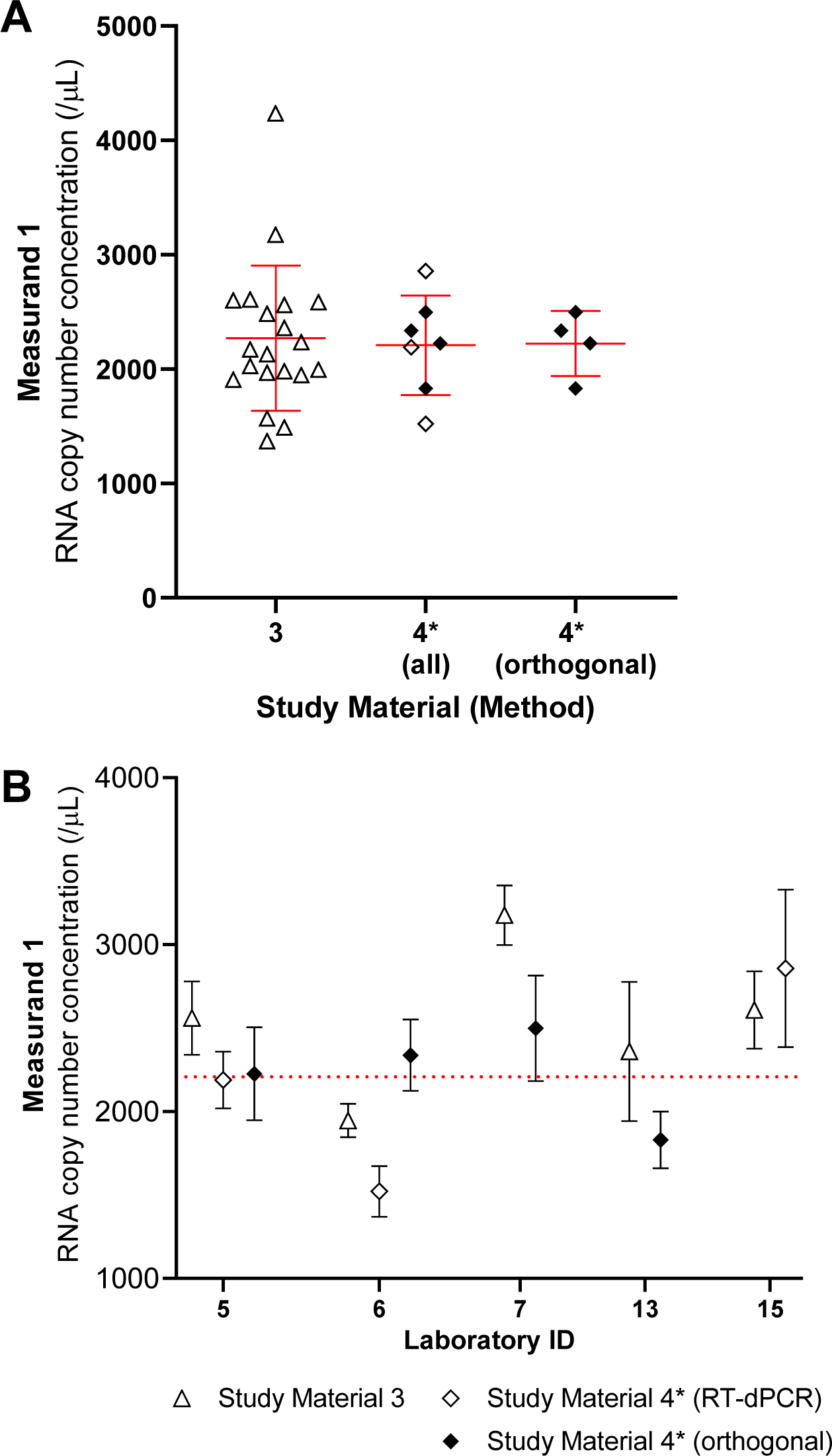
Comparison of Study Material 3 and 4 extrapolated results. (A) All nominated results for Study Material 3 are compared with Study Material 4 results (all or those using orthogonal methods). Red line and error bars show mean and SD. (B) Results for laboratories analysing both Study Materials are compared. Dotted line shows Study Material 4 extrapolated mean value. Asterisk (*) designates extrapolated Study Material 4 results. Orthogonal methods (filled diamonds) refer to non RT-dPCR approaches: ID-MS (Laboratories 5, 6 and 7) and SMFC (Laboratory 13).

## Discussion

### RT-dPCR approach and RT efficiency

Laboratory 21 was the only laboratory which performed RT-qPCR calibrated to a QC material with concentration assigned by RT-dPCR (Bio-Rad). The RT-qPCR results for Study Material 1 were consistent with most other laboratories: the result for Measurand 1 was slightly lower than the median, while the result for Measurand 2 was the highest reported, but with a large uncertainty. For Study Material 2, the Laboratory 21 result was 3-fold higher than the median for Measurand 1 and 2-fold higher than the median for Measurand 2 and not consistent with other laboratories, according to the reported uncertainty.

Three laboratories submitted results based on two-step RT-dPCR (7, 8 and 11) and Laboratory 20 provided supplementary results based on two-step RT-dPCR analysis (Appendix K). The one- and two-step RT-dPCR results are compared for Measurand 1 for Study Materials 1 to 3 in Figure 9. Alternative RT kits were used by the laboratories who performed two-step RT-dPCR, with three laboratories (7, 11, 20) using gene-specific primers and Laboratory 8 using random primers (Appendix H, Table H-1 and Appendix K). Laboratory 7 was also the only laboratory to use the QS3D dPCR platform (Thermo Fisher Scientific) compared to the majority who used the QX100/200 system (Bio-Rad). As noted in the “Interlaboratory reproducibility and consistency” section, results of laboratories 7 and 8 for Study Materials 1 and 3 were higher the average of the one-step results. Laboratory 20’s additional two-step RT-dPCR results (Appendix K) also tended to be higher than the average values. This suggests that systematic differences in the RT or PCR efficiency (“molecular dropout” [20]) or biases related to dPCR instruments such as partition volume may have led to differences between laboratories. Bias in partition volume of the QS3D is unlikely as this was characterised in-house by Laboratory 7 and DNA copy number concentration measurements performed by Laboratory 7 in CCQM P184 were consistent with other laboratories’ results [21].

**Figure 9:**
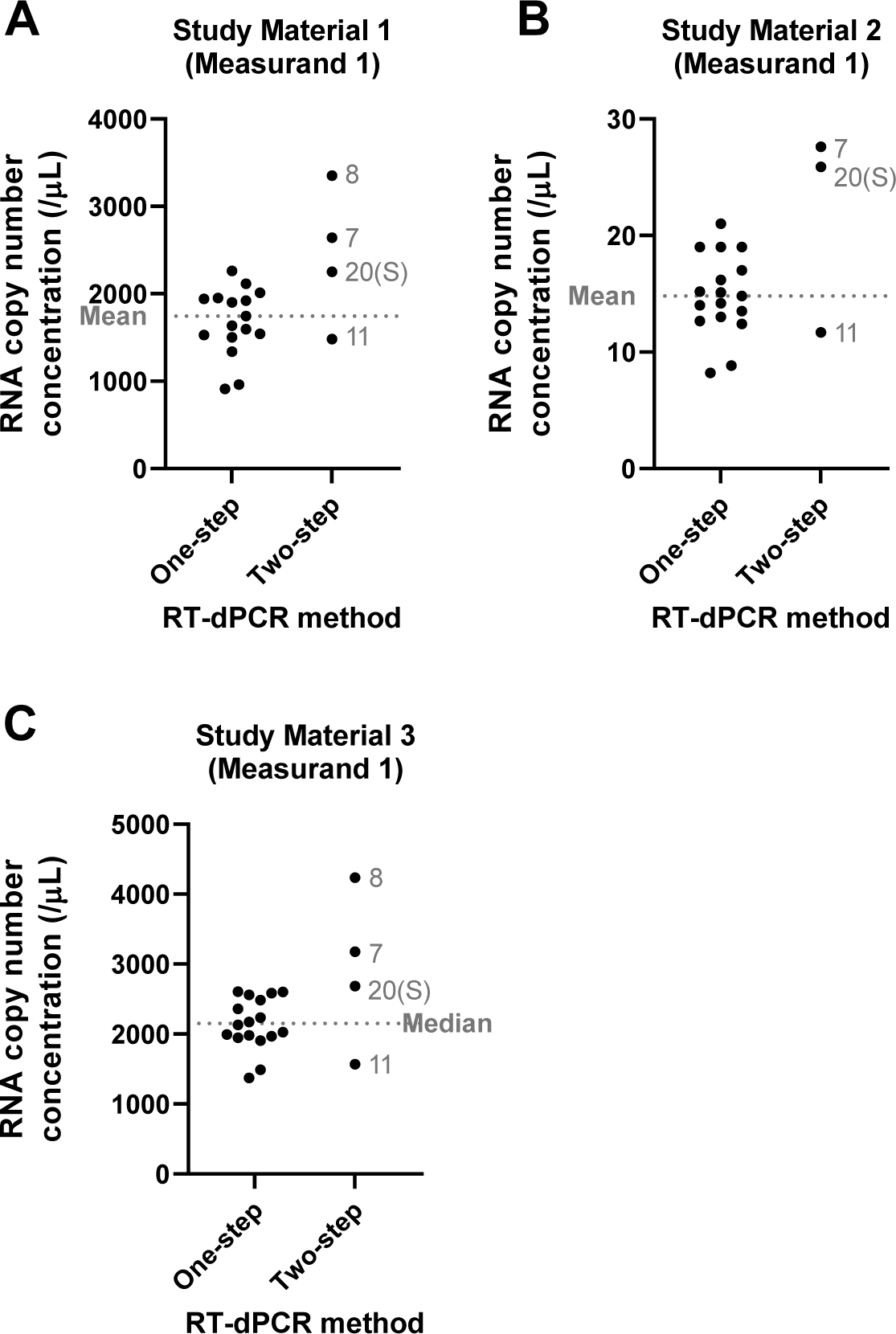
Comparison of alternative RT-dPCR approaches. Study results (nominated values) for one- and two-step RT-dPCR approaches and supplementary two-step RT-dPCR results (Laboratory 20) are shown for Study Materials 1 to 3 (Measurand 1) as black circles (labelled with Laboratory ID for two-step data). The dashed line represents the mean or median value for each Study Material as indicated.

Laboratory 5 was the only participant to include direct correction for RT efficiency in their results for Study Materials 1 to 3. Laboratory 7 also evaluated the general performance of the RT and dPCR reagents using NMIJ CRM 6204b to confirm a lack of bias. Laboratory 5’s SARS-CoV-2 assay-specific RT efficiency estimates varied between ≈ (70 to 100) %, depending on assay and calibrant (RNA oligomers or Study Material 4; personal communication, data not shown). Therefore it is possible that some laboratories’ values were affected by a negative bias due to incomplete conversion of RNA to cDNA or molecular drop out of the PCR. Non-specificity of RT priming leading to > 1 cDNA molecule per RNA may also be a cause of positive bias for two-step RT-dPCR using random primers (Laboratory 8). Following discussion of the study results at the CCQM NAWG meeting November 2020, Laboratory 8 reviewed results which were not submitted from experiments performed using the one-step RT-dPCR approach (QX200) and also performed additional measurements using Study Material 4 following the CCQM meeting (Appendix K). These RT-dPCR results were consistent with interlaboratory average values (Table 18).

### RT-dPCR assay

Primer choice and optimisation are factors which may affect both RT and PCR efficiency. For Measurand 1, laboratories predominantly used RT-dPCR applied versions of public health laboratory-developed RT-qPCR assays: US CDC N1, N2, N3 and China CDC N assay, with three laboratories (8, 10 and 13) using in house developed primers/probes for all or some of the measurements and one (11) performing measurements with both the China CDC assay and a commercial kit (Figure 10 A, B, C). The CDC N1 assay which targets a region at the 5’ end of the *N* gene was associated with lower reported results for all three Study Materials. Comparing values for one-step RT-dPCR-performing laboratories using the CDC N1 assay (*n* = 3, excluding Laboratory 5 who applied RT efficiency correction), CDC N2 (*n* = 7) and China CDC N (*n* = 4), significant differences were found by one-way ANOVA (*p* = 0.0015), with CDC N1 results lower than those for CDC N2 (*p* = 0.0011) and China CDC N (*p* = 0.033). The other approaches for *N* gene quantification appeared generally comparable with the CDC N2 and China CDC N-assay values. Analysis of predicted *N* gene secondary structure affecting the CDC N1, N2 and N3 amplicon regions (Appendix L) suggested stronger intramolecular interactions in the CDC N1 region which could be a reason for the lower observed copy number concentration results. In addition, an intramolecular hairpin in the CDC N1 reverse primer was predicted (Appendix L) which may cause reduced RT efficiency.

**Figure 10:**
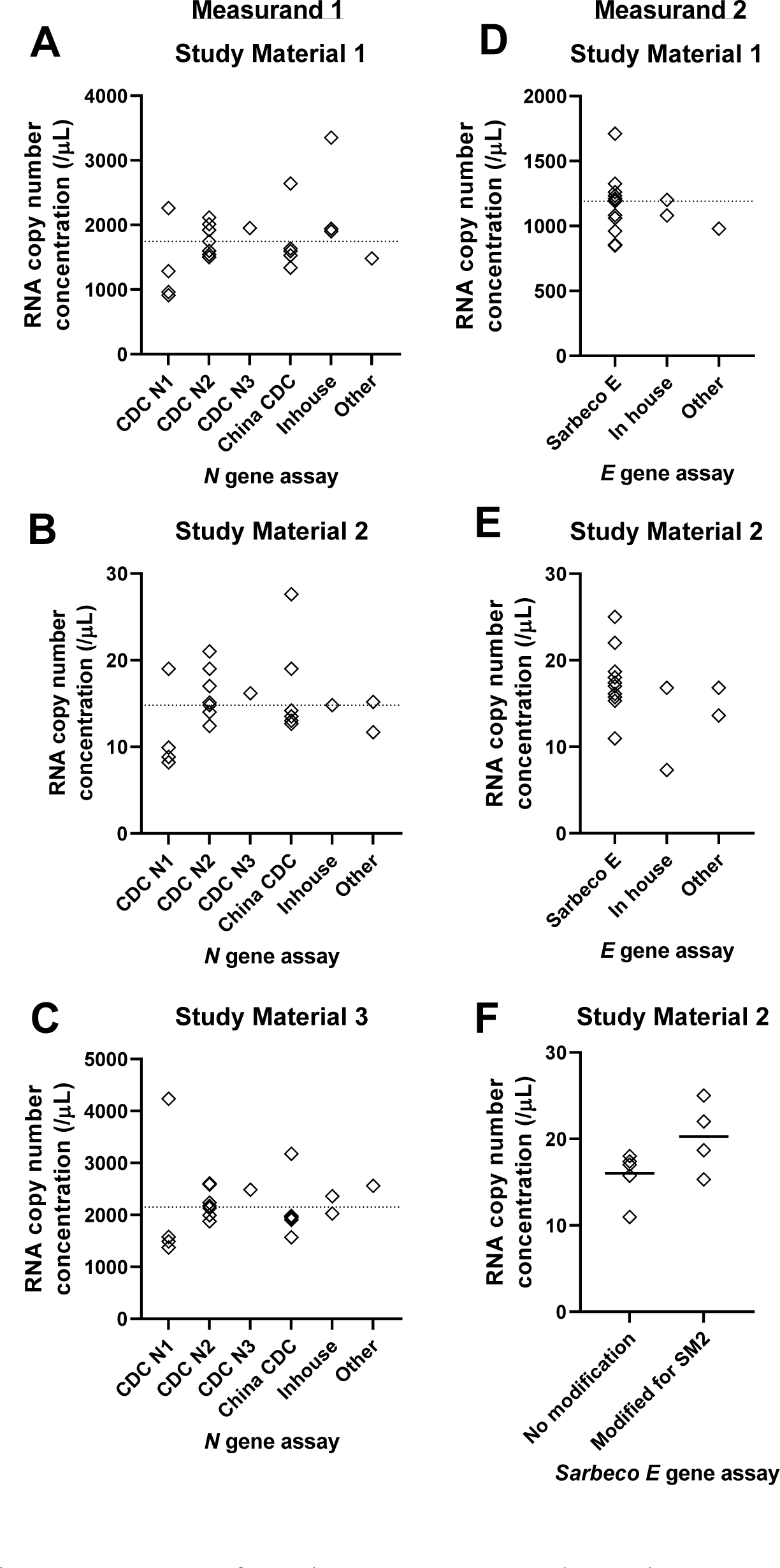
Impact of RT-dPCR assays on study results. Comparison of RT-dPCR assays used by CCQM-P199b participants. Assay used by laboratories for Measurand 1 for Study Materials 1 to 3 (A-C) and Measurand 2 for Study Materials 1 and 2 (D-E). Results for laboratories using the Sarbeco E assay with/without correction for SNVs in primer regions present in the material (F). Participants reported values shown as diamonds. Dashed line show median value of participants using RT-dPCR (A-E). Solid line shows median value (F).

For Measurand 2 (Figure 10D and 10E), laboratories predominantly used the “Sarbeco E” assay developed by Corman and colleagues [22]. Two laboratories (10 and 13) developed inhouse assays for the *E* gene and two laboratories (11 and 12) used a combination of two assays (Sarbeco E and a commercial assay (11); Sarbeco E and an inhouse design (12)). Results from the alternative approaches were comparable with the Sarbeco E assay, however for Study Material 2, Laboratory 10’s result was the lowest reported value (Figure 10E, Figure 5). This was not attributable to primer mismatches in the Study Material 2 sequence (which differed to the reference sequence (Annex 2 of the Study Protocol (Appendix C)), but may be caused by inefficiency of cDNA conversion at the low template concentrations found in the material. The Sarbeco E forward and reverse primers each contained a single base mismatch in Study Material 2, therefore some laboratories (*n* = 4) modified the primers to correct for this and others used the assay as published (*n* = 6) (Figure 10F). The use of non-modified primers did not result in a significant difference in mean *E* gene copy number concentration (*t*-test, *p* = 0.08) compared to modified primers, suggesting that the assay was tolerant of single base changes within the primer sequences.

### dPCR partition volume

Approaches to dPCR partition volume varied between laboratories (Table H4; summarised in Table 20). Six laboratories directly determined dPCR partition volume by microscopy. Laboratory 7 applied a partition volume 0.7532 nL (753.2 pL) for the QS3D system based on inhouse determination by scanning electron microscopy which was close to the manufacturer’s value of 755 pL. The partition volume of the QX100/QX200 system has been shown to vary according to type of reagents [23–26]. There were four direct measurements of QX100/200 partition volume for the One-Step RT-ddPCR Advanced Kit for Probes (Bio-Rad). Laboratory 9 performed a reagent comparison to adjust the previously determined partition volume for the ddPCR Supermixes for Probes with and without dUTP [25]. Laboratory 2 applied a directly determined value from another laboratory based on personal communication. Three laboratories applied a partition volume based on published values for ddPCR Supermix for Probes without dUTP, with Laboratory 19 adjusting this based on a reagent comparison. Ten laboratories using the QX100/200 system applied the manufacturer’s values (0.85 nL (*n* = 9), 0.868 nL (*n* = 1)). Comparing the applied manufacturer’s partition volumes with the mean of the directly determined values (Table 20), the manufacturer’s values were 10 % to 12 % higher. As partition volume is inversely proportional to copy number concentration, this would equate to a 9 % to −11 % negative bias in reported results, if the true partition volume were closer to those measured by NMIs during this study.

**Table 20:**
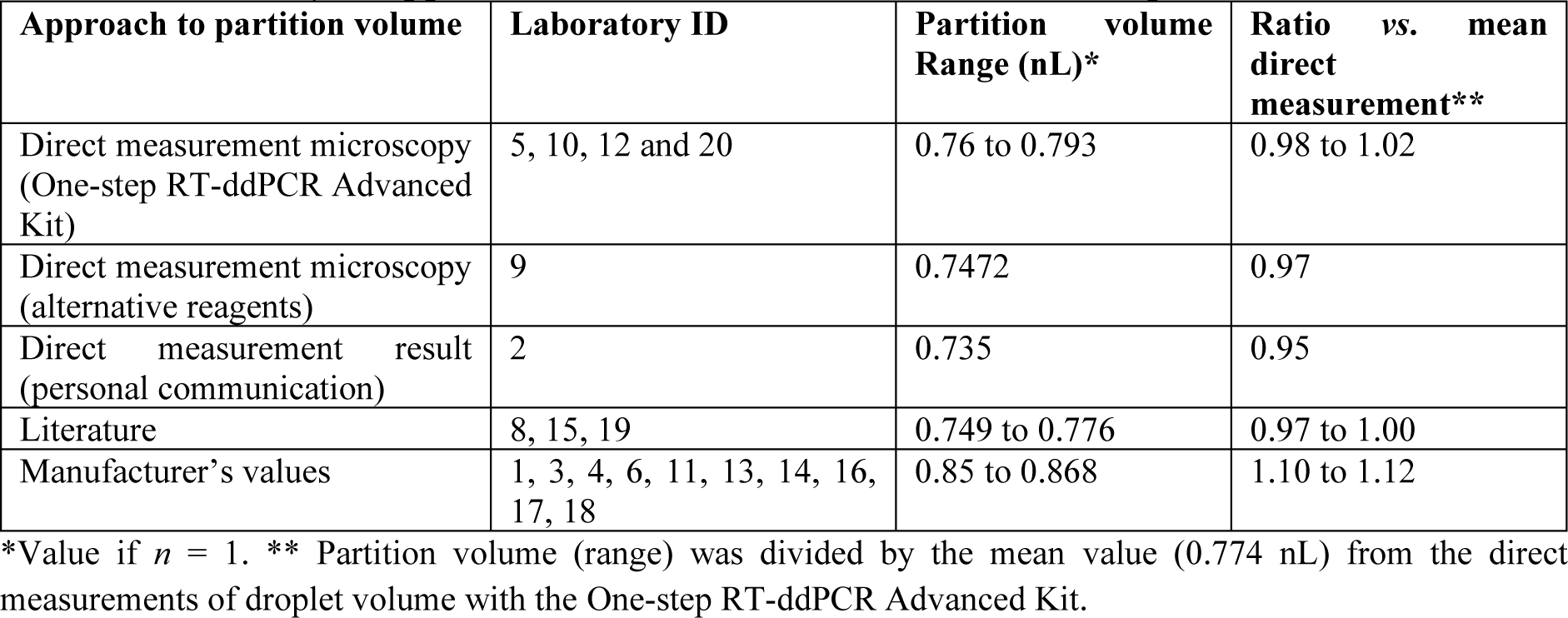
Summary of approaches and values for QX100/200 dPCR partition volume.

RNA copy number concentration results were compared between laboratories who applied a partition volume of < 0.85 nL (based on direct measurement or literature) and ≥ 0.85 nL based on the manufacturer’s values (Figure 11). As reagents may influence the underlying true partition volume, the impact of partition volume on RNA copy number concentration measurements was evaluated within the subset of data from the laboratories who used One-Step RT-ddPCR Advanced Kit for Probes (Bio-Rad), as there were few laboratories (*n* = 3) who performed two-step RT-dPCR using ddPCR Supermix for Probes (both with and without dUTP (Appendix H, Table H3)). As discussed above, the CDC N1 assay was associated with lower quantitative measurements (Figure 10), therefore results from two laboratories using this assay for their nominated results were also excluded from the analysis, as this could be another confounding factor. For Measurand 1 (*N* gene), the results reported by laboratories applying a partition volume of ≥0.85 nL were lower than the laboratories who applied a partition volume <0.85 nL for Study Materials 1, 2 and 3 (*t*-test, *p* = 0.036, *p* = 0.036 and *p* = 0.01 respectively). No significant differences were found between the grouping of laboratories’ results according to partition volume for Measurand 2 in Study Material 1 or 2.

**Figure 11:**
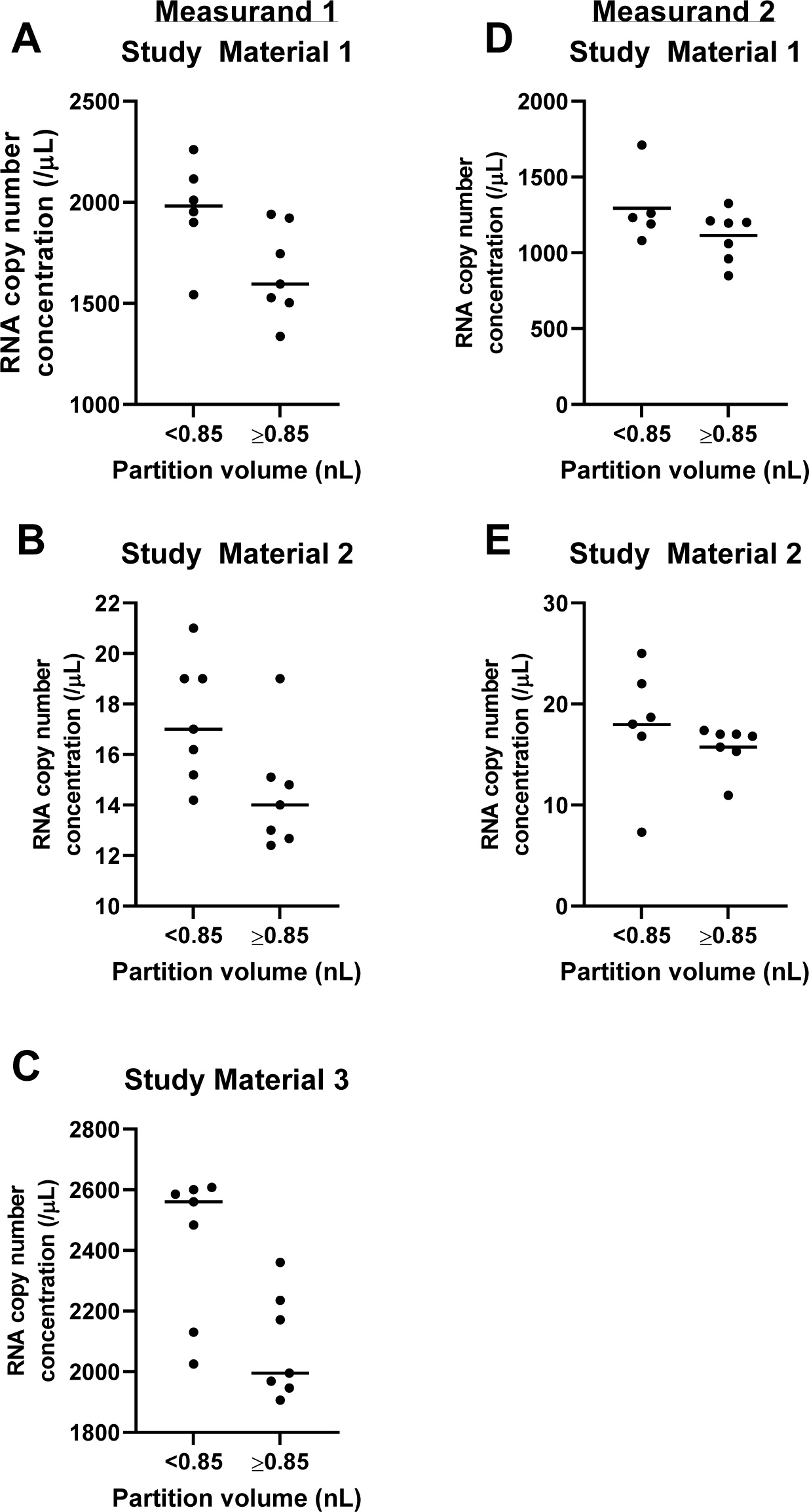
Impact of applied dPCR partition volume on study results. Comparison of a subset of CCQM-P199b results (nominated values) between laboratories applying a dPCR partition volume of <0.85 nL or ≥ 0.85 nL. For reasons explained in the text, results are included for only laboratories performing one-step RT-dPCR (Bio-Rad dPCR platform). Results from laboratories using the CDC N1 assay (Measurand 1) were excluded.

In conclusion, a systematic effect of partition volume value on RNA quantification was not observed, potentially due to additional sources of interlaboratory variation masking the effect of partition volume bias.

The impact of partition volume on the interlaboratory reproducibility of one-step RT-dPCR measurements using the QX100/QX200 system was further evaluated by modelling the scenario that all laboratories applied the same partition volume. The mean value (0.774 nL) of the One-Step RT-ddPCR Advanced Kit for Probes reagent partition volumes measured directly by participating laboratories (Table 20) was chosen to normalise results of laboratories who did not directly measure partition volume. These laboratories’ results for Study Materials 1 and 3 were scaled according to Equation 1 based on the fact that partition volume is inversely proportional to concentration. The datasets for these Materials were chosen as measurements of Study Material 2 were less precise/more influenced by stochastic variation and therefore partition volume contributed less to the uncertainty in the participants’ results.

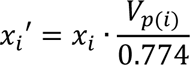

Equation 1
where *xi* is the original reported value of laboratory *i*, *xi’* is the normalised value, V_p(i)_ is the partition volume applied by the laboratory.

Table 21 compares the distribution of values as reported for laboratories performing one-step RT-dPCR (QX100/200) with and without normalization. The normalised data resulted in an increase in the mean values by 4 % to 5 % and showed improved reproducibility as indicated by a reduction in % CV and the range of results, suggesting that the application of a more standardized approach to partition volume within metrology laboratories could lead to improved interlaboratory agreement.

**Table 21:** Impact of partition volume normalisation on interlaboratory variation.

| Study Material (Measurand) | Study Material 1 (1) |  | Study Material 1 (2) |  | Study Material 3 (1) |  |
| --- | --- | --- | --- | --- | --- | --- |
| *Dataset* | *original* | *normalised* | *original* | *normalized* | *original* | *normalised* |
| Number of values | 16 | 16 | 13 | 13 | 17 | 17 |
| Minimum | 911.9 | 1001 | 848.8 | 932.2 | 1371 | 1506 |
| Maximum | 2260 | 2260 | 1710 | 1710 | 2608 | 2647 |
| Range | 1348 | 1259 | 861.2 | 777.8 | 1237 | 1141 |
| **Mean** | **1679** | **1753** | **1182** | **1239** | **2142** | **2233** |
| Std. Deviation | 383 | 363 | 205 | 201 | 369 | 330 |
| **% CV** | **22.8 %** | **20.7 %** | **17.4 %** | **16.3 %** | **17.2 %** | **14.8 %** |

Results for laboratories performing one-step RT-dPCR (QX100/200 system) are compared before and after normalization with a fixed partition volume of 0.774 nL for laboratories not directly measuring this parameter.

### Measurement uncertainty estimation

Inconsistencies between subsets of laboratories results were observed for all Study Materials, particularly in the larger Study Material 1 and 3 Measurand 1 datasets. Underestimation of measurement uncertainties (Appendix I) may have contributed to this; in particular, where laboratories estimated uncertainties taking only Type A precision data into account, very low uncertainties were reported in some cases. In addition to method (im)precision, the majority of participants included uncertainty sources due to between-unit variability (Type A or B approach) and uncertainties related to sample/reaction preparation (gravimetric or volumetric depending on procedure used). However, uncertainties related to the factors highlighted in Discussion of Results: namely assay, RT efficiency and partition volume, were not included by all laboratories. The majority of NMIs/DIs included an allowance for partition volume uncertainty, however only four laboratories measured partition volume directly during the study (Table H8), therefore the measurement uncertainty allowance may not be accurate. It is not always feasible for laboratories to investigate the highlighted factors, due to limiting time/cost (alternative assays) or equipment/capability (microscopy facility for partition volume and applying orthogonal measurements for RT efficiency).

## PILOT STUDY CONSENSUS REFERENCE VALUES (RV)

The consensus reference value (RV) was estimated following the CCQM guidance note CCQM13-22 [27]. Only nominated results from laboratories 1 and 2 were included for the Study Materials 1 to 3 datasets. Laboratory 21’s results were not included in the calculation of the consensus values for Study Materials 1, 2 and 3; the RT-qPCR approach used (calibration to a commercial QC material) was not traceable to the SI in its current format (i.e. the QC material was not assigned with a primary reference method and was not given an assigned uncertainty). Also, the Laboratory 21 results were outliers in the datasets for Study Material 1 (Measurand 2) and Study Material 2 (both Measurands) and the inclusion of the result led to the data not passing tests for a normal distribution (Table 18). Also, as previously noted, the assay used for the *N* gene was not compatible with Study Material 3, therefore the result is also not included in this dataset. For Study Material 4, the RT-dPCR and ID-MS results from laboratories 5 and 6 were considered independently in the consensus value calculations for the Study Material 4. As study material homogeneity uncertainties were low for Study Materials 1 to 3, this factor was not considered in the consensus RV uncertainty, however as Study Material 4 had a higher between-unit uncertainty (Table 3), an allowance for this was included in the consensus RV for this dataset.

Candidate RVs were calculated using classical and robust estimators as shown for each Study Material in Tables 22 to 25 and Figure 12. The recommended estimator is indicated by an asterisk (*) and further discussed below.

**Figure 12:**
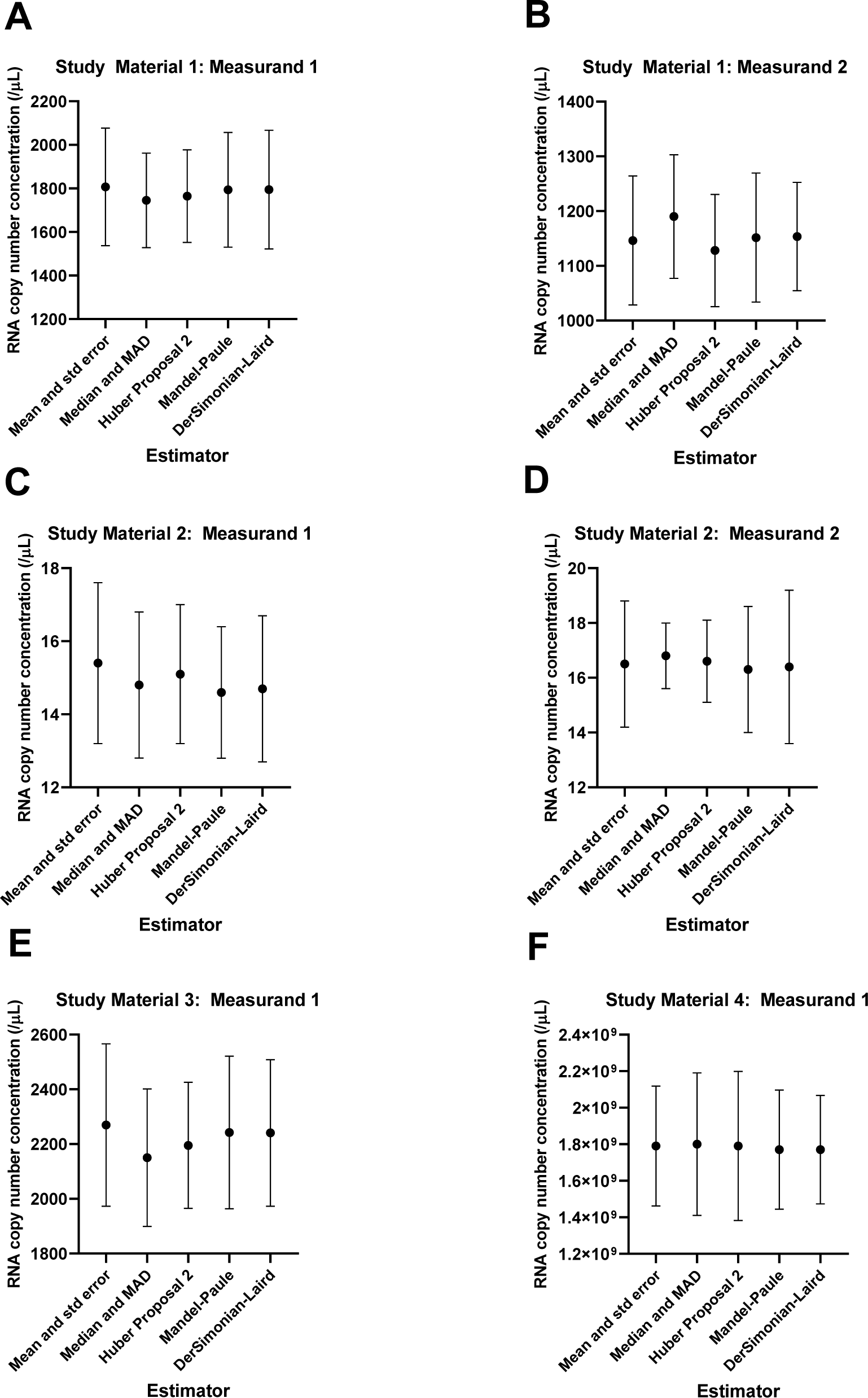
CCQM-P199b candidate consensus RVs.

**Table 22:**
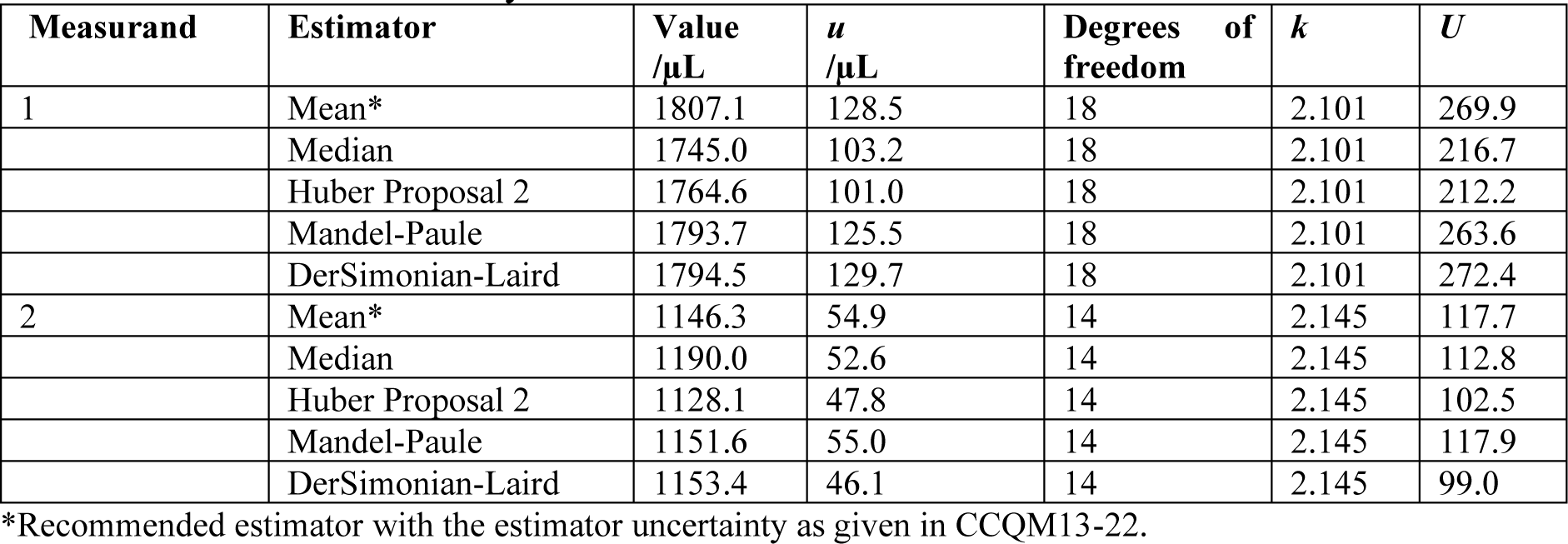
Candidate RV: Study Material 1.

| Measurand | Estimator | Value<br>/μL | *u*<br>/μL | Degrees of<br>freedom | *k* | *U* |
| --- | --- | --- | --- | --- | --- | --- |
| 1 | Mean* | 1807.1 | 128.5 | 18 | 2.101 | 269.9 |
|  | Median | 1745.0 | 103.2 | 18 | 2.101 | 216.7 |
|  | Huber Proposal 2 | 1764.6 | 101.0 | 18 | 2.101 | 212.2 |
|  | Mandel-Paule | 1793.7 | 125.5 | 18 | 2.101 | 263.6 |
|  | DerSimonian-Laird | 1794.5 | 129.7 | 18 | 2.101 | 272.4 |
| 2 | Mean* | 1146.3 | 54.9 | 14 | 2.145 | 117.7 |
|  | Median | 1190.0 | 52.6 | 14 | 2.145 | 112.8 |
|  | Huber Proposal 2 | 1128.1 | 47.8 | 14 | 2.145 | 102.5 |
|  | Mandel-Paule | 1151.6 | 55.0 | 14 | 2.145 | 117.9 |
|  | DerSimonian-Laird | 1153.4 | 46.1 | 14 | 2.145 | 99.0 |

\*Recommended estimator with the estimator uncertainty as given in CCQM13-22.

**Table 23:** Candidate RV: Study Material 2.

| Measurand | Estimator | Value /$\mu\text{L}$ | $u$ /$\mu\text{L}$ | Degrees of freedom | $k$ | $U$ |
| --- | --- | --- | --- | --- | --- | --- |
| 1 | Mean * | 15.4 | 1.0 | 18 | 2.101 | 2.2 |
|  | Median | 14.8 | 0.9 | 18 | 2.101 | 2.0 |
|  | Huber Proposal 2 | 15.1 | 0.9 | 18 | 2.101 | 1.9 |
|  | Mandel-Paule | 14.6 | 0.9 | 18 | 2.101 | 1.8 |
|  | DerSimonian-Laird | 14.7 | 1.0 | 18 | 2.101 | 2.0 |
| 2 | Mean * | 16.5 | 1.1 | 14 | 2.145 | 2.3 |
|  | Median | 16.8 | 0.6 | 14 | 2.145 | 1.2 |
|  | Huber Proposal 2 | 16.6 | 0.7 | 14 | 2.145 | 1.5 |
|  | Mandel-Paule | 16.3 | 1.1 | 14 | 2.145 | 2.3 |
|  | DerSimonian-Laird | 16.4 | 1.3 | 14 | 2.145 | 2.8 |

\*Recommended estimator with the estimator uncertainty as given in CCQM13-22.

**Table 24:** Candidate RV: Study Material 3.

| Estimator | Value /$\mu\text{L}$ | $u$ /$\mu\text{L}$ | Degrees of freedom | $k$ | $U$ |
| --- | --- | --- | --- | --- | --- |
| Mean | 2269.8 | 141.8 | 19 | 2.093 | 296.8 |
| Median * | 2150.5 | 120.1 | 19 | 2.093 | 251.3 |
| Huber Proposal 2 | 2195.2 | 110.1 | 19 | 2.093 | 230.4 |
| Mandel-Paule | 2242.5 | 133.1 | 19 | 2.093 | 278.7 |
| DerSimonian-Laird | 2240.7 | 127.8 | 19 | 2.093 | 267.5 |

\*Recommended estimator with the estimator uncertainty as given in CCQM13-22.

**Table 25:** Candidate RV: Study Material 4.

| Estimator | Value /$\mu\text{L}$ | $u$ /$\mu\text{L}$ | Degrees of freedom | $k$ | $U$ |
| --- | --- | --- | --- | --- | --- |
| Mean * | 1.79E+09 | 1.39E+08 | 7.0 | 2.36 | 3.28E+08 |
| Median | 1.80E+09 | 1.60E+08 | 6.8 | 2.45 | 3.90E+08 |
| Huber Proposal 2 | 1.79E+09 | 1.67E+08 | 6.7 | 2.45 | 4.08E+08 |
| Mandel-Paule | 1.77E+09 | 1.38E+08 | 7.0 | 2.36 | 3.26E+08 |
| DerSimonian-Laird | 1.77E+09 | 1.25E+08 | 7.3 | 2.36 | 2.97E+08 |

*Note:* Relative standard uncertainties based on distribution of participant data (according to the estimator used) were combined with a homogeneity relative uncertainty of 2.22 % (standard uncertainty/ $\sqrt{n}$ (Table 3)) where $n$ is the number of units supplied to each participant ( $n = 4$ ). Degrees of freedom were calculated using the Welch Satterthwaite equation (degrees of freedom: participant data $\nu = 6$ , homogeneity $\nu = 5$ ). \*Recommended estimator with the estimator uncertainty as given in CCQM13-22.

The candidate RVs and uncertainties were generally comparable between Study Materials and Measurands (*E* and *N* genes), although the uncertainty of median-based RV was smaller in some cases (most noticeably Study Material 2 Measurand 2). The weighted methods employed (Mandel Paul and DerSimonian Laird) were not deemed to be the most appropriate as measurement uncertainties varied in magnitude between laboratories using the same approach in the Study Material 1 to 3 datasets, as well as in the different factors which were included in uncertainty calculation (Appendix I), and notably where laboratories only included Type A precision effects in their budgets which led to very small uncertainties. Therefore, the uncertainties for these laboratories would need to be recalculated (or the results omitted) for an uncertainty weighted estimator to be valid. Likewise, the “excess variance” component modelled by these estimator is unlikely to reflect a factor which is equal in its effect on all laboratories. The Huber Proposal 2 (also known as H15) is a robust estimator which weights laboratories’ contribution to the average based on their similarity with other laboratories. This was considered not to be the most appropriate approach as it assumes that the true concentration value resides with the majority, however as highlighted previously, the majority of laboratories did not attempt to evaluate RT efficiency whereas Laboratory 5 applied a comprehensive approach to the estimation of RT efficiency. This resulted in the Laboratory 5 values being relatively high compared to the majority for Study Materials 1 and 2 and therefore further from the median and Huber estimates/uncertainties in some cases.

Therefore, to give all of the included results equal weighting, the mean (with its uncertainty estimated as the SD/√*n*) are recommended as the consensus values for the datasets which are normally distributed (all datasets with the exception of Study Material 3). The Study Material 4 dataset is also relatively small (*n* = 7) making the mean the most appropriate. Study Material 3 results were not normally distributed (Table 18) and the mean value is influenced by the high result reported by Laboratory 8. This result was confirmed as an outlier by Grubb’s test (*p* = 0.0026). Therefore, the median (with its uncertainty estimated as 1.24*MADe/√*n*) is recommended as the consensus RV for Study Material 3. The study results are shown in ascending order with the recommended estimator and expanded uncertainty in Figures 13 to 15.

**Figure 13:**
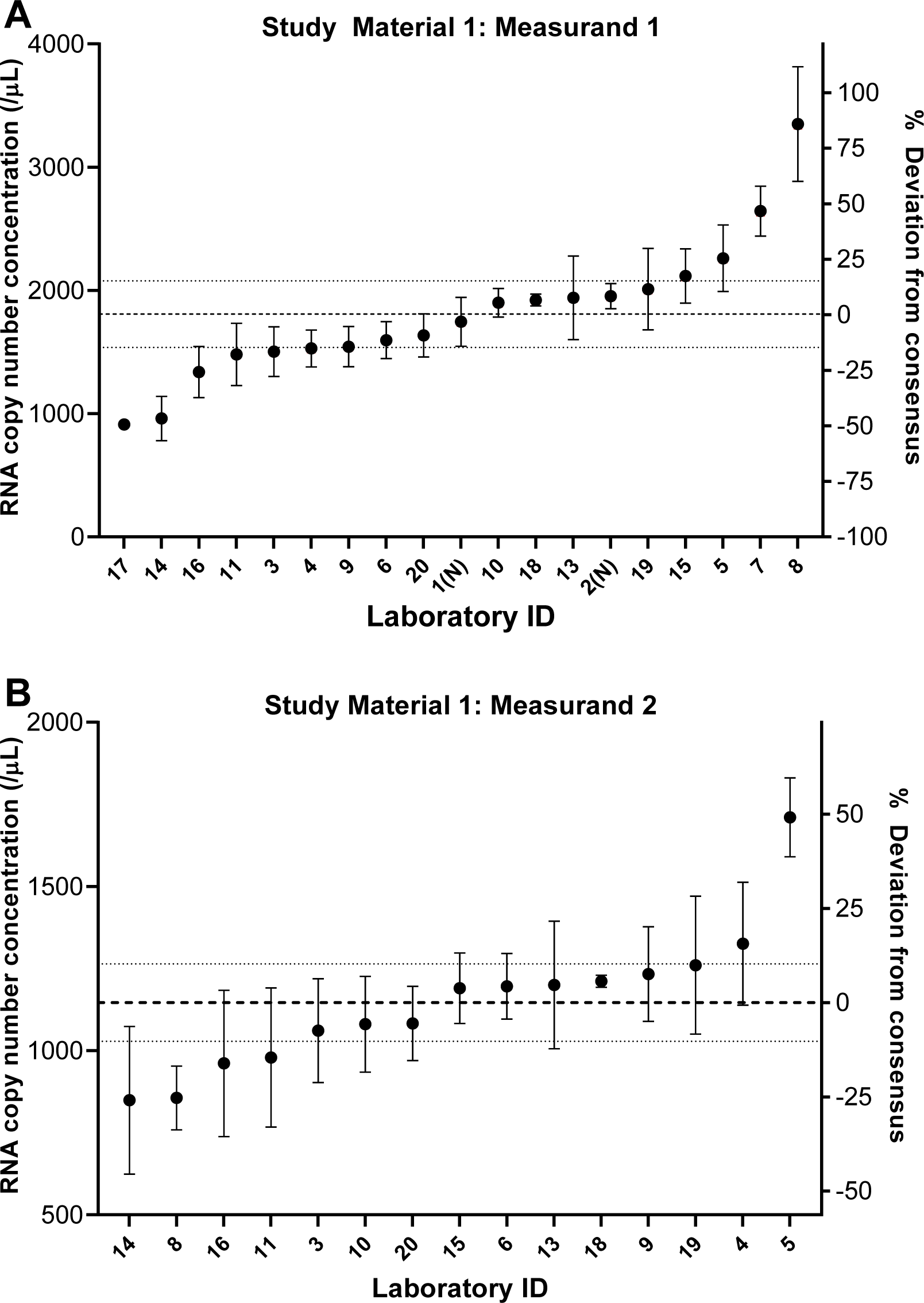
Study Material 1 consensus values, reported results and uncertainties. (A) Measurand 1. (B) Measurand 2. Dashed line showed recommended consensus value and dotted line, RV expanded uncertainty. Participants results are displayed as values (circles) and expanded uncertainty (error bars).

**Figure 14:**
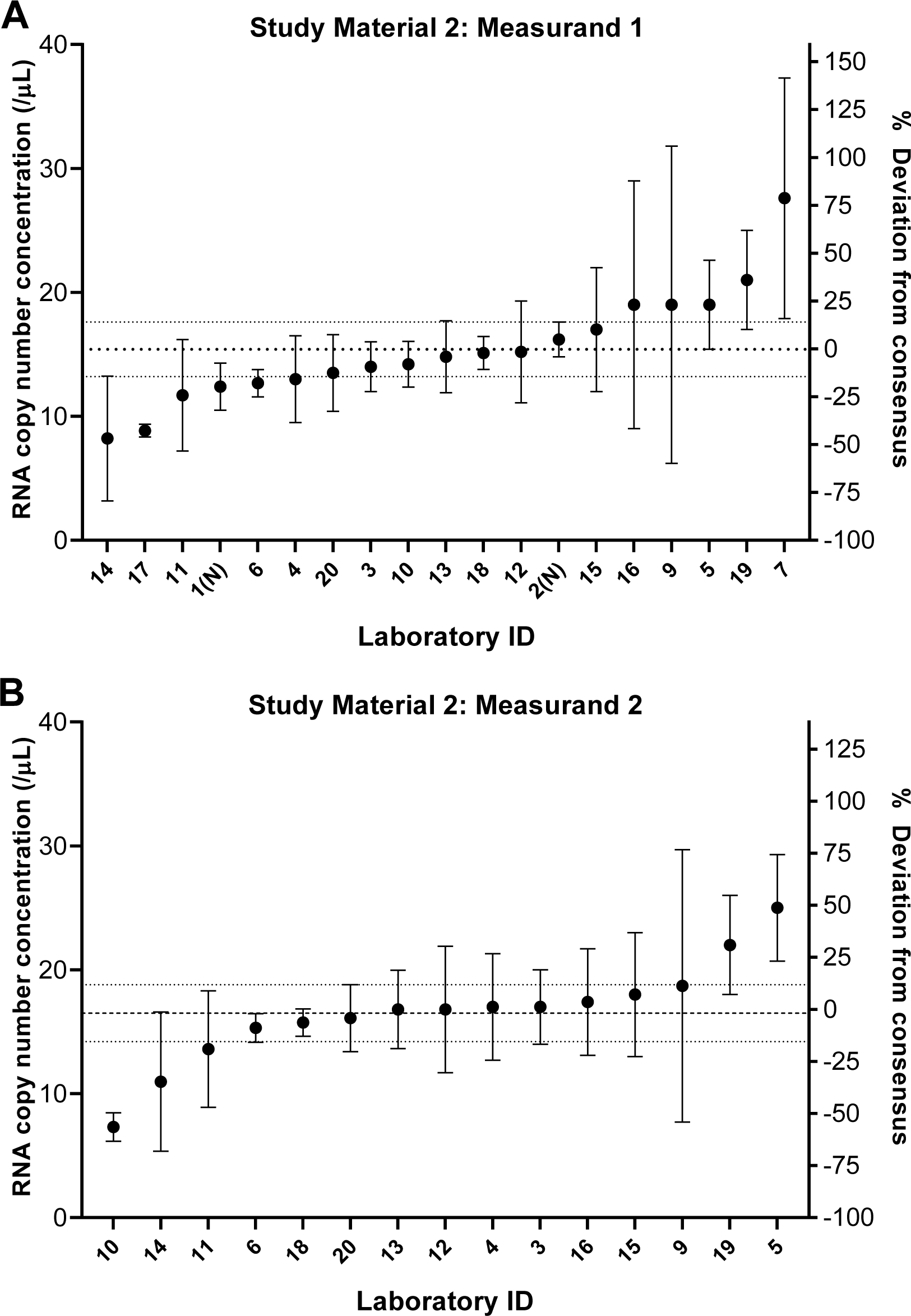
Study Material 2 consensus values, reported results and uncertainties. (A) Measurand 1. (B) Measurand 2. Dashed line showed consensus RV and dotted line, RV expanded uncertainty. Participants results are displayed as values (circles) and expanded uncertainty (error bars).

**Figure 15:**
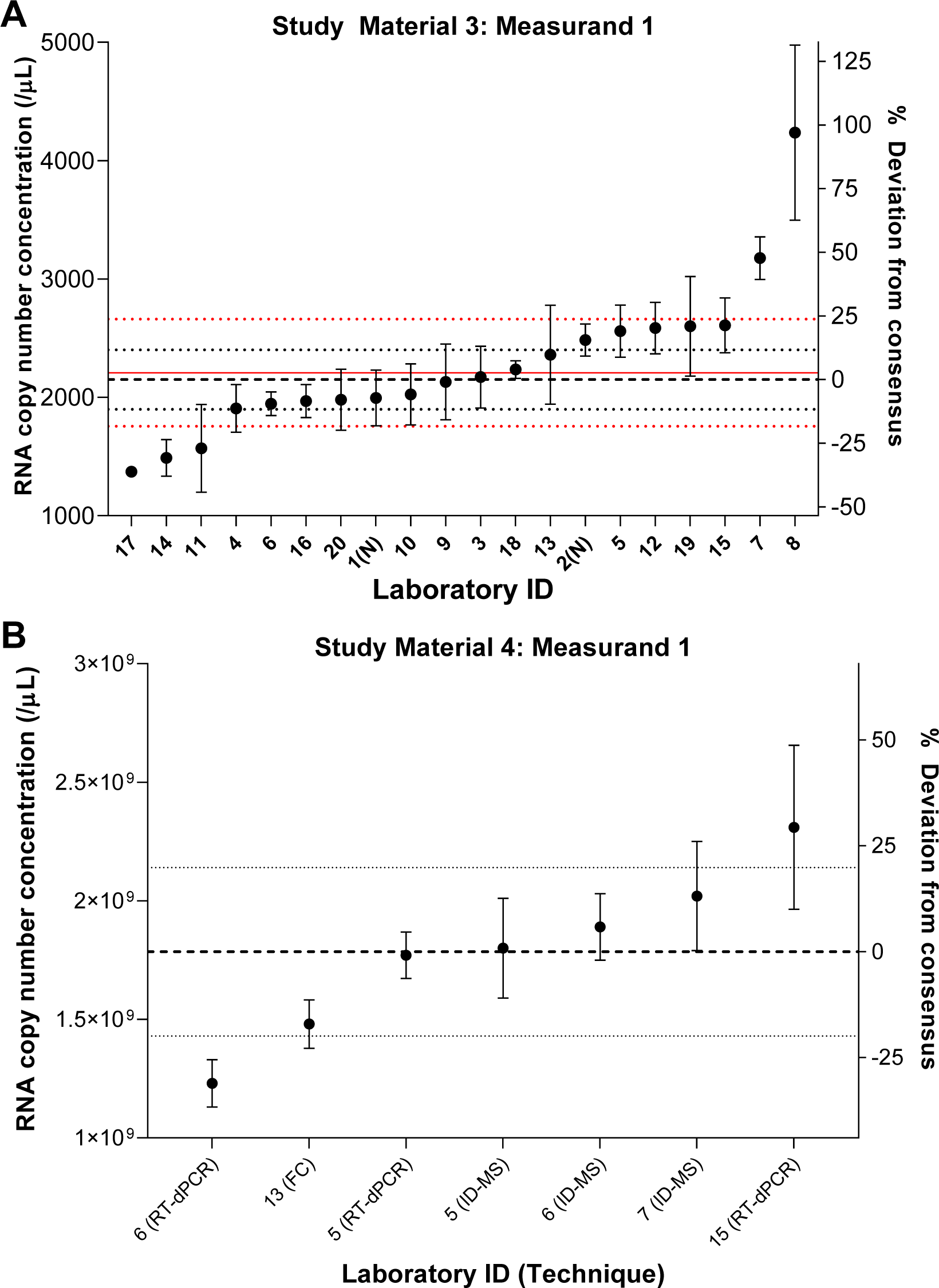
Study Material 3 and 4 consensus values, reported results and uncertainties. (A) Study Material 3 (Measurand 1). (B) Study Material 4 (Measurand 1). Black dashed and dotted lines showed RVs and expanded uncertainties. Red solid and dashed lines show Study Material 4 RV and expanded uncertainty extrapolated to the range of Study Material 3 (Table 2). Participants results are displayed as values (circles) and expanded uncertainty (error bars).

## CONCLUSIONS

CCQM-P199b assessed the capabilities of participating laboratories for targeted RNA copy number concentration measurements, and viral gene quantification in purified RNA materials. Candidate higher order methodologies were those based on enumeration: RT-dPCR and SMFC, or ID-MS methods based on traceability to amount-of-substance concentration of nucleotide standards of defined purity. RNA copy number concentration was reported for SARS-CoV-2 nucleocapsid (*N*) gene and, optionally, envelope (*E*) gene targets in number of copies per µL. Study Materials 1, 3 and 4 consisted of enzymatically synthesized RNA molecules and spanned six orders of magnitude from (10^3^ to 10^9^) /μL. Study Material 2 consisted of extracted RNA from lentiviral constructs of the SARS-CoV-2 genome and had the lowest RNA copy number concentration (≈ 15/μL) of the study materials.

Interlaboratory reproducibility of RT-dPCR, presented as the coefficient of variation (CV) in this study, was between (19 and 31) % across the four Study Materials and two measurands. Reproducibility was better for Measurand 2 (*E* gene) compared to Measurand 1 (*N* gene), which is likely to be associated with a wider variety of assays and techniques being used for Measurand 1 compared to Measurand 2. Possible sources of variability in RT-dPCR measurements include RT-dPCR approach (one-or two-step) and reagents, assay and partition volume, although systematic evaluation of these factors was not possible within the context of this study. RT-dPCR performance has previously been shown to be influenced by numerous factors including assay choice, reverse transcriptase efficiency and template type [15, 28]. Measurement of RNA copy number concentration did not appear to be influenced by choice of assay with the exceptions of the CDC N1 assay and an inhouse-designed *E* gene assay where lower results were observed. Interassay comparison and prediction of RNA secondary structure may help to identify assays with lower RT efficiency. Study results also highlighted partition volume as a source of potential bias, with wider characterisation of partition volume between instruments and reagents of the commonly used QX200 system warranted. Comparison between dPCR instruments can help to highlight systematic variation [29] but does not directly inform bias in applied partition volume, as different platforms often use alternative reagents and/or assays which can affect RT or PCR efficiency in parallel.

The trueness of RT-dPCR measurements for Study Material 3 was evaluated by comparison of results of orthogonal methods for gravimetrically linked Study Material 4, with no significant difference in mean values. This suggests that RT-dPCR measurements of the *N* gene do not show an overall negative bias, reflecting an average RT efficiency of close to 100 %. Where differences do occur, RT efficiency and assay performance may differ with different target templates which may be a cause of discrepancies with orthogonal techniques. To reduce measurement uncertainty and improve the accuracy of RT-dPCR-based quantification, further evaluation is needed, along with testing of appropriate controls for RT efficiency. The high reproducibility (CV of 5.8 %) demonstrated by the three laboratories using ID-MS suggests this approach is a candidate method for evaluation of RT efficiency controls/RT-dPCR calibrators, however as a non-sequence specific method, it is limited to synthetic RNA templates. This approach has been applied and further developed for SI-traceable DNA quantification, including the use of stable isotope-labelled DNA which acts as a control for all steps which the unknown sample undergoes [30], unlike the spiked nucleotide standards used in this study.

However, interlaboratory reproducibility suggests that overall RT-dPCR accuracy corresponds to a standard uncertainty of ≈ 20 %, which is fit-for-purpose in calibrating quantitative measurements of clinical viral genomes which can vary by several orders of magnitude. This study also supports the establishment of RMPs by providing evidence of method comparability, accuracy and reproducibility, which is required for recognition in the Joint Committee for Traceability in Laboratory Medicine (JCTLM)’s database of higher order methods [31]. Successful participation in formal, relevant international comparisons also provides evidence for calibration and measurement capability claims (CMCs) by NMIs and DIs [32], as well as underpinning NMIs’ capabilities to perform measurements in related diagnostic areas (other RNA viruses).

The development of RM and QC materials and external quality assessment (EQA) schemes by the metrology community, standards manufacturers and EQA organisers during the COVID-19 pandemic helped to address challenges in diagnostic method evaluation and lack of comparability. Therefore, the evaluation of RMPs being used by NMI/DIs [33] through CCQM-P199b underpins confidence in the values assigned to such materials and samples. This further highlights the role of reference procedures to support established material standards. This study also illustrates that RT-dPCR can be quickly deployed in this way making it an ideal approach to support the diagnostic responses to future disease outbreaks and pandemics. To support viral genome quantification in whole virus biological standards and matrix materials, approaches to develop traceability for reference measurement procedures which include RNA extraction [34] are also required. Further work should explore the role of RT-dPCR, a method capable of facilitating harmonization and supporting standardization of viral nucleic acid quantification [35, 36], in method performance evaluation and calibration hierarchies [37].

## Supporting information

CCQM P199b supplementary file Appendix A

CCQM P199b supplementary file Appendix B

CCQM P199b supplementary file Appendix C

CCQM P199b supplementary file Appendix D

CCQM P199b supplementary file Appendix E

CCQM P199b supplementary file Appendix F

CCQM P199b supplementary file Appendix G

CCQM P199b supplementary file Appendix H

CCQM P199b supplementary file Appendix I

CCQM P199b supplementary file Appendix J

CCQM P199b supplementary file Appendix K

CCQM P199b supplementary file Appendix L

## ACKNOWLEDGEMENTS

The study coordinators thank the participating laboratories for providing the requested information used in this study. NML would like to thank Stephen Ellison and Simon Cowen for their input with statistical analysis of the study results.

Points of view in this document are those of the authors and do not necessarily represent the official position or policies of the U.S. Deparent of Commerce. Certain commercial software, instruments, and materials are identified in order to specify experimental procedures as completely as possible. In no case does such identification imply a recommendation or endorsement by NIST, nor does it imply that any of the materials, instruments, or equipment identified are necessarily the best available for the purpose.

## Footnotes

1 The molecular biology community express amount of substance concentration with units of molarity (M) (SI units: mmol/L).

## REFERENCES

[1] Arnaout R, Lee RA, Lee GR, Callahan C, Cheng A, Yen CF, et al. The Limit of Detection Matters: The Case for Benchmarking Severe Acute Respiratory Syndrome Coronavirus 2 Testing. Clin Infect Dis (2021); 73(9):ee3042–e6. 10.1093/cid/ciaa1382.

[2] Jacot D, Greub G, Jaton K, Opota O. Viral load of SARS-CoV-2 across patients and compared to other respiratory viruses. Microbes Infect (2020); 22(10):617–21. 10.1016/j.micinf.2020.08.004.

3. World Health Organization. WHO In-house Assays. (2020). https://www.who.int/docs/default-source/coronaviruse/whoinhouseassays.pdf. Accessed 10th March 2021.

[4] MacKay MJ, Hooker AC, Afshinnekoo E, Salit M, Kelly J, Feldstein JV, et al. The COVID-19 XPRIZE and the need for scalable, fast, and widespread testing. Nat Biotechnol (2020); 38(9):1021–4. 10.1038/s41587-020-0655-4.

5. World Health Organization. Coronavirus disease (COVID-19) technical guidance: Laboratory testing for 2019-nCoV in humans: In-house developed molecular assays Summary document (2020). https://www.who.int/emergencies/diseases/novel-coronavirus-2019/technical-guidance/laboratory-guidance]. Accessed 2020/04/27 2020.

[6] Fung B, Gopez A, Servellita V, Arevalo S, Ho C, Deucher A, et al. Direct Comparison of SARS-CoV-2 Analytical Limits of Detection across Seven Molecular Assays. J Clin Microbiol (2020); 58(9):e01535–20. 10.1128/jcm.01535-20.

7. UK Medicines and Healthcare products Regulatory Agency. Target Product Profile: Point of Care SARS-CoV-2 Detection Tests. (2020). https://www.gov.uk/government/publications/how-tests-and-testing-kits-for-coronavirus-covid-19-work/target-product-profile-point-of-care-sars-cov-2-detection-tests]. Accessed 10th March 2021.

8. UK Medicines and Healthcare products Regulatory Agency. Target Product Profile: Laboratory-Based SARS-CoV-2 Viral Detection tests. (2020). https://www.gov.uk/government/publications/how-tests-and-testing-kits-for-coronavirus-covid-19-work/target-product-profile-laboratory-based-sars-cov-2-viral-detection-tests]. Accessed 10th March 2021.

[9] Han MS, Byun J-H, Cho Y, Rim JH. RT-PCR for SARS-CoV-2: quantitative versus qualitative. Lancet Infect Dis (2021); 21(2):165. 10.1016/S1473-3099(20)30424-2.

[10] Fajnzylber J, Regan J, Coxen K, Corry H, Wong C, Rosenthal A, et al. SARS-CoV-2 viral load is associated with increased disease severity and mortality. Nat Commun (2020); 11(1):5493. 10.1038/s41467-020-19057-5.

[11] Bustin S, Mueller R, Shipley G, Nolan T. COVID-19 and Diagnostic Testing for SARS-CoV-2 by RT-qPCR—Facts and Fallacies. Int J Mol Sci (2021); 22(5):2459. https://www.mdpi.com/1422-0067/22/5/2459.

[12] Cleveland MH, Romsos EL, Steffen CR, Olson ND, Servetas SL, Valiant WG, Vallone PM. Rapid production and free distribution of a synthetic RNA material to support SARS-CoV-2 molecular diagnostic testing. Biologicals (2023); 82:101680. 10.1016/j.biologicals.2023.101680.

13. Bentley E, Mee ET, Routley S, Mate R, Fritzsche M, Hurley M, et al. Collaborative Study for the Establishment of a WHO International Standard for SARS-CoV-2 RNA (2020).

[14] Vierbaum L, Wojtalewicz N, Grunert HP, Lindig V, Duehring U, Drosten C, et al. RNA reference materials with defined viral RNA loads of SARS-CoV-2-A useful tool towards a better PCR assay harmonization. PloS one (2022); 17(1):e0262656. 10.1371/journal.pone.0262656.

[15] Schwaber J, Andersen S, Nielsen L. Shedding light: The importance of reverse transcription efficiency standards in data interpretation. Biomol Detect Quantif (2019); 17:100077. 10.1016/j.bdq.2018.12.002.

16. Joint Committee for Guides in Metrology. International vocabulary of metrology – Basic and general concepts and associated terms (VIM). (2012). https://jcgm.bipm.org/vim/en/index.html. Accessed 08/03/2024.

[17] Kim D, Lee JY, Yang JS, Kim JW, Kim VN, Chang H. The Architecture of SARS-CoV-2 Transcriptome. Cell (2020); 181(4):914–21.e10. 10.1016/j.cell.2020.04.011.

18. Centers for Disease Control and Prevention (CDC). Research Use Only 2019-Novel Coronavirus (2019-nCoV) Real-time RT-PCR Primers and Probes. (2020). https://www.cdc.gov/coronavirus/2019-ncov/lab/rt-pcr-panel-primer-probes.html. Accessed 3rd March 2021.

[19] The dMIQE Group, Huggett J. The Digital MIQE Guidelines Update: Minimum Information for Publication of Quantitative Digital PCR Experiments for 2020. Clin Chem (2020); 66(8):1012–29. 10.1093/clinchem/hvaa125.

[20] Huggett JF, Foy CA, Benes V, Emslie K, Garson JA, Haynes R, et al. The Digital MIQE Guidelines: Minimum Information for Publication of Quantitative Digital PCR Experiments. Clin Chem (2013); 59(6):892–902. 10.1373/clinchem.2013.206375.

[21] Burke D, Devonshire AS, Pinheiro LB, Jones GM, Griffiths KR, Gonzalez AF, et al. Standardisation of cell-free DNA measurements: An International Study on Comparability of Low Concentration DNA Measurements using cancer variants. bioRxiv (2023):2023.09.06.554514. 10.1101/2023.09.06.554514.

[22] Corman VM, Landt O, Kaiser M, Molenkamp R, Meijer A, Chu DKW, et al. Detection of 2019 novel coronavirus (2019-nCoV) by real-time RT-PCR. Euro Surveill (2020); 25(3):2000045. 10.2807/1560-7917.ES.2020.25.3.2000045.

[23] Corbisier P, Pinheiro L, Mazoua S, Kortekaas AM, Chung PY, Gerganova T, et al. DNA copy number concentration measured by digital and droplet digital quantitative PCR using certified reference materials. Anal Bioanal Chem (2015); 407(7):1831–40. 10.1007/s00216-015-8458-z.

[24] Kosir AB, Divieto C, Pavsic J, Pavarelli S, Dobnik D, Dreo T, et al. Droplet volume variability as a critical factor for accuracy of absolute quantification using droplet digital PCR. Anal Bioanal Chem (2017); 409(28):6689–97. 10.1007/s00216-017-0625-y.

[25] Dagata JA, Farkas N, Kramar JA. Method for Measuring the Volume of Nominally 100 μm Diameter Spherical Water-in-Oil Emulsion Droplets. NIST Special Publication 260–184 (2016). 10.6028/NIST.SP.260-184.

[26] Pinheiro LB, Coleman VA, Hindson CM, Herrmann J, Hindson BJ, Bhat S, Emslie KR. Evaluation of a droplet digital polymerase chain reaction format for DNA copy number quantification. Anal Chem (2012); 84(2):1003–11. 10.1021/ac202578x.

[27] CCQM 13/22. CCQM guidance note: Estimation of a consensus KCRV and associated degrees of equivalence, version: 10 dated 2013-04-12.

[28] Kiselinova M, Pasternak AO, De Spiegelaere W, Vogelaers D, Berkhout B, Vandekerckhove L. Comparison of Droplet Digital PCR and Seminested Real-Time PCR for Quantification of Cell-Associated HIV-1 RNA. PloS one (2014); 9(1):e85999. 10.1371/journal.pone.0085999.

[29] Whale AS, Jones GM, Pavšič J, Dreo T, Redshaw N, Akyurek S, et al. Assessment of Digital PCR as a Primary Reference Measurement Procedure to Support Advances in Precision Medicine. Clinical Chemistry (2018); 64(9):1296–307. 10.1373/clinchem.2017.285478.

[30] Kwon HJ, Jeong JS, Bae YK, Choi K, Yang I. Stable Isotope Labeled DNA: A New Strategy for the Quantification of Total DNA Using Liquid Chromatography-Mass Spectrometry. Anal Chem (2019); 91(6):3936–43. 10.1021/acs.analchem.8b04940.

[31] JCTLM: Laboratory medicine and in vitro diagnostics. JCTLM Database: higher order methods, materials and services. https://www.jctlmdb.org/#/app/home. Accessed 08/03/2024.

[32] Bureau international des poids et mesures. CIPM Mutual Recognition Arrangement (CIPM MRA). https://www.bipm.org/en/cipm-mra.

[33] Dong L, Zhou J, Niu C, Wang Q, Pan Y, Sheng S, et al. Highly accurate and sensitive diagnostic detection of SARS-CoV-2 by digital PCR. Talanta (2021); 224:121726. 10.1016/j.talanta.2020.121726.

[34] Falak S, Macdonald R, Busby EJ, O’Sullivan DM, Milavec M, Plauth A, et al. An assessment of the reproducibility of reverse transcription digital PCR quantification of HIV-1. Methods (2021); 201:34–40. 10.1016/j.ymeth.2021.03.006.

[35] Fryer JF, Baylis SA, Gottlieb AL, Ferguson M, Vincini GA, Bevan VM, et al. Development of working reference materials for clinical virology. J Clin Virol (2008); 43(4):367–71. 10.1016/j.jcv.2008.08.011.

[36] Evans D, Cowen S, Kammel M, O’Sullivan DM, Stewart G, Grunert HP, et al. The Dangers of Using Cq to Quantify Nucleic Acid in Biological Samples: A Lesson From COVID-19. Clin Chem (2021); 68(1):153–62. 10.1093/clinchem/hvab219.

[37] ISO 17511:2020. In vitro diagnostic medical devices — Requirements for establishing metrological traceability of values assigned to calibrators, trueness control materials and human samples.

