## Supplementary material for "CCQM-P199b: Interlaboratory comparability study of SARS-CoV-2 RNA copy number quantification": CCQM P199b supplementary file Appendix A

### APPENDIX A: Sequence information

### Study Material 1

**Box A1** and **Box A2** show the relevant RNA sequences in Material 1 for Measurands 1 and 2 respectively.

**Box A1: Study Material 1 target sequence (Measurand 1)**

```
>NC_045512.2:28274-29533 Severe acute respiratory syndrome
coronavirus 2 isolate Wuhan-Hu-1, complete genome

AUGUCUGAUAAUGGACCCCAAAAUCAGCGAAAUGCACCCCGCAUUACGUUUGGUGGACCCU
CAGAUUCAACUGGCAGUAACCAGAAUGGAGAACGCAGUGGGGCGCGAUCAAAACAACGUCG
GCCCCAAGGUUUACCCAAUAAUACUGCGUCUUGGUUCACCGCUCUCACUCAACAUGGCAAG
GAAGACCUUAAAUUCCCUCGAGGACAAGGCGUUCCAAUUAACACCAAUAGCAGUCCAGAUG
ACCAAAUUGGCUACUACCGAAGAGCUACCAGACGAAUUCGUGGUGGUGACGGUAAAAUGAA
AGAUCUCAGUCCAAGAUGGUAUUUCUACUACCUAGGAACUGGGCCAGAAGCUGGACUUCCC
UAUGGUGCUAACAAAGACGGCAUCAUAUGGGUUGCAACUGAGGGAGCCUUGAAUACACCAA
AAGAUCACAUUGGCACCCGCAAUCCUGCUAACAAUGCUGCAAUCGUGCUACAACUUCCUCA
AGGAACAACAUUGCCAAAAGGCUUCUACGCAGAAGGGAGCAGAGGCGGCAGUCAAGCCUCU
UCUCGUUCCUCAUCACGUAGUCGCAACAGUUCAAGAAAUUCAACUCCAGGCAGCAGUAGGG
GAACUUCUCCUGCUAGAAUGGCUGGCAAUGGCGGUGAUGCUGCUCUUGCUUUGCUGCUGCU
UGACAGAUUGAACCAGCUUGAGAGCAAAAUGUCUGGUAAAGGCCAACAACAACAAGGCCAA
ACUGUCACUAAGAAAUCUGCUGCUGAGGCUUCUAAGAAGCCUCGGCAAAAACGUACUGCCA
CUAAAGCAUACAAUGUAACACAAGCUUUCGGCAGACGUGGUCCAGAACAAACCCAAGGAAA
UUUUGGGGACCAGGAACUAAUCAGACAAGGAACUGAUUACAAACAUUGGCCGCAAAUUGCA
CAAUUUGCCCCCAGCGCUUCAGCGUUCUUCGGAAUGUCGCGCAUUGGCAUGGAAGUCACAC
CUUCGGGAACGUGGUUGACCUACACAGGUGCCAUCAAAUUGGAUGACAAAGAUCCAAAUUU
CAAAGAUCAAGUCAUUUUGCUGAAUAAGCAUAUUGACGCAUACAAAACAUUCCCACCAACA
GAGCCUAAAAAGGACAAAAAGAAGAAGGCUGAUGAAACUCAAGCCUUACCGCAGAGACAGA
AGAAACAGCAAACUGUGACUCUUCUUCCUGCUGCAGAUUUGGAUGAUUUCUCCAAACAAUU
GCAACAAUCCAUGAGCAGUGCUGACUCAACUCAGGCCUAA
```

**Box A2: Study Material 1 target sequence (Measurand 2)**

```
>NC_045512.2:26245-26472 Severe acute respiratory syndrome coronavirus 2 isolate Wuhan-Hu-1, complete genome

AUGUACUCAUUCGUUUCGGAAGAGACAGGUACGUUAAUAGUUAAUAGCGUACUUCUUUUUC
UUGCUUUCGUGGUAUUCUUGCUAGUUACACUAGCCAUCCUUACUGCGCUUCGAUUGUGUGC
GUACUGCUGCAAUAUUGUUAACGUGAGUCUUGUAAAACCUUCUUUUUACGUUUACUCUCGU
GUUAAAAAUCUGAAUUCUUCUAGAGUUCCUGAUCUUCUGGUCUAA
```

### Study Material 2

**Boxes A3** and **A4** show the target RNA sequences for Measurands 1 and 2. Single nucleotide mutations have been randomly inserted in the SARS-CoV-2 sequences to prevent protein expression. Single nucleotide mutations which occur within the Measurand 1 and 2 target sequences are listed in Table A-1 and highlighted in **Boxes A3-A4**.

**Box A3: Study Material 2 target sequence (Measurand 1).** Positions where the construct sequence differs from the reference sequence (NC_045512.2) are shown in red.

```
>MT299805.1:9051-10016 Cloning vector pSF_lenti_SARS-CoV-2_partial-S/E/M/N, complete sequence

AUGUCUGAUAAUGGACCCCAAAAUCAGCGAAAUGCACCCCGCAUUACGUUUGGUGGACCCU
CAGAUUCAACUGGCAGUAACCAGAAUGGAGAACGCAGUGGGGCGCGAUGAAAACAACGUCG
GCCCCAAGGUUUACCCAAUAAUACUGCGUCUUGGUUGACCGCUCUCACUCAACAUGGCAAG
GAAGACCUUAAAUUCCCUCGAGGACAAGGCGUUCCAAUUAACACCAAUAGCAGUCCAGAUG
ACCAAAUUGGCUACUACCGAAGAGCUACCAGACGAAUUCGUGGUGGUGACGGUAAAAUGAA
AGAUCUCAGUCCAAGAUGGUAUUUCUACUACCUAGGAACUGGGCCAGAAGCUGGACUUCCC
UAUGGUGCUAACAAAGACGGCAUCAUAUGGGUUGCAACUGAGGGAGCCUUGAAUACACCAA
AAGAUCACAUUGGCACCCGCAAUCCUGCUAACAAUGCUGCAAUCGUGCUACAACUUCCUCA
AGGAACAACAUUGCCAUAAGGCUUCUACGCAGAAGGGAGCAGAGGCGGCAGUCAAGCCUCU
UCUCGUUCCUCAUCACGUAGUCGCAACAGUUCAAGAAAUUCAACUCCAGGCAGCAGUAGGG
GAACUUCUCCUGCUAGAAUGGCUGGCAAUGGCGGUGAUGCUGCUCUUGCUUUGCUGCUGCU
UGACAGAUUGAACCAGCUUGAGAGCAAAAUGUCUGGUAAAGGCCAACAACAACAAGGCCAA
ACUGUCACUAAGAAAUCUGCUGCUGAGGCUUCUAAGAAGCCUCGGCAAAAACGUACUGCCA
CUAAAGCAUACAAUGUAACACAAGCUUUCGGCAGACGUGGUCCAGAACAAACCCAAGGAAA
UUUUGGGGACCAGGAACUAAUCAGACAAGGAACUGAUUACAAACAUUGGCCGCAAAUUGCA
CAAUUUGCCCCCAGCGCUUCAGCGUUCUUCGGAAUGUCGCGCAUUGGCAUG
```

**Box A4: Study Material 2 target sequence (Measurand 2).** Positions where the construct sequence differs from the reference sequence (NC_045512.2) are shown in red.

```
>MT299805.1:7022-7249 Cloning vector pSF_lenti_SARS-CoV-2_partial-S/E/M/N, complete sequence

AUGUACUCAUUCGUUUCGGAAGAGACAGGUACGUAAAUAGUUAAUAGCGUACUUCUUUUUC
UUGCUUUCGUGGUAUUCUUGCUAGUUACACUAGCCAUCCUUACUGCGCUUCGAUUGUGUGC
GUAGUGCUGCAAUAUUGUUAACGUGAGUCUUGUAAAACCUUCUUUUUACGUUUACUCUCGU
GUUAAAAAUCUGAAUUCUUCUAGAGUUCCUGAUCUUCUGGUCUAA
```

Table A-1: Study Material 2 base substitutions.

| Measurand | NC_045512.2 position | Substitution (vs NC_045512.2) | MT299805 position | Measurand position |
|---|---|---|---|---|
| 1 | 28383 | C → G | 9160 | 110 |
| 1 | 28432 | C → G | 9209 | 159 |
| 1 | 28778 | A → U | 9555 | 505 |
| 2 | 26279 | U → A | 7056 | 35 |
| 2 | 26370 | C → G | 7147 | 126 |

### Study Materials 3 and 4

Study Materials 3 and 4 are composed of IVT RNA molecules at two concentrations in (i) human Jurkat cell line total RNA (Material 3) or (ii) buffered solution (Material 4). The IVT RNA is a partial sequence of the SARS-CoV-2 *N* gene (NC_045512.2: 28274-29239). Nucleotides 1-3 (underlined) of the below sequence represent the G-terminal of the T7 promoter used for in vitro transcription of RNA (1). The total sequence length is 974 nucleotides.

**Box A5: Study Materials 3 / 4 target RNA sequence (Measurand 1).** Underlined sequences correspond to the transcription initiation site of T7 RNA polymerase.

```
> NML_pEX-A128_SARS-CoV-2_partial-N_IVT
GGGAUGUCUGAUAAUGGACCCCAAAAUCAGCGAAAUGCACCCCGCAUUACGUUUGGUGGACCCUCAGAUUCAACU
GGCAGUAACCAGAAUGGAGAACGCAGUGGGGCGCGAUCAAAACAACGUCGGCCCCAAGGUUUACCCAAUAAUACU
GCGUCUUGGUUCACCGCUCUCACUCAACAUGGCAAGGAAGACCUUAAAUUCCCUCGAGGACAAGGCGUUCCAAUU
AACACCAAUAGCAGUCCAGAUGACCAAAUUGGCUACUACCGAAGAGCUACCAGACGAAUUCGUGGUGGUGACGGU
AAAAUGAAAGAUCUCAGUCCAAGAUGGUAUUUCUACUACCUAGGAACUGGGCCAGAAGCUGGACUUCCCUAUGGU
GCUAACAAAGACGGCAUCAUAUGGGUUGCAACUGAGGGAGCCUUGAAUACACCAAAAGAUCACAUUGGCACCCGC
AAUCCUGCUAACAAUGCUGCAAUCGUGCUACAACUUCCUCAAGGAACAACAUUGCCAAAAGGCUUCUACGCAGAA
GGGAGCAGAGGCGGCAGUCAAGCCUCUUCUCGUUCCUCAUCACGUAGUCGCAACAGUUCAAGAAAUUCAACUCCA
GGCAGCAGUAGGGGAACUUCUCCUGCUAGAAUGGCUGGCAAUGGCGGUGAUGCUGCUCUUGCUUUGCUGCUGCUU
GACAGAUUGAACCAGCUUGAGAGCAAAAUGUCUGGUAAAGGCCAACAACAACAAGGCCAAACUGUCACUAAGAAA
UCUGCUGCUGAGGCUUCUAAGAAGCCUCGGCAAAAACGUACUGCCACUAAAGCAUACAAUGUAACACAAGCUUUC
GGCAGACGUGGUCCAGAACAAACCCAAGGAAAUUUUGGGGACCAGGAACUAAUCAGACAAGGAACUGAUUACAAA
CAUUGGCCGCAAAUUGCACAAUUUGCCCCCAGCGCUUCAGCGUUCUUCGGAAUGUCGCGCAUUGGCAUGGGAUC
```

### REFERENCES


1. Ambion™ Life Technologies. MEGAscript® kit User Guide. 2012;Revision G.
