## Supplementary material for "CCQM-P199b: Interlaboratory comparability study of SARS-CoV-2 RNA copy number quantification": CCQM P199b supplementary file Appendix B

### APPENDIX B: Coordinating laboratory methodology

#### Coordinating laboratory methodology NIMC

##### *Construct design*

Sequences containing *E* gene (NC_045512.2:26245-26472) and *N* gene (NC_045512.2:28274-29533) of SARS-CoV-2 were synthesized by BGI (Beijing, China) to generate *in vitro* transcribed RNA molecules. These sequences was cloned into a pBluescript II SK(+) vector.

##### *In vitro transcription of RNA*

Four microgram of the SARS-CoV-2 *ORF1ab*, *E* and *N* gene plasmids were linearised with 15 U/µL *Bam*HI (1010S), 10X K buffer (1010S, both Takara) and nuclease-free water in a final reaction volume of 100 µL for 3 h at 30 °C. The digest was separated by gel electrophoresis and the corresponding bands were purified using the Universal DNA Purification Kit (DP214, TIANGEN BIOTECH (BEIJING) CO., LTD) with elution into 30 µL elution buffer. DNA concentration was estimated using Nanodrop.

To generate positive sense strand RNA *in vitro* transcription (IVT) was performed using the MEGAscript T7 kit (AM1334, ThermoFisher). Two replicate reactions were included, each containing 7.5 mM[1] of each of ATP, CTP, GTP and UTP, 1X Reaction Buffer, 2 µL T7 enzyme mix and 8 µL (approximately 0.2 µg to 1.1 µg) of plasmid. Incubation was performed at 37 °C for 4 h followed by TURBO DNase treatment. The resulting RNA was purified using the MEGAclear Kit (AM 1908, ThermoFisher). RNA transcripts were eluted in 100 µL RNase-free water. An aliquot of RNA was diluted 10-fold in The RNA Storage Solution (Ambion) and the nucleic acid concentration estimated using Nanodrop. Successful *in vitro* transcription was confirmed by analysing the 1000-fold dilution with the 2100 Bioanalyzer RNA 6000 Pico kit (Agilent) (Figure B-1). Transcripts were expected to be 1139 nt and 1572 nt in length. Total molecular mass (MM, g/mol) of the single stranded RNA transcript was estimated by multiplying the number of each nucleotide present (A, C, G, U) by the respective MM. Mass per RNA molecule (g) was calculated using the Avogadro number ($6.022 \times 10^{23}$ $mol^{-1}$). Copy number concentration in the stock RNA solution was calculated using Nanodrop results and the mass per RNA molecule in g. Diluted RNA solution were prepared at approximately $10^{10}$ /µL in RNA Storage Solution and stored at -80 °C along with the neat RNA stock.

[1] The molecular biology community express amount of substance concentration with units of molarity (M) (SI units: mmol/L).

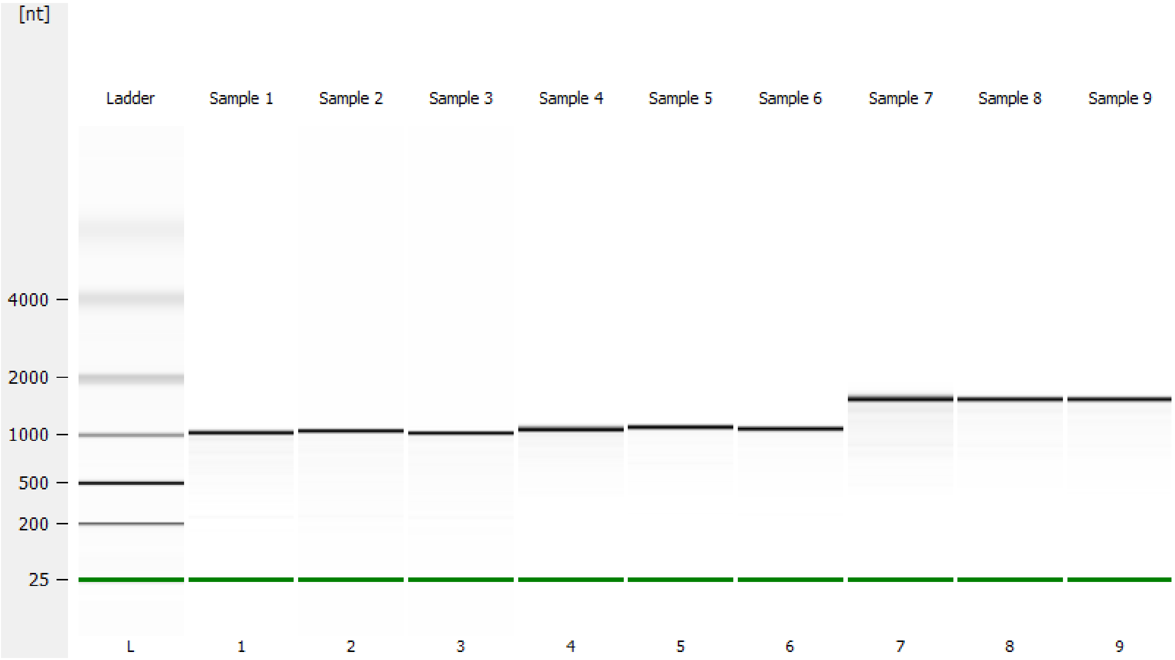


Figure B-1: Sample 1: ORF 1ab -V1 (5 ng/µL), Sample 2/3: ORF 1ab V1 (0.5 ng/µL), Sample 4: E (14 ng/µL); Sample 5/6: E (1.4 ng/µL); Sample 7: N (16 ng/µL); Sample 8/9: N (1.6 ng/µL)

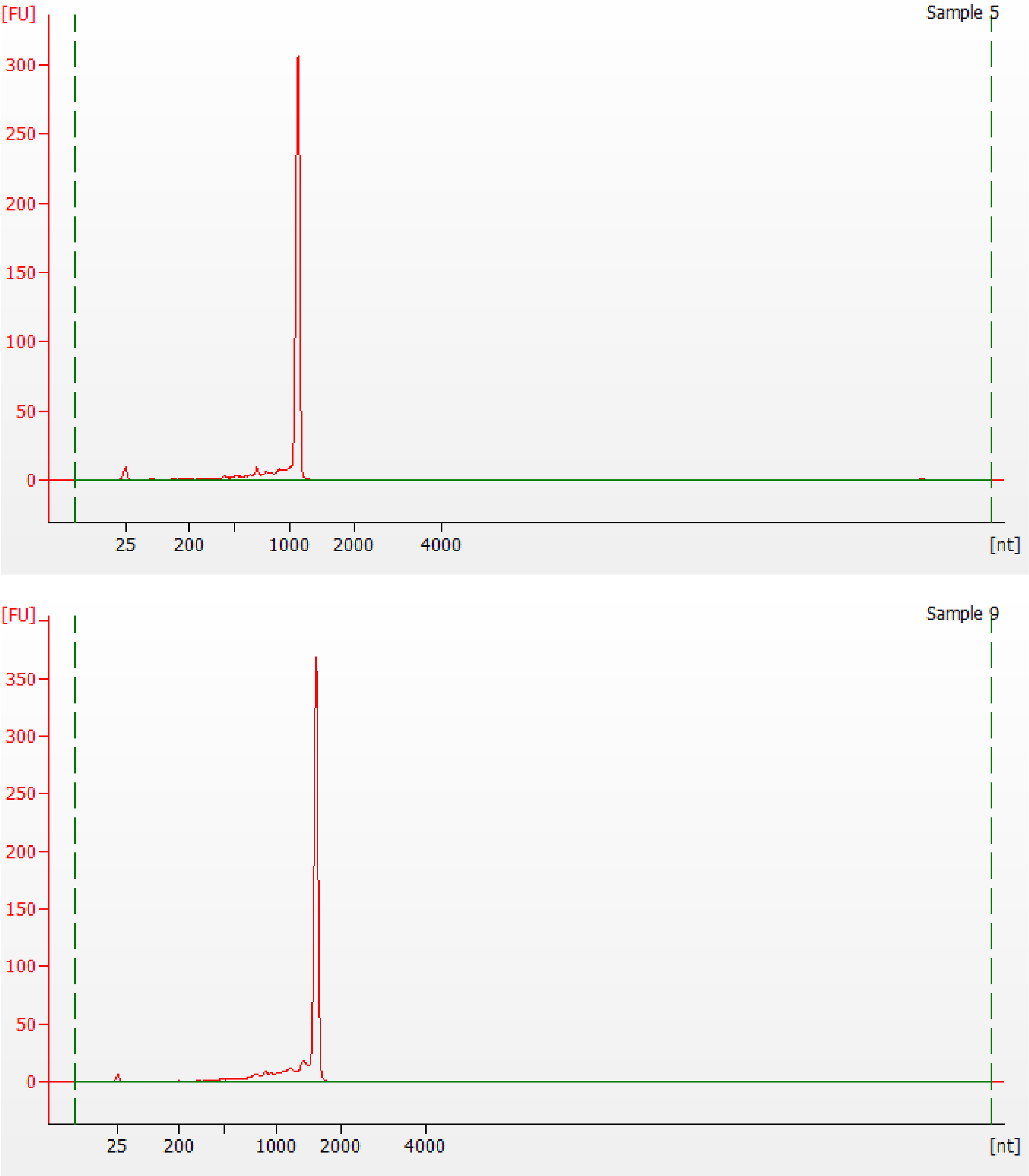


Figure B-2: In vitro transcribed RNA molecules (diluted 1:1000 in RNA Storage Solution) of *E* (top) and *N* gene (bottom) assessed with the 2100 Bioanalyzer RNA 6000 Pico kit (Agilent), confirming the presence of ≈ 1139 and ≈ 1572 nt RNA molecules, respectively.

### *RT-dPCR: Oligonucleotide sequences*

Table B-1: Oligonucleotide sequences

| **Assay designation** | **Genbank accession** | **Gene locus** | **Name** | **5' to 3' *** | **Source** |
|---|---|---|---|---|---|
| E | NC_045512.2 | 26269 to 26387 | E-F1 | ACAGGTACGTTAATAGTTAATAGCGT | (1) |
| | | | E-R2 | ATATTGCAGCAGTACGCACACA | |
| | | | E-P1 | FAM-ACACTAGCCATCCTTACTGCGCTTCGBBQ | |
| Orf1ab (duplex with E) | NC_045512.2 | 13342 to 13460 | Orf-F1 | CCCTGTGGGTTTTACACTTAA | (2) |
| | | | Orf-R2 | ACGATTGTGCATCAGCTGA | |
| | | | Orf-P1 | 5'-FAM-CCGTCTGCGGTATGTGGAAAGGTTATGG-BHQ1-3' | |
| China CDC N (duplex with RNaseP) | NC_045512.2 | 28881 to 28979 | N1 | GGGGAACTTCTCCTGCTAGAAT | (2) |
| | | | N2 | CAGACATTTTGCTCTCAAGCTG | |
| | | | N3 | FAM-TTGCTGCTGCTTGACAGATT-BHQ1 | |
| RNaseP | NR_002312.1 | 9 to 92 | RNaseP-F | GAGGGAAGCTCATCAGTGG | In-house |
| | | | RNaseP-R | CCCTAGTCTCAGACCTTCC | |
| | | | RNaseP-P | VIC-CCACGAGCTGAGTGC-MGB | |

### *RT-dPCR methodology*

One-step RT-dPCR experiments were performed using the One-Step RT-ddPCR Advanced Kit for Probes (Cat no. 1864022, Bio-Rad). Reactions were prepared in a total volume of 22 µL containing 1X [final] Supermix, 20 U/µL [final] reverse transcriptase, 15 mM [final] DTT, 4 µL of RNA template, primers and probes at a concentration of 600 nM and 50 nM for *E* gene, 600 nM and 200 nM for *N* gene, respectively. The probe was labelled with 5' FAM and 3' BHQ1. dPCR was performed using the QX200 Droplet Digital PCR System (Bio-Rad). 20 µL was pipetted into the sample well of a DG8 cartridge, and droplets generated as previously described. Thermocycling conditions were as follows: Reverse transcription at 45 °C for 10 mins, 5 mins at 95 °C, 40 cycles of 95 °C for 15 s, and 58 °C for 30 s, followed by 98 °C for 10 min and a 4 °C hold. The ramp rate for each step was 2 °C/s. Droplets were read using the QX200 Droplet Reader, and the data were analyzed using QuantaSoft version 1.7.4.0917. No Template Controls (NTCs) of nuclease-free water were employed as controls, and in all cases returned a negative result. A partition volume of 0.85 nL was used to calculate copy number concentration for preparation of the Study Materials. Data from dPCR experiments were subject to threshold and baseline setting in QuantaSoft software (Bio-Rad), and were exported as .csv files to be analysed in Microsoft Excel 2010.

*Study Material 1 preparation*

Study Material 1 (SM1) was prepared using the one-step RT-dPCR result as an initial indicator of copy number concentration of the neat SARS-CoV-2 *E*, *ORF* and *N* gene RNA transcript stock. The pooled stock of SARS-CoV-2 RNA transcripts were thawed and diluted in RNA Storage Solution with ≈ 5 ng/µL 293T human cell line total RNA (Table B-2) to prepare SM1 by three steps of dilution listed in Table B-2.

Table B-2: Preparation of SM1

| **Sample name** | **Sample type added** | **Diluent vol. added (mL)** | **Sample vol. added (µL)** | **Volumetric DF** |
|---|---|---|---|---|
| S0 | RNA stock | 3.93 | *E* gene: 21.7<br>*ORF*1ab gene: 10.4<br>*N* gene: 34.7 | *E* gene: 185<br>*ORF*1ab gene: 384<br>*N* gene: 115 |
| S1 | S0 | 24.75 | 250 | 100 |
| S2(SM1) | S1 | 9.98 | 20 | 500 |

*Homogeneity study*

Homogeneity of Study Materials (SM) 1 was assessed by one-step RT-dPCR using the China CDC N assay (Table B-1) and a duplex assay to *ORF1ab*/*E*. Ten undiluted units of SM1 were assessed, and 3 RT-dPCR replicates were analysed per unit.

*Short-term stability study*

Short-term stability (STS) of Study Materials 1 was assessed following incubation on dry ice, at 4 °C and at 27 °C for 3 days, 7 days and 14 days in comparison to a reference temperature of -80 °C. Three units of each material were included per condition. Stability was assessed by one-step RT-dPCR using ($n$ = 3) using a duplex assay to *ORF1ab*/*E*.

*Long-term stability study*

Long-term stability of the materials was assessed at 0 months, 4 months and 9 months after preparation. Three units of SM1 were analysed following storage at the reference temperature of −80 ° C. Long-term stability was assessed by one-step RT-dPCR using the China CDC *N* assay and a duplex assay to *ORF1ab*/*E* (Table B-1).

**Coordinating laboratory methodology: NIBSC**

*RNA purification from NIBSC SARS-CoV-2 Research Reagent (Study Material 2)*

RNA was extracted from two pooled 0.5 mL units of Research Reagent for SARS-CoV-2 RNA (NIBSC code 19/304) using a QIAamp UltraSens Virus Kit (Cat no 53704, Qiagen). Five replicate extractions were performed from the 1 mL pooled volume, each requiring 140 µL. Each extract was eluted in 60 µL buffer AVE (Qiagen) and pooled to give approximately 250 µL of purified RNA. This was transported to the NML on dry ice and stored at -80 °C on arrival.

**Coordinating laboratory methodology: NML**

***Study Materials 3-4: Synthetic construct design***

A region of the nucleocapsid (*N*) gene was identified from the SARS-CoV-2 reference genome (NC_045512.2: 28274-29239) for generation of a synthetic RNA test molecule. A 986 bp insert, which included a 20 bp T7 RNA polymerase promoter sequence TAATACGACTCACTATAGGG (3), was cloned into a standard pEX-A2 vector. Gene insertion in the correct orientation was confirmed by Sanger sequencing (Eurofins Genomics).

***In vitro transcription of RNA***

One microgram of the SARS-CoV-2 *N* gene plasmid was linearised with 0.4 units/µL *BamH*I-HF (R3136S), 1X CutSmart buffer (B72045, both from New England Biolabs) and nuclease-free water (AM9937) in a final reaction volume of 50 µL for 1 h at 37 °C. The digest was purified using the QIAquick PCR purification kit (28104, Qiagen) with elution into 50 µL elution buffer. Successful linearisation of a 3,450 bp molecule was confirmed using the 2100 Bioanalyzer with DNA 7500 series II kit (Agilent) (Figure B-3). DNA concentration was estimated using the Qubit 2.0 fluorometer with the dsDNA HS Assay Kit (Q32851, Invitrogen).

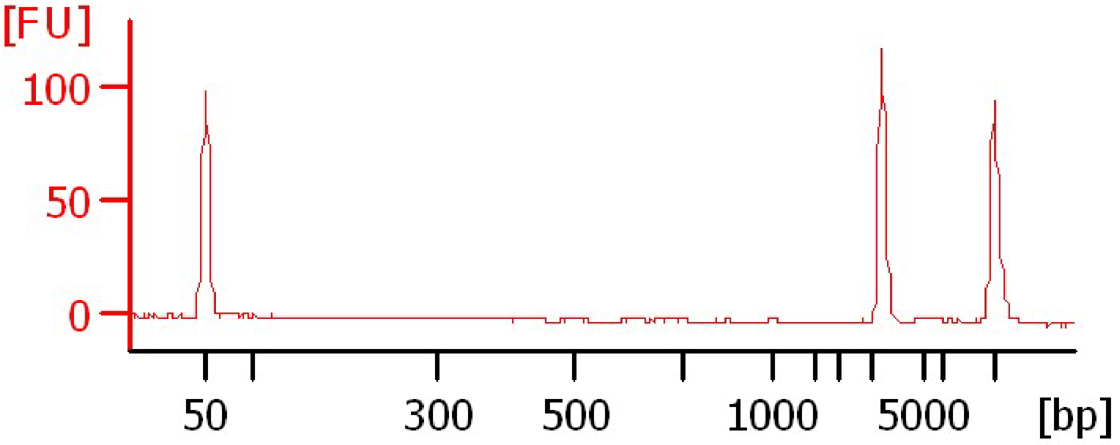


Figure B-3: Linearised plasmid assessed using the 2100 Bioanalyzer with DNA 7500 series II kit (Agilent), confirming the presence of a ≈ 3,450 bp molecule.

To generate positive sense strand RNA *in vitro* transcription (IVT) was performed using the MEGAscript T7 kit (AM1334, Life Technologies). Two replicate reactions were included, each containing 7.5 mM of each of ATP, CTP, GTP and UTP, 1X Reaction Buffer, 2 µL T7 enzyme mix and 8 µL (approximately 41 ng) of plasmid. Incubation was performed at 37 °C for 4 h followed by TURBO DNase treatment (AM2238, Life Technologies). The resulting RNA was purified using the RNeasy Mini Kit for RNA clean up protocol (Cat no 74104, Qiagen), which included an additional on-column treatment with RNase-free DNase I (Cat no 79254, Qiagen). RNA transcripts were eluted in 50 µL RNase-free water. An aliquot of RNA was diluted 10-fold in The RNA Storage Solution (Ambion) and the nucleic acid concentration estimated using the Qubit RNA HS Assay Kit (Q32852, Invitrogen). Successful *in vitro* transcription was confirmed by analysing the neat RNA stock and a 10-fold dilution with the 2100 Bioanalyzer RNA 6000 Nano kit (Agilent) (Figure B-4). Transcripts were expected to be 974 nt in length. The two IVT replicates were pooled (approximately 90 µL total volume) and the concentration confirmed again using a Qubit 2.0 fluorometer. Total molecular weight (MM, g/mol) of the single stranded RNA transcript was estimated by multiplying the number of each nucleotide present (A, C, G, U) by the respective MM. Mass per RNA molecule (g) was calculated using the Avogadro number ($6.022 \times 10^{23}$ $mol^{-1}$). Copy number concentration in the stock RNA solution was calculated using the Qubit results and the mass per RNA molecule in g. Aliquots of the pooled RNA solution were prepared at approximately $10^7$ /µL in RNA Storage Solution and stored at -80 °C along with the neat RNA stock.

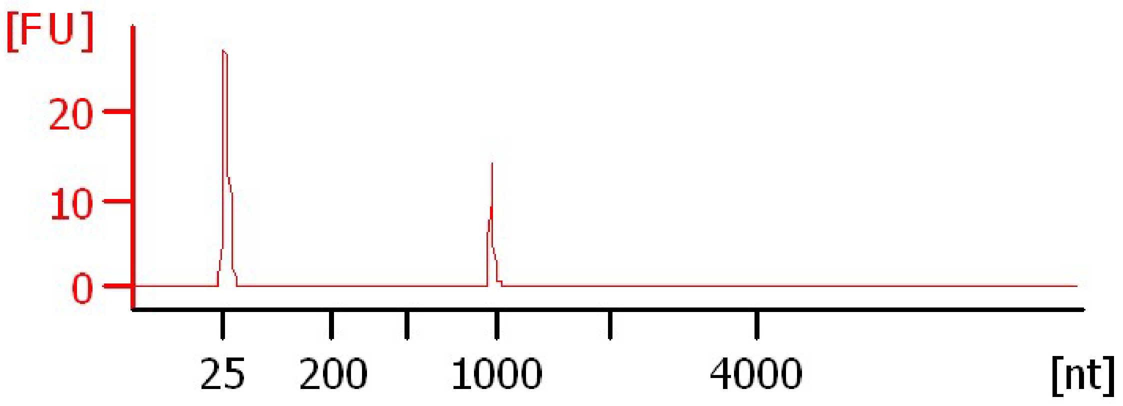


Figure B-4: In vitro transcribed RNA (diluted 1:10 in RNA Storage Solution) assessed with the 2100 Bioanalyzer RNA 6000 Nano kit (Agilent), confirming the presence of a ≈ 974 nt molecule.

#### *RT-dPCR: Oligonucleotide sequences*

Table B-3: Primer and probe sequences homologous to the SARS-CoV-2 nucleocapsid (N) gene target used for analysis and development of Study Materials

| Assay designation | GenBank accession/locus | Name | 5' to 3' * | Source |
|---|---|---|---|---|
| CDC N1 | NC_045512.2: 28287 to 28358 | Forward | GAC CCC AAA ATC AGC GAA AT | (4) |
| | | Reverse | TCT GGT TAC TGC CAG TTG AAT CTG | |
| | | Probe | ACC CCG CAT TAC GTT TGG TGG ACC | |
| CDC N2 | NC_045512.2: 29164 to 29230 | Forward | TTA CAA ACA TTG GCC GCA AA | |
| | | Reverse | GCG CGA CAT TCC GAA GAA | |
| | | Probe | ACA ATT TGC CCC CAG CGC TTC AG | |
| CDC N3 | NC_045512.2: 28681 to 28752 | Forward | GGG AGC CTT GAA TAC ACC AAA A | |
| | | Reverse | TGT AGC ACG ATT GCA GCA TTG | |
| | | Probe | AYC ACA TTG GCA CCC GCA ATC CTG | |

***Two-step RT-dPCR methodology***

The *in vitro* transcribed SARS-CoV-2 N gene RNA material was analysed initially using two-step RT-dPCR. One of the prepared aliquots (stored at a concentration of $10^7$ /µL) was diluted to approximately $10^4$ /µL in RNA Storage Solution. A cDNA synthesis reaction mix was prepared using SuperScript III First-Strand Synthesis System (18080051, Invitrogen). Each reaction contained 1X RT Buffer, 5mM.$MgCl_2$, 10mM DTT, 2 U/µL RNaseOUT, 10 U/µL SuperScript III reverse transcriptase, 0.5 nM dNTP mix, 0.1 µM gene specific reverse primer for CDC N1, N2 or N3 assay (Table B-3), 2.5 µL RNA template and DEPC-treated water to a final volume of 10 µL. RNA templates were heat denatured at 65 °C for 5 mins and immediately quenched on ice prior to addition into the reaction. cDNA synthesis was performed at 50 °C for 50 mins, with an enzyme inactivation step at 85 °C for 5 mins. Resulting cDNA was incubated at 37 °C for 20 mins with 0.5 µL of RNase H before being stored at -20 °C prior to dPCR.

dPCR was performed on the QX200 Droplet Digital PCR System (Bio-Rad). cDNA was diluted 10-fold in nuclease-free water (Ambion), and 5.5 µL was added to a prepared reaction volume of 22 µL containing 1X ddPCR Supermix for Probes without dUTP (Bio-Rad), sterile nuclease-free water and either the CDC N1, N2 or N3 primers and probes (Table B-3). Probes were labelled with 5' FAM and contained a 3'BHQ1 quenching moiety. Primers were included at a concentration of 900 nM, and probes at a concentration of 250 nM. 20 µL was pipetted into the sample well of a DG8 cartridge, and droplets generated as previously described (5). Thermocycling conditions were as follows: 10 mins at 95 °C, 40 cycles of 94 °C for 30 s, and 55 °C for 1 min, followed by 98 °C for 10 min and a 4 °C hold. The ramp rate for each step was 2 °C/s. Droplets were read using the QX200 Droplet Reader, and the data were analyzed using QuantaSoft version 1.7.4.0917. No Template Controls (NTCs) of nuclease-free water were employed as controls, along with RT negative controls (water in place of reverse transcriptase), and in all cases returned a negative result. A partition volume of 0.834 nL was used to calculate copy number concentration. Data from dPCR experiments were subject to threshold and baseline setting in QuantaSoft software (Bio-Rad), and were exported as .csv files to be analysed in Microsoft Excel 2010. The average number of copies per droplet ($\lambda$) was calculated as described previously (6).

***One-step RT-dPCR methodology***

One-step RT-dPCR experiments were performed using the One-Step RT-ddPCR Advanced Kit for Probes (Cat no. 1864021, Bio-Rad). Reactions were prepared in a total volume of 22 µL containing 1X [final] Supermix, 20 U/µL [final] reverse transcriptase, 15 mM [final] DTT, 5.5 µL of RNA template, nuclease-free water and primers and probes at a concentration of 900 nM and 250 nM, respectively. For one-step RT-dPCR, only the CDC N2 assay was used (Table B-3). The probe was labelled with 5' FAM and was BHQnova double-quenched. RNA templates were heat denatured at 65 °C for 5 mins and immediately quenched on ice prior to addition into the reaction. dPCR was performed using the QX200 Droplet Digital PCR System (Bio-Rad). 20 µL was pipetted into the sample well of a DG8 cartridge, and droplets generated as previously described (5). Thermocycling conditions were as follows: Reverse transcription at 47.5 °C for 60 mins, 10 mins at 95 °C, 40 cycles of 95 °C for 30 s, and 55 °C for 1 min, followed by 98 °C for 10 min and a 4 °C hold. The ramp rate for each step was 2 °C/s. Droplets were read using the QX200 Droplet Reader, and the data were analyzed using QuantaSoft version 1.7.4.0917. No Template Controls (NTCs) of nuclease-free water were employed as controls, along with RT negative controls (water in place of reverse transcriptase), and in all cases returned a negative result. A partition volume of 0.776 nL was used to calculate copy number concentration for preparation of the study materials. Data from dPCR experiments were subject to threshold and baseline setting in QuantaSoft software (Bio-Rad), and were exported as .csv files to be analysed in Microsoft Excel 2010. The average number of copies per droplet (λ) was calculated as described previously (6).

***Gravimetry***

Gravimetric preparation of Study Materials and the Study Material 4 homogeneity study was performed using a Mettler Toledo XP205 balance to 5 decimal places. Following cleaning of the balance, linearity was tested using a set of laboratory standard weights covering the range 0.1 g to 200 g. Standard uncertainty of measurement for the balance was ± 0.000159 g.

For the Short-term and long-term stability assessment of Study Material 4, an Ohaus E10640 balance reporting to 4 decimal places was used. Following cleaning of the balance, linearity was tested using a set of laboratory standard weights covering the range 0.001 g to 0.5 g. Standard uncertainty of measurement for the balance was ± 0.000085 g

***Study Material 3 and 4 preparation***

Study Material 4 (SM4) was prepared using the two-step RT-dPCR result as an initial indicator of copy number concentration of the neat SARS-CoV-2 RNA N gene transcript stock. An average of the results obtained using the CDC N1, CDC N2 and CDC N3 assays was taken. A target dilution factor (DF) of 89.16 was chosen to prepare SM4 from the neat stock. The pooled stock of SARS-CoV-2 N gene RNA transcript was thawed and diluted gravimetrically in RNA Storage Solution (Table B-4).

Table B-4: Gravimetric preparation of SM4.

| **Mass of tube (g)** | **Diluent vol. added (mL)** | **Mass of tube + diluent (g)** | **Mass of diluent (g)** | **Sample vol. added (mL)†** | **Mass of tube + diluent + sample (g)** | **Mass of sample (g)** | **Volumetric DF** | **Gravimetric DF** |
|---|---|---|---|---|---|---|---|---|
| 14.4610 | 6.526 | 20.8824 | 6.421 | 0.074 | 20.9485 | 0.066 | 89.16 | 98.1467 |

† <74 µL of sample was found to be available following thawing.

The dilution factor to prepare SM4 from the stock solution was 98.15 (± 0.33) based on gravimetry. The prepared solution of SM4 was placed on a roller mixer at 4 °C for 30 mins to obtain a homogenous solution prior to aliquoting the units. A total of 58 units were prepared using a E3 multipipette (Eppendorf) to a volume of 100 µL. Filling accuracy of the pipette was checked prior to aliquoting of the units using RNA Storage Solution (Table B-5).

Table B-5: Record of filling accuracy using 100 µL (0.1 g) of RNA Storage Solution (RSS).

| **Measurement** | **Weight of tube (g)** | **Weight after filling with 100 µL RSS (g)** | **Added (g)** | **Vol according to mass (µL)** |
|---|---|---|---|---|
| 1 | 1.0053 | 1.103 | 0.0977 | 97.7 |
| 2 | 1.0042 | 1.1056 | 0.1014 | 101.4 |
| 3 | 1.0040 | 1.1036 | 0.0996 | 99.6 |
| 4 | 0.9978 | 1.0975 | 0.0997 | 99.7 |
| 5 | 0.9957 | 1.0957 | 0.1000 | 100.0 |
| 6 | 0.9952 | 1.0947 | 0.0995 | 99.5 |
| 7 | 1.0083 | 1.1083 | 0.1000 | 100.0 |
| 8 | 0.9978 | 1.0979 | 0.1001 | 100.1 |
| 9 | 0.9910 | 1.0912 | 0.1002 | 100.2 |
| 10 | 0.9980 | 1.0989 | 0.1009 | 100.9 |
| | | **Mean** | 0.0999 | |
| | | **SD** | 0.0010 | |

Following gravimetric preparation of SM4, ten 55 µL aliquots were retained for RT-dPCR analysis, and for preparation of Study Material 3 (SM3). For three separate aliquots, volumetric dilutions were prepared to an input concentration of 5 x $10^3$ /µL (based on the two-step RT-dPCR result) and analysed by one-step RT-dPCR using the CDC N2 assay. SM3 was gravimetrically prepared based on the one-step RT-dPCR result using a thawed 55 µL aliquot of SM4 (termed 'Mat 4'). A target dilution factor (DF) of 7.89 x $10^5$ was planned to prepare SM3 from SM4, using Human Jurkat cell total RNA as carrier (Ambion AM7858). Carrier solution was gravimetrically prepared to a concentration of 2.5 ng/µL in RNA Storage Solution using the same 5-figure balance as used to prepare SM4 and SM3.

Table B-6: Gravimetric preparation of SM3.

| Sample | Mass of tube (g) | Diluent vol. added (µL) | Mass of tube + diluent (g) | Mass of diluent (g) | Sample vol. added (µL) | Mass of tube + diluent + sample (g) | Mass of sample (g) | Volumetric DF | Gravimetric DF |
|---|---|---|---|---|---|---|---|---|---|
| D1 | 1.1014 | 450.0 | 1.5471 | 0.4457 | 50.0 | 1.5963 | 0.0492 | 10.0 | 10.0589 |
| D2 | 1.0977 | 950.0 | 2.0363 | 0.9386 | 50.0 | 2.0862 | 0.0499 | 20.0 | 19.8096 |
| D3 | 1.0977 | 900.0 | 2.0076 | 0.9099 | 100.0 | 2.1064 | 0.0988 | 10.0 | 10.2095 |
| D4 | 1.0956 | 900.0 | 1.9883 | 0.8927 | 100.0 | 2.0876 | 0.0993 | 10.0 | 9.9899 |
| SM3 | 12.9977 | 30700.0 | 43.6529 | 30.6552 | 798.1 | 44.4432 | 0.7903 | 39.5 | 39.7893 |

Based on gravimetry, there was a dilution factor of $8.09 \times 10^5$ ($\pm 5.83 \times 10^3$) between SM4 and SM3 (Table B-6). The prepared solution of SM3 was placed on a roller mixer at 4 °C for 30 mins to obtain a homogenous solution prior to aliquoting the units. A total of 310 units were prepared using a E3 multipipette (Eppendorf) to a volume of 100 µL.

***Study Material 2 preparation***

Purified RNA from Research Reagent for SARS-CoV-2 RNA (NIBSC code 19/304) received from NIBSC was thawed and two 10 µL aliquots removed for analysis, and the RNA stock solution returned to storage at -80 °C.

An initial one-step RT-dPCR experiment was performed to determine the CDC N2 target copy number concentration of the purified RNA. Based on the result, Study Material 2 (SM2) was gravimetrically prepared in Human Jurkat cell total RNA (Ambion AM7858), which had been gravimetrically prepared to a concentration of 2.5 ng/µL in RNA Storage Solution. A target dilution factor (DF) of 99.56 was planned to prepare the material.

Table B-7: Gravimetric preparation of SM2.

| Sample | Mass of tube (g) | Diluent vol. added (mL) | Mass of tube + diluent (g) | Mass of diluent (g) | Sample vol. added (mL) | Mass of tube + diluent + sample (g) | Mass of sample (g) | Volumetric DF | Gravimetric DF |
|---|---|---|---|---|---|---|---|---|---|
| D1 | 1.0971 | 1.530 | 2.6079 | 1.511 | 0.220 | 2.8209 | 0.213 | 8.0 | 8.0930 |
| SM2 | 12.9215 | 19.322 | 32.0576 | 19.136 | 1.678 | 33.7066 | 1.649 | 12.5 | 12.6047 |

Based on gravimetry, a dilution factor of 102.01 (±0.11) was applied to prepare SM2 (Table B-7). The prepared solution of SM2 was placed on a roller mixer at 4 °C for 30 mins to obtain a homogenous solution prior to aliquoting the units. A total of 206 units were prepared using a E3 multipipette (Eppendorf) to a volume of 100 µL. All materials were stored at -80 °C.

***Homogeneity study***

Homogeneity of Study Materials (SM) 2, 3 and 4 was assessed by one-step RT-dPCR using the CDC N2 assay (Table B-3). Ten undiluted units of each of SM2 and SM3 were assessed, and 8 RT-dPCR replicates were analysed per unit.

For homogeneity testing of SM4, 6 units were diluted gravimetrically in 0.5 ng/µL Human Jurkat cell total RNA using a Mettler Toledo XP205 balance. Following cleaning of the balance, linearity was tested using a set of laboratory standard weights covering the range 0.01 g to 200 g. Standard uncertainty of measurement for the balance was ±0.000159 g. Two independent dilution series were per unit were included in the analysis. A target dilution factor (DF) of $8.10 \times 10^5$ was planned per dilution series, so that the final dilution of each unit of SM4 could be analysed directly by RT-dPCR. An aliquot of the lowest dilution from each series per unit ($n$ = 12) was analysed in triplicate, with 5.5 µL template added to a prepared volume of 22 µL. 20 µL of the reaction mix was pipetted into a DG8 cartridge, and RT-dPCR was performed using the QX200 digital PCR system as described above. In addition to RT-dPCR, the 6 undiluted units of SM4 were analysed in triplicate on a Qubit 2.0 fluorometer. 10 µL of RNA was added to a total reaction volume of 200 µL including Qubit RNA HS buffer and reagent.

Datasets are not balanced, so mixed effects models with maximum likelihood estimation were used to obtain the unit-to-unit variances. Plate row and column were initially included as fixed effects in order to check for significance. This was decided by comparison of model AIC value. For both M2 and M3 the lowest AIC was obtained without either row or column.

***Short-term stability study***

Short-term stability (STS) of Study Materials 2 and 3 was assessed following incubation on dry ice, at 4 °C and at 27 °C for 3 days and 7 days in comparison to a reference temperature of -80 °C. Three units of each material were included per condition. For Study Material 4, two units were incubated on dry ice and at 27 °C for 7 days, and at 4 °C for 18 days (in error), and compared to the reference temperature. Each unit of SM4 was diluted gravimetrically in 0.5 ng/µL Human Jurkat cell total RNA carrier using an Ohaus E10640 balance. Two independent dilution series per unit were included in the analysis for estimation of method precision. A target dilution factor (DF) of 8.10 x $10^5$ was planned per dilution series, so that the lowest dilution of each unit of SM4 could be analysed directly by RT-dPCR.

For SM2, SM3 and SM4 short-term stability was assessed by one-step RT-dPCR using the CDC N2 assay (Table B-3). In addition to RT-dPCR, the 6 undiluted STS units of SM4 were analysed on the Agilent Bioanalyser 2100 using an RNA 6000 Pico kit, according to the manufacturer's instructions. The STS samples (SM4) were analysed and transcript size compared to units stored at -80 °C (same units as for homogeneity testing).

***Long-term stability study***

Long-term stability of the materials was assessed 6 months to 7 months after the homogeneity and short-term stability studies. Four units of each of SM2, SM3 and SM4 were analysed following storage at the reference temperature of −80 °C. For stability testing of SM4, the units were diluted gravimetrically in 0.5 ng/µL Human Jurkat cell total RNA using an Ohaus E10640 balance to 4 decimal places. One dilution series per unit was included in the analysis. A target dilution factor (DF) of 8.10 x $10^5$ was planned per dilution series, so that the lowest dilution of each unit of SM4 could be analysed directly by RT-dPCR. Long-term stability was assessed by one-step RT-dPCR using the CDC N2 assay (Table B-3).

***Evaluation of assay uncertainty***

Assay uncertainty was estimated by measurement of an *in vitro* transcribed RNA material containing partial *RdRp* gene, *E* and *N* genes (full length) on the same construct. US CDC N2, N3, Sarbeco E assays and three inhouse designed assays (two designed to *N*, one to *E*) were tested in three independent experiments ($n$ = 4 per assay) with average λ values of 0.4. Results from one of the inhouse assays which led to reduced concentration results were interpreted as showing a negative bias and not included in combined analysis. Results from the five remaining comparable assays were analysed by linear mixed effects model using R software (7, 8), which estimated a between-assay relative SD of 4 %.

#### *Statistical analysis of study results*

Graphpad Prism version 9 (Graphpad, San Diego, CA) was used for calculation of descriptive statistics of study results, comparison of RT-dPCR assays (one-way ANOVA and unpaired *t*-test, assuming equal variances of groups).

R version 3.6.1 running inside RStudio version 1.2.5001 was used for normality testing (Shapiro Wilk test) and comparison of Study Material 3 vs. Study Material 4 extrapolated results by Welch Two Sample t-test and Wilcoxon rank sum test. Pairwise evaluation of result consistency and calculation of DerSimonian-Laird and Mandel-Paul RV estimators were performed using the metrology R package (https://cran.r-project.org/package=metRology). The Huber Proposal 2 estimator was calculated using the MASS R package (https://cran.r-project.org/package=MASS). Mean/SD and Median/MADe estimators were calculated using standard R statistical functions.

### Coordinating laboratory methodology NIST

#### *Gravimetry*

NIST used a Mettler Toledo AB54 balance to record the gravimetric diltions for M4. Following a protocol by NML, NIST made sequential dilutions of M4 on three separate days (Table B8). The diluent was RSS with 0.5 ng/µL Jurkat RNA. Both M4 and the diluent were allowed to reach room temperature before the gravimetric measurements began.

#### *One-step RT-dPCR methodology*

NIST used the Bio-Rad One Step RT-ddPCR kit along with the manual droplet generator with measure the material with the N2 and N3 CDC assays. 5 µL of the diluted material (either D4 or D5) was added to a 20 µL total volume reaction. The RT-dPCR reaction conditions were: 250 nM for gene primers, 250 nM for probe; Stage 1= 60 min at 50 °C, Stage 2=10 min at 95 °C, 60 cycles of Stage 3, Step 1, 30 sec at 95 °C, then Step 2 = 1 min at 55 °C, Stage 4= 10 min at 98 °C, Stage 5= Hold 4 °C. After thermocycling was completed, the plate was read on a QX200.

Table B-8: Gravimetric preparation

Table B-8: Part A: Unit No. 002

| Sample | Tare (g) | Mass of tube (g) | Mass of tube (g) minus tare | Tare (g) | Mass of tube + DILUENT (g) | Mass of tube + DILUENT minus tare(g) | Mass of DILUENT (g) | Tare (g) | Mass of tube + diluent + sample (g) | Mass of tube + diluent + sample minus tare (g) | Mass of SAMPLE (g) | Volumetric DF | Gravimetric DF |
|---|---|---|---|---|---|---|---|---|---|---|---|---|---|
| D1 | 0.00000 | 1.01750 | 1.01750 | 0.00000 | 1.45110 | 1.45110 | 0.43360 | 0.00000 | 1.46590 | 1.46590 | 0.01480 | 30.00000 | 30.29730 |
| D2 | 0.00000 | 1.01240 | 1.01240 | 0.00000 | 1.44770 | 1.44770 | 0.43530 | 0.00000 | 1.46250 | 1.46250 | 0.01480 | 30.00000 | 30.41216 |
| D3 | 0.00000 | 1.01070 | 1.01070 | 0.00000 | 1.44590 | 1.44590 | 0.43520 | 0.00000 | 1.46080 | 1.46080 | 0.01490 | 30.00000 | 30.20805 |
| D4 | 0.00000 | 1.02390 | 1.02390 | 0.00000 | 1.45820 | 1.45820 | 0.43430 | 0.00000 | 1.47310 | 1.47310 | 0.01490 | 30.00000 | 30.14765 |
| | | | | | | | | | | | Total | 8.10E+05 | 8.39E+05 |

Table B-8: Part B: Unit No. 023

| Sample | Tare (g) | Mass of tube (g) | Mass of tube (g) minus tare | Tare (g) | Mass of tube + DILUENT (g) | Mass of tube + DILUENT minus tare(g) | Mass of DILUENT (g) | Tare (g) | Mass of tube + diluent + sample (g) | Mass of tube + diluent + sample minus tare (g) | Mass of SAMPLE (g) | Volumetric DF | Gravimetric DF |
|---|---|---|---|---|---|---|---|---|---|---|---|---|---|
| D1 | 0.00000 | 1.00800 | 1.00800 | 0.00000 | 1.44230 | 1.44230 | 0.43430 | 0.00000 | 1.45720 | 1.45720 | 0.01490 | 30.00000 | 30.14765 |
| D2 | 0.00000 | 1.01550 | 1.01550 | 0.00000 | 1.45120 | 1.45120 | 0.43570 | 0.00000 | 1.46600 | 1.46600 | 0.01480 | 30.00000 | 30.43919 |
| D3 | 0.00000 | 1.01300 | 1.01300 | 0.00000 | 1.44940 | 1.44940 | 0.43640 | 0.00000 | 1.46440 | 1.46440 | 0.01500 | 30.00000 | 30.09333 |
| D4 | 0.00000 | 1.01690 | 1.01690 | 0.00000 | 1.45260 | 1.45260 | 0.43570 | 0.00000 | 1.46730 | 1.46730 | 0.01470 | 30.00000 | 30.63946 |
| | | | | | | | | | | | Total DF | 8.10E+05 | 8.46E+05 |

Table B-8: Part C: Unit No. 051

| Sample | Tare (g) | Mass of tube (g) | Mass of tube (g) minus tare | Tare (g) | Mass of tube + DILUENT (g) | Mass of tube + DILUENT minus tare(g) | Mass of DILUENT (g) | Tare (g) | Mass of tube + diluent + sample (g) | Mass of tube + diluent + sample minus tare (g) | Mass of SAMPLE (g) | Volumetric DF | Gravimetric DF |
|---|---|---|---|---|---|---|---|---|---|---|---|---|---|
| D1 | 0.00000 | 1.01390 | 1.01390 | 0.00000 | 1.45030 | 1.45030 | 0.43640 | 0.00000 | 1.46510 | 1.46510 | 0.01480 | 30.00000 | 30.48649 |
| D2 | 0.00000 | 1.01250 | 1.01250 | 0.00000 | 1.44770 | 1.44770 | 0.43520 | 0.00000 | 1.46250 | 1.46250 | 0.01480 | 30.00000 | 30.40541 |
| D3 | 0.00000 | 1.01040 | 1.01040 | 0.00000 | 1.44630 | 1.44630 | 0.43590 | 0.00000 | 1.46120 | 1.46120 | 0.01490 | 30.00000 | 30.25503 |
| D4 | 0.00000 | 1.02370 | 1.02370 | 0.00000 | 1.45970 | 1.45970 | 0.43600 | 0.00000 | 1.47460 | 1.47460 | 0.01490 | 30.00000 | 30.26174 |
| | | | | | | | | | | | Total DF | 8.10E+05 | 8.49E+05 |

### REFERENCES


1. Corman VM, Landt O, Kaiser M, Molenkamp R, Meijer A, Chu DKW, Bleicker T, Brünink S, Schneider J, Schmidt ML*, et al.* Detection of 2019 novel coronavirus (2019-nCoV) by real-time RT-PCR. Euro Surveill. 2020;25(3):2000045.
2. China CDC. New coronavirus nucleic acid detection primers and probe sequences (Specific primers and probes for detection 2019 novel coronavirus) 2020 [updated 21st January 2020]. Available from: http://ivdc.chinacdc.cn/kyjz/202001/t20200121_211337.html.
3. Ambion™ Life Technologies. MEGAscript® kit User Guide. 2012;Revision G.
4. Centers for Disease Control and Prevention (CDC). Research Use Only 2019-Novel Coronavirus (2019-nCoV) Real-time RT-PCR Primers and Probes 2020 [updated 6th June 2020]. Available from: https://www.cdc.gov/coronavirus/2019-ncov/lab/rt-pcr-panel-primer-probes.html.
5. Devonshire AS, Honeyborne I, Gutteridge A, Whale AS, Nixon G, Wilson P, Jones G, McHugh TD, Foy CA, Huggett JF. Highly reproducible absolute quantification of Mycobacterium tuberculosis complex by digital PCR. Anal Chem. 2015;87(7):3706-13.
6. Whale AS, Bushell C, Grant PR, Cowen S, Guttierrez-Aguirre I, O'Sullivan DM, Zel J, Milavec M, Foy CA, Nastouli E*, et al.* Detection of rare drug resistance mutations by digital PCR in a human influenza A virus model system and clinical samples. J Clin Microbiol. 2016;54(2):392-400.
7. R Core Team. R: A language and environment for statistical computing R Foundation for Statistical Computing, Vienna, Austria.2018 [Available from: https://www.R-project.org/].
8. Bates D MM, Bolker B, Walker S. Fitting Linear Mixed-Effects Models Using lme4. J Stat Softw. 2015;67(1):1–48.
