## Supplementary material for "CCQM-P199b: Interlaboratory comparability study of SARS-CoV-2 RNA copy number quantification": CCQM P199b supplementary file Appendix C

#### APPENDIX C: Protocol

See next page

### STUDY PROPOSAL AND PROTOCOL

### CCQM NAWG P199b: SARS-CoV-2 RNA copy number quantification

Study coordinators NML at LGC, NIMC, NIBSC and NIST

CCQM NAWG P199b will expand on capabilities demonstrated in the P199 study (HIV-1 RNA copy number quantification) for **targeted RNA copy number concentration** and **viral gene quantification** by measuring:

A. multiple different gene targets from different positions within the same viral genome (reflecting the variety of sequences measured by different diagnostic tests);
B. a different pathogen (to build evidence that the chosen reference measurement approaches are broadly applicable to a range of RNA sequence templates (multiple RNA viruses, prokaryotic and mammalian RNA molecules).

CCQM NAWG P199b will also establish candidate reference measurement procedures for detecting the viral genome of the causative agent of the Covid-19 pandemic (2019-nCoV, termed as SARS-CoV-2). This will expedite the ability of NMIs/DIs to demonstrate capability for reference measurement of SARS-CoV-2 Reference Materials (RMs), Quality Control (QC) materials and sources of whole viral genome materials enabling them to support diagnostic manufacturers, clinical laboratory-developed tests and international test standardization efforts. As with CCQM P199, the anticipated impact is standardisation of the performance of analytical methods used to diagnose Covid-19; all materials used in this study will be non-infectious purified RNA templates.

#### 1. Aim

The study rationale is to support higher order measurement of RNA copy number concentration of specific genetic sequences present in the SARS-CoV-2 RNA genome.

The specific aims are:

1. To measure the RNA copy number concentration of the nucleocapsid (*N*) gene (Study Materials 1-3), and Envelope (*E*) gene targets (Study Materials 1 and 2), in the 'low' copy number concentration range ($10^1$-$10^4$ copies/µL).
2. To evaluate the intra- and inter-laboratory variability in results reported between laboratories in the measurement range $10^1$ to $10^4$ copies/µl.
3. To measure the copy number concentration of the *N* gene RNA sequence fragment in the pure 'high' copy number concentration range ($10^9$-$10^{10}$ copies/µL) (Study Material 4).
4. To assess the trueness of the methods used for Study Materials 1, 2 and 3 with orthogonal methods (such as mass-spectrometry or flow cytometric single molecule

counting) by comparing measurements of high and low concentration materials linked by gravimetric dilution (Study Materials 3 and 4).

5. To compare *N* gene target measurements from more simple and complex RNA templates (*in vitro* transcribed RNA single fragments and more complex mixed larger viral fragments (simulated viral genomic RNA (Study Material 2)).
6. To provide evidence for CMC claims by participating laboratories when measuring purified RNA to consider multiple gene targets from different RNA viruses.

#### 2. Background

This study is proposed to apply the aims and approach of study "CCQM-P154 Absolute Quantification of DNA" to 'absolute' quantification of RNA and will follow on from the P199 study that is due to be reported in May 2020.

This study will be fast tracked to provide NMIs with the tools to be able to respond to national needs in the global response to the SARS-CoV-2 pandemic.

The aim of CCQM-P154 was to assess the quantification of low-level amounts of DNA in an absolute manner without the aid of calibration using enumeration-based techniques (eight laboratories performed dPCR with one laboratory using flow cytometric single molecule counting). The results reported for the low level 'Level II' material (consensus value 7970 copies/mg) were compared to values reported by laboratories using orthogonal methods (ID-MS, UV-CE) for the approximately 100,000 times more concentrated 'Level I' material from which the Level II material was prepared. By calculating the gravimetric dilution factors between the Level I and II materials, a direct comparison of the newer methods was possible with the established SI traceable methods used to measure the Level I material (Yoo *et al.*, 2016) . The close agreement between the mean results of the four orthogonal approaches tested (CV 1.8%) strongly supported the accuracy of more recently developed enumeration-based techniques.

Previous CCQM pilot studies have demonstrated NMI capabilities to perform accurate measurements of RNA copy number concentration (P103 "Measurement of Multiplexed Biomarker Panel of RNA Transcripts") and RNA copy number ratio (P103.1: "Multiple cancer cell biomarker measurement"). Reverse transcription-digital PCR (RT-dPCR) was used by sub-set of laboratories in these studies. Reported values for copy number concentration using RT-dPCR were within 1.5-fold (P103) and 1.4-fold (P103.1) of those assigned on the basis of molecule weight and UV spectroscopy. In CCQM P155 "Multiple cancer cell biomarker measurement", RT-dPCR was used to value assign the copy number concentration of three *in vitro* transcribed transcripts in the study calibration material which enable reporting of mRNA copy number concentration for the two cellular study materials.

Published studies have demonstrated the utility of 'absolute quantification' of mRNA and viral RNA in a number of applications using methods like RT-dPCR (Chen *et al.*, 2013, Whale *et al.*, 2016, White *et al.*, 2012). However error in reverse transcription, currently a necessary step for

RNA measurement using molecular biology methodologies, has also been shown to significantly impact dPCR measurements of mRNA (Sanders *et al*., 2013) and miRNA (Stein *et al*., 2017).

In P199, HIV-1 was chosen as a model as it is a pathogen of major global importance which is quantified clinically to guide treatment and monitor resistance, and is reported as an 'absolute' concentration (such as copies/mL plasma). This differs from other RNA targets measured clinically, which are reported as a copy number ratio, such as the transcript of the *BCR-ABL* fusion gene. The P199 study data is being compiled and will be reported shortly.

For the present study, SARS-CoV-2 will be the chosen model. Current molecular diagnostic approaches mainly apply reverse transcription real time quantitative PCR (RT-qPCR) to measure the viral RNA as a Laboratory Developed Test (**Table 1**) with an increasing number of commercial tests coming on line (FIND, 2020).

**Table 1. Molecular assays developed by public health laboratories[1] for diagnosis of COVID-19.**

| Country | Institute | Genetic target(s) | Genome position(s) (NC_045512.2) |
|---|---|---|---|
| CN | Chinese Center for Disease Control and Prevention | *ORF*1ab | 13342-13460 |
| | | *N* | 28881-28979 |
| DE, NL, UK | Charité, Tib-Molbiol, Erasmus MC and Public Health England | *RdRP** | 15431-15530 |
| | | *E* | 26269-26381 |
| HK | School of Public Health, The University of Hong Kong | ORF1b-nsp14 | 18778-18909 |
| | | N | 29145-29254 |
| JP | National Institute of Infectious Diseases | *ORF*1a | 484-896; 492-837 |
| | | *S* | 24354-24900; 24364-24856 |
| | | *N* | 29125-29282 |
| TH | Ministry of Public Health | *N* | 28320-28376 |
| USA | Centers for Disease Control and Prevention | *N* | 28287-28358; 29164-29230;28681-28752 |
| | | Human *RNASE P* mRNA (positive control target) | N/A |
| FR | Institut Pasteur, Paris | *RdRP** | 12690-12797; 14080-14186 |
| | | *E*** | 26269-26381 |

[1]Coronavirus disease (COVID-19) technical guidance: Laboratory testing for 2019-nCoV in humans: In-house developed molecular assays https://www.who.int/emergencies/diseases/novel-coronavirus-2019/technical-guidance/laboratory-guidance). *E* – envelope, *N* – nucleocapsid, *nsp* - non-structural protein, *ORF* - open reading frame, *RdRp* - RNA dependent RNA polymerase, *RNase* P - Ribonuclease P, *S* – spike protein. **RdRp* gene is within the *ORF*1ab sequence. **Same assay to *E* gene as DE/NL/UK assay.

The quantitative results of SARS-CoV-2 RNA load is not currently widely factored into clinical decisions, however variation in pre-analytical steps, such as specimen choice, specimen sampling, storage, extraction and nucleic acid pre-preparation (reverse transcription) can all impact on performance. Where suboptimal performance of such steps lead to false negative results, COVID-19 patients will be missed and may not be isolated; potentially spreading the infection further. RT-qPCR has already been shown to perform sub optimally, however, RT-dPCR formats have potentially provided greater confidence than RT-qPCR (Dong et al 2020, Suo et al 2020). Unfortunately, there are not enough dPCR instruments globally for such a technology to be used on a wider scale. However, the accuracy of dPCR can be harnessed to provide confidence in the value assignment of the variety of reference materials already available for SARS-CoV-2. Such materials could, in turn, provide greater confidence in the use of the established diagnostic approaches and improve their role in managing this pandemic.

This pilot study will aim to provide evidence of NAWG participants' capability in RT-dPCR measurements in support of global SARS-CoV-2 diagnostics.

#### 3. Study Materials

##### 3.1 Description

Four study materials have been designed and prepared by three different institutes: Study Material 1 (NIM China), Study Material 2 (NIBSC), Study Materials 3 and 4 (NML at LGC). All materials are synthetic or purified RNA and are non-infectious. Sequence information is provided in **Annex A** and is summarised in **Table 2**. It is intended that all study participants analyse Study Material 1, 2 and 3, whereas analysis of Study Materials 4 is optional. Study participants will be provided with four units of each study material.

**Table 2: Summary of Study Material genomic information**

| Study Material | Gene construct name (GenBank accession) | SARS-CoV-2 genome region (NC_045512.2) | Gene targets included |
|---|---|---|---|
| 1 | ORF1ab | 13201-15600 | *ORF1ab* (partial) |
| | E | 26245-26472 | *E* |
| | N | 28274-29533 | *N* |
| 2 | Construct 1 (MT299802) | 1-7515 | *ORF1ab* (partial) |
| | Construct 2 (MT299803) | 7416-14915 | *ORF1ab* (partial) |
| | Construct 3 (MT299804) | 14816-22421 | *ORF1ab* (partial) |
| | Construct 4 (MT299805) | 22322-29903 | *E* and *N* |
| 3 | N | 28274-29239 | *N (partial)* |
| 4 | N | 28274-29239 | *N (partial)* |

**Study Material 1** contains *in vitro* transcribed SARS-CoV-2 RNA fragments of ORF1ab, full length *E* gene and full length *N* gene, at an approximate concentration ~$10^1$-$10^4$ copies/µl in ~5 ng/µL 293T human cell line total RNA in buffered solution (1 mM sodium citrate, pH 6.5 (RNA Storage Solution Thermo Fisher Scientific P/N AM7001)). A total of 200 units, each containing 100 µL, were prepared.

**Study Material 2** is composed of purified RNA from four lentiviral constructs (MT299802, MT299803, MT299804, MT299805). Construct 4 (MT299805) contains the SARS

-CoV-2 genomic region corresponding to NC_045512.2: 22322-29903 which includes the *E* and *N* genes. The SARS-CoV-2 genome has been divided into four overlapping fragments and inserted within the long terminal repeats of a lentiviral vector (LVV) plasmid. Single nucleotide mutations have been randomly inserted in the SARS-CoV-2 sequences to prevent protein expression. Please note that the two base substitutions in the *E* gene (**Annex A, Table A2**) occur within the primers for the "Sarbeco" *E* gene assay developed by Charité – Universitätsmedizin Berlin (Corman *et al*., 2020) (**Tables 1 and 3**).

The four LVV-SARS-CoV-2 constructs were each individually transfected into HEK293T/17 cells together with a HIV-1 packaging plasmid and subsequently pooled as detailed in the Data Sheet Research Reagent for SARS-CoV-2 RNA NIBSC code 19/304 (https://www.nibsc.org/documents/ifu/19-304.pdf). The material contains a similar RNA copy number concentration to Study Material 1. RNA was purified using the QIAamp® UltraSens® Virus (Qiagen) and diluted in ~5 ng/µl FirstChoice® Human T-Cell Leukemia (Jurkat) Total RNA (Ambion P/N AM7858) in RNA Storage Solution (as above). Each unit of material contains 100 µL sample. A total of 300 units were prepared.

**Study Material 3** contains *in vitro* transcribed SARS-CoV-2 RNA fragment of the partial *N* gene in a similar RNA copy number concentration to Study Material 1, in ~2.5 ng/µL human Jurkat cell line total RNA (as above) in RNA Storage Solution (as above) and was prepared by gravimetric dilution of Study Material 4. A total of 300 units, each containing 100 µL, were prepared.

**Study Material 4** contains the same *in vitro* transcribed SARS-CoV-2 RNA fragment of the Nucleocapsid gene as Study Material 3 at an approximate concentration $10^9$-$10^{10}$ copies/µl (~0.5-5 ng/µl) in RNA Storage Solution (as above) in a volume of 100 µl per unit. A total of ~60 units were prepared. No additional RNA molecules have been added to this material and it is designed to be suitable for analysis by chemical analysis methods and single molecule flow cytometry.

##### 3.2 Homogeneity

The homogeneity of all study materials will be evaluated by RT-dPCR analysis (BioRad QX200). The homogeneity of Study Materials 1 - 3 will be assessed by performing replicate measurements (sub-samplings) of 10 units. Triplicate measurements of the *E* and *N* genes will be performed for Study Materials 1-2. Eight replicate measurements of the *N* gene will be performed for Study Material 3.

Due to the smaller number of units of Material 4 produced, homogeneity will be evaluated in ~10% of units in accordance with ISO Guide 35. Six units of Study Material 4 will be assessed by performing triplicate volumetric dilutions with triplicate RT-dPCR measurements of each dilution. The homogeneity of Study Material 4 will also be analysed by fluorimetric assay (Qubit RNA HS Assay, Invitrogen/Thermo Fisher Scientific) with 6 units ($n$ = 3 assays).

##### 3.3 Stability

A short term stability study will be performed by incubation of Study Materials 1-3 on dry ice, 4°C and 27°C for 3 and 7 days (and 14 days, Study Material 1 only) and compared to reference temperature (-80°C) ($n$ = 3 units per condition). Due to the limitation in unit number, two units of Study Material 4 will be placed dry ice, 4°C or 27°C for 7 days and compared to the reference temperature.

Stability for all Study Materials will be assessed by RT-dPCR ($n$ = 3) using a duplex assay to *Orf1ab*/*E* (Study Material 1 (NIMC)) and an assay to the *N* gene (Study Materials 2-4; NML). Study Material 4 will be measured following dilution and will also be measured using the Agilent 2100 Bioanalyzer (for peak size/area at the expected size of 969 bp).

A long term stability study will be commenced prior to shipment of study materials, with analysis of four units of each material performed every three months until completion of the study, with the final analysis being performed after submission of results. Only units stored at the reference temperature of −80° C will be analysed as this is the storage temperature for RNA samples which is recommended to participants.

#### 4. Measurands

##### 4.1 Measurand 1 (All Study Materials)

RNA copy number concentration expressed in copies per µL (c/µL) of the *N* gene partial sequence (NC_045512.2: 28274-29239) which is common to all Study Materials (**Table 2**).

<u>Analysis of Study Material 4 is optional for participants</u>.

##### 4.2 Measurand 2 (Study Materials 1-2 only)

RNA copy number concentration expressed in copies per µL (c/µL) of the *E* full gene sequence (NC_045512.2: 26245-26472) which is present only in Study Materials 1 and 2 (**Table 2**).

<u>Measurand 2 is optional for participants</u>.

#### 5. Shipping and storage of study materials

Study materials, primers and probes will be shipped to participants on dry ice:

- Study Material 1 (and primers and probes, if required, see below) will be sent by NIM China
- Study Materials 2, 3 and 4 will be sent by NML

If the materials have thawed at the point of receipt please request a fresh shipment to be sent.

It is recommended to store the study materials at -80 °C on receipt until analysis. When study materials are to be analysed, they should be thawed, vortexed to briefly to homogenise the contents and kept on ice. It is recommended that study materials do not undergo more than three freeze-thaw cycles.

Primers and probes should be stored at -20 °C.

#### 6. PCR assays and recommended probe types for use with One-Step RT-ddPCR Advanced Kit for Probes

Participants are encourage to design or select their own assays to the target sequences defined for the Measurands (Appendix A).

For those who have problems ordering primers and probes, assays to the *N* and *E* genes based on those developed by the Chinese Center for Disease Control and Prevention (*N* gene) and Charite/PHE (*E* gene) (as per **Table 1**) can be provided on request for this study along with protocols for QX200 (Bio-Rad) (**Table 3**).

**Table 3: Optional Assays for RT-dPCR analysis**

| Gene | Primer/probe | Sequence (5'-3') |
|---|---|---|
| *N* | Forward primer | GGG GAA CTT CTC CTG CTA GAA T |
| | Reverse primer | CAG ACA TTT TGC TCT CAA GCT G |
| | Probe | FAM-TTG CTG CTG CTT GAC AGA TT-BHQ1 |
| *E* | Forward primer | ACA GGT ACG TTA ATA GTT AAT AGC GT |
| | Reverse primer | ATA TTG CAG CAG TAC GCA CAC A |
| | Probe | FAM-ACA CTA GCC ATC CTT ACT GCG CTT CG-BHQ1 |

As noted for CCQM P199, for participants who adopt a one-step RT-dPCR approach using the QX100 or QX200, it is recommended that participants check the compatibility of fluorescent probe types with One-Step RT-ddPCR Advanced Kit for Probes (Bio-Rad P/N 1864021 and 1864022). Double-quenched probes (such as ZEN probes (IDT)) and Minor Groove Binding Non-Fluorescent Quencher (MGB-NFQ) probes (Thermo Fisher Scientific) have been shown to give better separation of negative and positive droplets and lower background of negative droplets compared to single-quenched probes (Maier *et al*., 2019, Pinheiro-de-Oliveira *et al*., 2019; NML at LGC personal communication regarding BHQnova).

#### 7. Schedule*

| Date | Timeline | Actions (Participants) |
|---|---|---|
| 05 May 2020 | Protocol V1.0 circulated | Coordinators |
| 12 May 2020 | Registration for participation | Reply with Form 1 (Participant) |
| 18 May to 05 June 2020 | Study Material distribution | Reply with Form 2 (Participant) |
| 05 June 2020 | Protocol update to be circulated (Results of homogeneity and stability studies) | Coordinators |
| 14 September 2020 | Submission of results | Data return with Forms 3-4 (Participant) |
| October 2020 | Initial report at NAWG | Presentation of results |

*Timeline dates may be subject to disruption linked to current global situation associated with COVID-19. This may impact on study aspects such as shipping, material receipt and submission of results. This will be accommodated.

#### 8. Expected outcomes of the study

The study aims to support competencies in targeted measurement of RNA-based materials expressed as copy number concentration.

1. Evidence for CMC claims for measurement of RNA copy number concentration of *in vitro* synthetised RNA templates in a complex matrix (human total RNA) in the concentration range ~$10^1$-$10^4$ copies/µL (Study Materials 1, 3). The current study extends claims based on P199 to demonstrate the suitability of RMPs by measuring additional gene targets increase the measurement challenge due to factors such as variable GC content, sequence context and secondary structure.
2. Information on the appropriate **measurement uncertainties** to be applied to reference measurement procedures for viral RNA copy number quantification using RT-dPCR through evaluation of the intra- and inter-laboratory variability in results reported between laboratories in the measurement range $10^1$ to $10^4$ copies/µl.
3. Evidence for CMC claims of high concentration RNA materials ($10^9$-$10^{10}$ copies/µL) containing a single template molecule in buffered aqueous solution.
4. Information on biases affecting RT-dPCR methods through the comparison of results for high and low concentration materials linked by gravimetric dilution (Study Materials 3 and 4). This information, together with results of P199, will guide the development of SI-traceable calibration hierarchies for RNA copy number concentration.
5. Analysis of Study Material 2 would extend the claim related to Study Materials 1 and 3 to capability in analysis of more complex, simulated viral genomic RNA templates which have been purified from viral particles produced in cell culture, where RNA fragment length may be more variable due to the purification process.
6. Evidence for the fitness-for-purpose of RT-dPCR based reference measurement procedures in calibration and QC of RT-qPCR and other diagnostic end-user approaches.

Due to the Study Materials being purified RNA samples, the study is not designed to assess the influence of matrix (inhibitors etc) or pre-analytical factors, such as extraction, on RNA measurements. However demonstrating the competencies examined by this study would place NMIs and DIs in the ideal position to support investigation of these key additional considerations.

#### 9. Reporting and publication of Results

*For discussion CCQM NAWG May 2020*

#### 10. References

Chen WW, Balaj L, Liau LM, Samuels ML, Kotsopoulos SK, Maguire CA, *et al*. Beaming and droplet digital PCR analysis of mutant *IDH1* mrna in glioma patient serum and cerebrospinal fluid extracellular vesicles. *Mol. Ther. Nucleic Acids* 2013;2:e109.

Corman VM, Landt O, Kaiser M, Molenkamp R, Meijer A, Chu DK *et al*. Detection of 2019 novel coronavirus (2019-nCoV) by real-time RT-PCR. *Euro Surveill*. 2020;25(3). doi: 10.2807/1560-7917.ES.2020.25.3.2000045

Dong L, Zhou J, Niu C, Wang Q, Pan Y, Wang X, *et al*. Highly accurate and sensitive diagnostic detection of SARS-CoV-2 by digital PCR. *medRxiv* 2020. https://doi.org/10.1101/2020.03.14.20036129

FIND. SARS-CoV-2 Diagnostic Pipeline: Molecular Assays https://www.finddx.org/covid-19/pipeline/?section=molecular-assays#diag_tab (access date 27/04/2020)

Maier J, Lange T, Cross M, Wildenberger K, Niederwieser D, Franke GN. Optimized digital droplet PCR for *BCR-ABL*. *J. Mol. Diagn.*.2019;21:27-37.

Pinheiro-de-Oliveira TF, Fonseca-Junior AA, Camargos MF, Laguardia-Nascimento M, Giannattasio-Ferraz S, Cottorello ACP, *et al*. Reverse transcriptase droplet digital PCR to identify the emerging vesicular virus Senecavirus A in biological samples. Transbound Emerg Dis 2019;66:1360-9.

Sanders R, Mason DJ, Foy CA, Huggett JF. Evaluation of digital PCR for absolute RNA quantification. *PLoS One* 2013;8:e75296.

Stein EV, Duewer DL, Farkas N, Romsos EL, Wang L, Cole KD. Steps to achieve quantitative measurements of microRNA using two step droplet digital PCR. *PLoS One* 2017;12:e0188085.

Suo T, Liu Z, Guo M, Feng J, Hu W, Yang Y, *et al*. ddPCR: A more sensitive and accurate tool for SARS-CoV-2 detection in low viral load specimens. *medRxiv* 2020. https://doi.org/10.1101/2020.02.29.20029439

Whale AS, Bushell CA, Grant PR, Cowen S, Gutierrez-Aguirre I, O'Sullivan DM, *et al*. Detection of rare drug resistance mutations by digital PCR in a human influenza a virus model system and clinical samples. *J. Clin. Microbiol.* 2016;54:392-400.

White RA, 3rd, Quake SR, Curr K. Digital PCR provides absolute quantitation of viral load for an occult RNA virus. *J. Virol. Methods* 2012;179:45-50.

Yoo HB, Park SR, Dong L, Wang J, Sui Z, Pavsic J, *et al*. International comparison of enumeration-based quantification of DNA copy-concentration using flow cytometric counting and digital polymerase chain reaction. *Anal. Chem.* 2016;88:12169-76.

### Annex A: P199b Study Material Sequence information

#### Study Material 1

**Box A1** and **Box A2** show the relevant sequences in Material 1 for Measurands 1 and 2 respectively.

##### Box A1: Study Material 1 target sequence (Measurand 1)

```
>NC_045512.2:28274-29239 Severe acute respiratory syndrome coronavirus 2
isolate Wuhan-Hu-1, complete genome
ATGTCTGATAATGGACCCCAAAATCAGCGAAATGCACCCCGCATTACGTTTGGTGGACCCTCAGATTCAA
CTGGCAGTAACCAGAATGGAGAACGCAGTGGGGCGCGATCAAAACAACGTCGGCCCCAAGGTTTACCCAA
TAATACTGCGTCTTGGTTCACCGCTCTCACTCAACATGGCAAGGAAGACCTTAAATTCCCTCGAGGACAA
GGCGTTCCAATTAACACCAATAGCAGTCCAGATGACCAAATTGGCTACTACCGAAGAGCTACCAGACGAA
TTCGTGGTGGTGACGGTAAAATGAAAGATCTCAGTCCAAGATGGTATTTCTACTACCTAGGAACTGGGCC
AGAAGCTGGACTTCCCTATGGTGCTAACAAAGACGGCATCATATGGGTTGCAACTGAGGGAGCCTTGAAT
ACACCAAAAGATCACATTGGCACCCGCAATCCTGCTAACAATGCTGCAATCGTGCTACAACTTCCTCAAG
GAACAACATTGCCAAAAGGCTTCTACGCAGAAGGGAGCAGAGGCGGCAGTCAAGCCTCTTCTCGTTCCTC
ATCACGTAGTCGCAACAGTTCAAGAAATTCAACTCCAGGCAGCAGTAGGGGAACTTCTCCTGCTAGAATG
GCTGGCAATGGCGGTGATGCTGCTCTTGCTTTGCTGCTGCTTGACAGATTGAACCAGCTTGAGAGCAAAA
TGTCTGGTAAAGGCCAACAACAACAAGGCCAAACTGTCACTAAGAAATCTGCTGCTGAGGCTTCTAAGAA
GCCTCGGCAAAAACGTACTGCCACTAAAGCATACAATGTAACACAAGCTTTCGGCAGACGTGGTCCAGAA
CAAACCCAAGGAAATTTTGGGGACCAGGAACTAATCAGACAAGGAACTGATTACAAACATTGGCCGCAAA
TTGCACAATTTGCCCCCAGCGCTTCAGCGTTCTTCGGAATGTCGCGCATTGGCATG
```

##### Box A2: Study Material 1 target sequence (Measurand 2)

```
>NC_045512.2:26245-26472 Severe acute respiratory syndrome coronavirus 2
isolate Wuhan-Hu-1, complete genome
ATGTACTCATTCGTTTCGGAAGAGACAGGTACGTTAATAGTTAATAGCGTACTTCTTTTTCTTGCTTTCG
TGGTATTCTTGCTAGTTACACTAGCCATCCTTACTGCGCTTCGATTGTGTGCGTACTGCTGCAATATTGT
TAACGTGAGTCTTGTAAAACCTTCTTTTTACGTTTACTCTCGTGTTAAAAATCTGAATTCTTCTAGAGTT
CCTGATCTTCTGGTCTAA
```

### Study Material 2

**Boxes A3** and **A4** show the target sequences for Measurands 1 and 2. Single nucleotide mutations have been randomly inserted in the SARS-CoV-2 sequences to prevent protein expression. Single nucleotide mutations which occur within the Measurand 1 and 2 target sequences are listed in **Table 1** and highlighted in **Boxes A3-A4**.

**Box A3: Study Material 2 target sequence (Measurand 1).** Positions where the construct sequence differs from the reference sequence (NC_045512.2) are shown in red.

```
>MT299805.1:9051-10016 Cloning vector pSF_lenti_SARS-CoV-2_partial-
S/E/M/N, complete sequence
ATGTCTGATAATGGACCCCAAAATCAGCGAAATGCACCCCGCATTACGTTTGGTGGACCCTCAGATTCAA
CTGGCAGTAACCAGAATGGAGAACGCAGTGGGGCGCGATGAAAACAACGTCGGCCCCAAGGTTTACCCAA
TAATACTGCGTCTTGGTTGACCGCTCTCACTCAACATGGCAAGGAAGACCTTAAATTCCCTCGAGGACAA
GGCGTTCCAATTAACACCAATAGCAGTCCAGATGACCAAATTGGCTACTACCGAAGAGCTACCAGACGAA
TTCGTGGTGGTGACGGTAAAATGAAAGATCTCAGTCCAAGATGGTATTTCTACTACCTAGGAACTGGGCC
AGAAGCTGGACTTCCCTATGGTGCTAACAAAGACGGCATCATATGGGTTGCAACTGAGGGAGCCTTGAAT
ACACCAAAAGATCACATTGGCACCCGCAATCCTGCTAACAATGCTGCAATCGTGCTACAACTTCCTCAAG
GAACAACATTGCCATAAGGCTTCTACGCAGAAGGGAGCAGAGGCGGCAGTCAAGCCTCTTCTCGTTCCTC
ATCACGTAGTCGCAACAGTTCAAGAAATTCAACTCCAGGCAGCAGTAGGGGAACTTCTCCTGCTAGAATG
GCTGGCAATGGCGGTGATGCTGCTCTTGCTTTGCTGCTGCTTGACAGATTGAACCAGCTTGAGAGCAAAA
TGTCTGGTAAAGGCCAACAACAACAAGGCCAAACTGTCACTAAGAAATCTGCTGCTGAGGCTTCTAAGAA
GCCTCGGCAAAAACGTACTGCCACTAAAGCATACAATGTAACACAAGCTTTCGGCAGACGTGGTCCAGAA
CAAACCCAAGGAAATTTTGGGGACCAGGAACTAATCAGACAAGGAACTGATTACAAACATTGGCCGCAAA
TTGCACAATTTGCCCCCAGCGCTTCAGCGTTCTTCGGAATGTCGCGCATTGGCATG
```

**Box A4: Study Material 2 target sequence (Measurand 2).** Positions where the construct sequence differs from the reference sequence (NC_045512.2) are shown in red.

```
>MT299805.1:7022-7249 Cloning vector pSF_lenti_SARS-CoV-2_partial-
S/E/M/N, complete sequence
ATGTACTCATTCGTTTCGGAAGAGACAGGTACGTAAATAGTTAATAGCGTACTTCTTTTTCTTGCTTTCG
TGGTATTCTTGCTAGTTACACTAGCCATCCTTACTGCGCTTCGATTGTGTGCGTAGTGCTGCAATATTGT
TAACGTGAGTCTTGTAAAACCTTCTTTTTACGTTTACTCTCGTGTTAAAAATCTGAATTCTTCTAGAGTT
CCTGATCTTCTGGTCTAA
```

**Table A1: Study Material 2 base substitutions.**

| Measurand | NC_045512.2 position | Substitution (vs NC_045512.2) | MT299805 position | Measurand position |
|---|---|---|---|---|
| 1 | 28383 | C → G | 9160 | 110 |
| 1 | 28432 | C → G | 9209 | 159 |
| 1 | 28778 | A → T | 9555 | 505 |
| 2 | 26279 | T → A | 7056 | 35 |
| 2 | 26370 | C → G | 7147 | 126 |

### Study Materials 3 and 4

Study Materials 3 and 4 are composed of IVT RNA molecules at two concentrations in (i) human Jurkat cell line total RNA (Material 3) or (ii) buffered solution (Material 4). The IVT RNA is a partial sequence of the SARS-CoV-2 *N* gene (NC_045512.2 positions 28274-29239).

Nucleotides 1-3 (underlined) of the below sequence represent the G-terminal of the T7 promoter used for *in vitro* transcription of RNA. The total sequence length is 969 nucleotides.

**Box A5: Study Materials 3 / 4 target sequence (Measurand 1).** Underlined sequences correspond to the transcription initiation site of T7 RNA polymerase.

```
> NML_pEX-A128_SARS-CoV-2_partial-N_IVT
GGGATGTCTGATAATGGACCCCAAAATCAGCGAAATGCACCCCGCATTACGTTTGGTGGACCCTCAGATTCAA
CTGGCAGTAACCAGAATGGAGAACGCAGTGGGGCGCGATCAAAACAACGTCGGCCCCAAGGTTTACCCAATAA
TACTGCGTCTTGGTTCACCGCTCTCACTCAACATGGCAAGGAAGACCTTAAATTCCCTCGAGGACAAGGCGTT
CCAATTAACACCAATAGCAGTCCAGATGACCAAATTGGCTACTACCGAAGAGCTACCAGACGAATTCGTGGTG
GTGACGGTAAAATGAAAGATCTCAGTCCAAGATGGTATTTCTACTACCTAGGAACTGGGCCAGAAGCTGGACT
TCCCTATGGTGCTAACAAAGACGGCATCATATGGGTTGCAACTGAGGGAGCCTTGAATACACCAAAAGATCAC
ATTGGCACCCGCAATCCTGCTAACAATGCTGCAATCGTGCTACAACTTCCTCAAGGAACAACATTGCCAAAAG
GCTTCTACGCAGAAGGGAGCAGAGGCGGCAGTCAAGCCTCTTCTCGTTCCTCATCACGTAGTCGCAACAGTTC
AAGAAATTCAACTCCAGGCAGCAGTAGGGGAACTTCTCCTGCTAGAATGGCTGGCAATGGCGGTGATGCTGCT
CTTGCTTTGCTGCTGCTTGACAGATTGAACCAGCTTGAGAGCAAAATGTCTGGTAAAGGCCAACAACAACAAG
GCCAAACTGTCACTAAGAAATCTGCTGCTGAGGCTTCTAAGAAGCCTCGGCAAAAACGTACTGCCACTAAAGC
ATACAATGTAACACAAGCTTTCGGCAGACGTGGTCCAGAACAAACCCAAGGAAATTTTGGGGACCAGGAACTA
ATCAGACAAGGAACTGATTACAAACATTGGCCGCAAATTGCACAATTTGCCCCCAGCGCTTCAGCGTTCTTCG
GAATGTCGCGCATTGGCATG
```
