## Supplementary material for "CCQM-P199b: Interlaboratory comparability study of SARS-CoV-2 RNA copy number quantification": CCQM P199b supplementary file Appendix D

### APPENDIX D: Registration Form

#### Form 1: Confirmation of Participation

| |
|---|
| Organisation name |
| Main contact person |
| Email addresses to be included on P199b mailing list |
| Study Materials to be analysed by organisation (please tick): |
| Study Material 1 |
| Study Material 2 |
| Study Material 3 |
| Study Material 4 (optional) |
| Contact person and email address for sample shipment |
| Address for sample shipment |
| Contact telephone for sample shipment |
| Special considerations for import |
