## Supplementary material for "CCQM-P199b: Interlaboratory comparability study of SARS-CoV-2 RNA copy number quantification": CCQM P199b supplementary file Appendix F

### APPENDIX F: Result reporting

##### Form 3: Submission of Results

| |
|---|
| Organisation name |
| Contact person and email address |

| **MATERIAL** | **MEASURAND** (UNIT) |
|---|---|
| **STUDY MATERIAL 1** | **Measurand 1 (*N* gene)** (copies/μL) |
| Value (*x*) | |
| Standard uncertainty (*u*) | |
| Coverage factor (*k*) | |
| Expanded uncertainty (*U*) | |
| Relative expanded uncertainty (Rel *U*) | |
| | **Measurand 2 (*E* gene)** (copies/μL) |
| Value (*x*) | |
| Standard uncertainty (*u*) | |
| Coverage factor (*k*) | |
| Expanded uncertainty (*U*) | |
| Relative expanded uncertainty (Rel *U*) | |
| **STUDY MATERIAL 2** | **Measurand 1 (*N* gene)** (copies/μL) |
| Value (*x*) | |
| Standard uncertainty (*u*) | |
| Coverage factor (*k*) | |
| Expanded uncertainty (*U*) | |
| Relative expanded uncertainty (Rel *U*) | |
| | **Measurand 2 (*E* gene)** (copies/μL) |
| Value (*x*) | |
| Standard uncertainty (*u*) | |
| Coverage factor (*k*) | |
| Expanded uncertainty (*U*) | |
| Relative expanded uncertainty (Rel *U*) | |
| **STUDY MATERIAL 3** | **Measurand 1 (*N* gene)** (copies/μL) |
| Value (*x*) | |
| Standard uncertainty (*u*) | |
| Coverage factor (*k*) | |
| Expanded uncertainty (*U*) | |
| Relative expanded uncertainty (Rel *U*) | |

| **STUDY MATERIAL 4** | **Measurand 1 (*N* gene)** (copies/μL) |
|---|---|
| Value (*x*) | |
| Standard uncertainty (*u*) | |
| Coverage factor (*k*) | |
| Expanded uncertainty (*U*) | |
| Relative expanded uncertainty (Rel *U*) | |

#### Original submission (Laboratory 10)

| **MATERIAL** | **MEASURAND** (UNIT) |
|---|---|
| **STUDY MATERIAL 1** | **Measurand 1 (*N* gene)** (copies/μL) |
| Value (*x*) | 902,51 |
| Standard uncertainty (*u*) | 27,61 |
| Coverage factor (*k*) | 2,01 |
| Expanded uncertainty (*U*) | 55,51 |
| Relative expanded uncertainty (Rel *U*) | 6,15% |
| | **Measurand 2 (*E* gene)** (copies/μL) |
| Value (*x*) | 513,18 |
| Standard uncertainty (*u*) | 32,32 |
| Coverage factor (*k*) | 2,14 |
| Expanded uncertainty (*U*) | 69,32 |
| Relative expanded uncertainty (Rel *U*) | 13,51% |
| **STUDY MATERIAL 2** | **Measurand 1 (*N* gene)** (copies/μL) |
| Value (*x*) | 6,74 |
| Standard uncertainty (*u*) | 0,55 |
| Coverage factor (*k*) | 1,99 |
| Expanded uncertainty (*U*) | 1,10 |
| Relative expanded uncertainty (Rel *U*) | 16,26% |
| | **Measurand 2 (*E* gene)** (copies/μL) |
| Value (*x*) | 3,48 |
| Standard uncertainty (*u*) | 0,32 |
| Coverage factor (*k*) | 1,99 |
| Expanded uncertainty (*U*) | 0,64 |
| Relative expanded uncertainty (Rel *U*) | 18,44% |
| **STUDY MATERIAL 3** | **Measurand 1 (*N* gene)** (copies/μL) |
| Value (*x*) | 962,23 |
| Standard uncertainty (*u*) | 55,87 |
| Coverage factor (*k*) | 2,20 |
| Expanded uncertainty (*U*) | 122,97 |
| Relative expanded uncertainty (Rel *U*) | 12,78% |

| **STUDY MATERIAL 4** | **Measurand 1 (*N* gene)** (copies/μL) |
|---|---|
| Value (*x*) | - |
| Standard uncertainty (*u*) | - |
| Coverage factor (*k*) | - |
| Expanded uncertainty (*U*) | - |
| Relative expanded uncertainty (Rel *U*) | - |
