## Supplementary material for "CCQM-P199b: Interlaboratory comparability study of SARS-CoV-2 RNA copy number quantification": CCQM P199b supplementary file Appendix G

### APPENDIX G: Experimental details form

#### Form 4: Experimental details

**(1)** Experimental information**:** Please complete CCQM-P199b dMIQE form.

**(2)** Experimental Design: Please describe or show in diagrammatic form the experimental design which was applied.

**(3)** Measurement uncertainty. Please summarise the calculation approach used and list the factors which were included

#### Form 5: dMIQE checklist

See next page

| ITEM TO CHECK | PROVIDED | COMMENT |
|---|---|---|
| | **Y/N** | |
| **1. SPECIMEN** | | |
| Detailed description of specimen type and numbers | **N** | N/A |
| Sampling procedure (including time to storage) | **N** | N/A |
| Sample aliquotation, storage conditions and duration | **N** | N/A |
| **2. NUCLEIC ACID EXTRACTION** | | |
| Description of extraction method including amount of sample processed | **N** | N/A |
| Volume of solvent used to elute/resuspend extract | **N** | N/A |
| Number of extraction replicates | **N** | N/A |
| Extraction blanks included? | **N** | N/A |
| **3. NUCLEIC ACID ASSESSMENT AND STORAGE** | | |
| Method to evaluate quality of nucleic acids | **N** | N/A |
| Method to evaluate quantity of nucleic acids (including molecular weight and calculations when using mass) | **N** | N/A |
| Storage conditions: temperature, concentration, duration, buffer, aliquots | **Y** | |
| Clear description of dilution steps used to prepare working DNA solution | **Y** | |
| **4. NUCLEIC ACID MODIFICATION** | | |
| Template modification (digestion, sonication, pre-amplification, bisulphite etc.) | **Y** | *If applicable* |
| Details of repurification following modification if performed | **Y** | *If applicable* |
| **5. REVERSE TRANSCRIPTION** | | |
| cDNA priming method and concentration | **Y** | *For 2-step RT-dPCR only* |
| One or two step protocol (include reaction details for two step) | **Y** | |
| Amount of RNA added per reaction | **Y** | *For 2-step RT-dPCR only (for 1-step RT-dPCR, record in Row 40)* |

| | | |
|---|---|---|
| Detailed reaction components and conditions | **Y** | *For 2-step RT-dPCR only (for 1-step RT-dPCR, record Section "dPCR Protocol")* |
| Estimated copies measured with and without addition of RT* | **Y** | |
| Manufacturer of reagents used with catalogue and lot numbers | **Y** | *For 2-step RT-dPCR only (for 1-step RT-dPCR, record Section "dPCR Protocol")* |
| Storage of cDNA: temperature, concentration, duration, buffer and aliquots | **Y** | *For 2-step RT-dPCR only* |
| **6. dPCR OLIGONUCLEOTIDES DESIGN AND TARGET INFORMATION** | | |
| Sequence accession number or official gene symbol | **Y** | |
| Method (software) used for design and *in silico* verification | **Y** | |
| Location of amplicon | **Y** | |
| Amplicon length | **Y** | |
| Primer and probe sequences (or amplicon context sequence)** | **Y** | |
| Location and identity of any modifications | **Y** | |
| Manufacturer of oligonucleotides | **Y** | |
| **7. dPCR PROTOCOL** | | |
| Manufacturer of dPCR instrument and instrument model | **Y** | |
| Buffer/kit manufacturer with catalogue and lot number | **Y** | |
| Primer and probe concentration | **Y** | |
| Pre-reaction volume and composition (incl. amount of template and if restriction enzyme added) | **Y** | |
| Template treatment (initial heating or chemical denaturation) | **Y** | |
| Polymerase identity and concentration, Mg++ and dNTP concentrations*** | **Y** | |
| Complete thermocycling parameters | **Y** | |
| **8. ASSAY VALIDATION** | | |
| Details of optimisation performed | **Y** | |

| | | |
|---|---|---|
| Analytical specificity (vs. related sequences) and limit of blank (LOB) | **Y** | *if available* |
| Analytical sensitivity/LoD and how this was evaluated | **Y** | *if available* |
| Testing for inhibitors (from biological matrix/extraction) | **N** | CCQM P199b Study Materials free from inhibitors |
| **9. DATA ANALYSIS** | | |
| Description of dPCR experimental design | **Y** | |
| Comprehensive details negative and positive of controls (whether applied for QC or for estimation of error) | **Y** | |
| Partition classification method (thresholding) | **Y** | |
| Examples of positive and negative experimental results (including fluorescence plots in supplemental material) | **Y** | |
| Description of technical replication | **Y** | |
| Repeatability (intra-experiment variation) | **Y** | *if available* |
| Reproducibility (inter-experiment/user/lab etc. variation ) | **Y** | *if available* |
| Number of partitions measured (average and standard deviation ) | **Y** | |
| Partition volume | **Y** | *Partition volume and measurement uncertainty* |
| Copies per partition (λ or equivalent ) (average and standard deviation) | **Y** | *For each Study Material analysed* |
| dPCR analysis program (source, version) | **Y** | |
| Description of normalisation method | **N** | N/A |
| Statistical methods used for analysis | **Y** | |
| Data transparency | | *Data to be requested by P199b coordinators if required for follow-up* |

**Table S1.** dMIQE2020 checklist for authors, reviewers and editors. Authors should fill detail whether information is provided. Where 'yes' is selected use comment box to detail location of information or to include the information. Where 'no' is selected use comment box to outline rationale for omission. Sections 4 and 5 may not apply depending on experiment.

* Assessing the absence of DNA using a no RT assay (or where RT has been inactivated) is essential when first extracting RNA. Once the sample has been validated as DNA-free, inclusion of a no-RT control is desirable, but no longer essential.

** Disclosure of the primer and probe sequence is highly desirable and strongly encouraged. However, since not all commercial pre-designed assay vendors provide this information when it is not available assay context sequences must be submitted (Bustin et al. Primer sequence disclosure: A clarification of the miqe guidelines. Clin Chem 2011;57:919-21.)

*** Details of reaction components is highly desirable, however not always possible for commercial disclosure reasons. Inclusion of catalogue number is essential where component reagent details are not available.
