## Supplementary material for "CCQM-P199b: Interlaboratory comparability study of SARS-CoV-2 RNA copy number quantification": CCQM P199b supplementary file Appendix H

### APPENDIX H: Summary of Participants' Analytical Information

The following Tables summarize the detailed information about the analytical procedures each participant provided in their Study Forms 4-5 (Appendix G). The presentation of the information in many entries has been consolidated and standardized.

A summary of the participants' measurement uncertainty information is provided in Appendix I.

Table H-1: Summary of Analytical Techniques for CCQM P199b Study Materials 1-3

| Laboratory ID - Institute code | Analytical Technique | One- or two-step RT-PCR (RT priming approach, two-step only) | RT reagent (two-step only) | RT-PCR reagent (one-step) or PCR reagent (two-step) | dPCR/qPCR instrument | Thermal Cycler (for two-step, state if same or different cyclers used RT and dPCR) |
|---|---|---|---|---|---|---|
| 1 | RT-dPCR | One | n/a | One-Step RT-ddPCR Advanced Kit for Probes (Bio-Rad) #1864021 | QX200 | C1000 |
| 2 | RT-dPCR | One | n/a | One-Step RT-ddPCR Advanced Kit for Probes (Bio-Rad) #1864021 | QX200 | C1000 |
| 3 | RT-dPCR | One | n/a | One-Step RT-ddPCR Advanced Kit for Probes (Bio-Rad) #1864021 | QX100 | C1000 |
| 4 | RT-dPCR | One | n/a | One-Step RT-ddPCR Advanced Kit for Probes (Bio-Rad) #1864021 | QX200 | C1000 |
| 5 | RT-dPCR | One | n/a | One-Step RT-ddPCR Advanced Kit for Probes (Bio-Rad) #1864021 | QX100 | C1000 |
| 6 | RT-dPCR | One | n/a | One-Step RT-ddPCR Advanced Kit for Probes (Bio-Rad) #1864022 | QX200 | C1000 |
| 7 | RT-dPCR | Two (gene-specific) | SuperScript IV First-Strand Synthesis System and SuperScript IV Reverse Transcriptase | QuantStudio 3D Digital PCR Master Mix v2 (A26358, Lot#: 2160352) | QS3D | TaKaRa PCR Thermal Cycler, TaKaRa QuantStudio 3D Digital PCR System, Thermo Fisher Scientific |

| Laboratory ID - Institute code | Analytical Technique | One- or two-step RT-PCR (RT priming approach, two-step only) | RT reagent (two-step only) | RT-PCR reagent (one-step) or PCR reagent (two-step) | dPCR/qPCR instrument | Thermal Cycler (for two-step, state if same or different cyclers used RT and dPCR) |
|---|---|---|---|---|---|---|
| 8 | RT-dPCR | Two (random hexamers) | SensiFAST cDNA Synthesis Kit, BIO-65054 | ddPCR Supermix for probes- Bio-Rad # 1863026 | QX200 | Verity, Thermofisher |
| 9 | RT-dPCR | One | n/a | One-Step RT-ddPCR Advanced Kit for Probes (Bio-Rad) #1864021 | QX200 | Thermofisher ProFlex |
| 10 | RT-dPCR | One | n/a | One-Step RT-ddPCR Advanced Kit for Probes (Bio-Rad) #1864022 | QX200 | C1000 |
| 11 | RT-dPCR | Two (gene-specific) | "DNA-Technology" kit (RevertAid reverse transcriptase, ThermoFisher) | ddPCR Supermix for probes (no dUTP) #1863025 | QX200 | Cycler for RT: CFX-96 (Bio-Rad)<br>Cycler for PCR: CFX-96 (Bio-Rad) |
| 12 | RT-dPCR | One | n/a | One-Step RT-ddPCR Advanced Kit for Probes (Bio-Rad) #1864021 | QX200 | C1000 |
| 13 | RT-dPCR | One | n/a | One-Step RT-ddPCR Advanced Kit for Probes (Bio-Rad) #1864022 | QX200 | C1000 |
| 14 | RT-dPCR | One | n/a | One-Step RT-ddPCR Advanced Kit for Probes (Bio-Rad) #1864021 | QX200 | C1000 |
| 15 | RT-dPCR | One | n/a | One-Step RT-ddPCR Advanced Kit for Probes (Bio-Rad) #1864021 | QX200 | C1000 |
| 16 | RT-dPCR | One | n/a | One-Step RT-ddPCR Advanced Kit for Probes (Bio-Rad) #1864021 | QX200 | GeneAmp PCR 9700, Applied Biosystems |
| 17 | RT-dPCR | One | n/a | One-Step RT-ddPCR Advanced Kit for Probes (Bio-Rad) #1864021 | QX200 | C1000 |
| 18 | RT-dPCR | One | n/a | One-Step RT-ddPCR Advanced Kit for Probes (Bio-Rad) #1864021 | QX200 | C1000 |
| 19 | RT-dPCR | One | n/a | One-Step RT-ddPCR Advanced Kit for Probes (Bio-Rad) #1864021 | QX200 | C1000 |
| 20 | RT-dPCR | One | N/A | One-step: One-Step RT-ddPCR Advanced Kit for Probes (Bio-Rad) #1864021 | QX200 | C1000 |

| Laboratory ID - Institute code | Analytical Technique | One- or two-step RT-PCR (RT priming approach, two-step only) | RT reagent (two-step only) | RT-PCR reagent (one-step) or PCR reagent (two-step) | dPCR/qPCR instrument | Thermal Cycler (for two-step, state if same or different cyclers used RT and dPCR) |
|---|---|---|---|---|---|---|
| 21 | RT-qPCR | One | n/a | Measurand 1: SARS-CoV-2 RNA detection system (Cat. # CDS-003N-200, Biolabmix) Measurand 2: LightMix® SarbecoV E-gene plus EAV control (Cat # 40-0776-10, TIB Molbiol) | qPCR Bio-Rad CFX96 | see Instrument |

Table H-2: Summary of Analytical Techniques for CCQM P199b Study Material 4

| Laboratory ID<br><br>(Institute code) | Analytical Technique | Instrument | Additional information |
|---|---|---|---|
| 5 | RT-dPCR | Bio-Rad QX100 | One-Step RT-ddPCR Advanced Kit for Probes (Bio-Rad) #1864021 |
| | ID-MS | Agilent 6510 LC-MS | |
| 6 | RT-dPCR | Bio-Rad QX200 | One-Step RT-ddPCR Advanced Kit for Probes (Bio-Rad) #1864022 |
| | ID-MS | Thermo TSQ Altis | |
| 7 | ID-MS | Shimadzu LCMS-8030+ | |
| 13 | Single molecule flow cytometry | Built in-house | |
| 15 | RT-dPCR | Bio-Rad QX200 | One-Step RT-ddPCR Advanced Kit for Probes (Bio-Rad) #1864021 |

Table H-3: PCR assay specifications CCQM-P199b

| Lab. ID (Institute code) | Study Material | Assay name/ abbreviation Duplex if performed | Oligonucleotide sequences (5' → 3')† | [Oligonucleotide final] (nM[1]) | Amplicon size (bp) | Supplier and purification |
|---|---|---|---|---|---|---|
| 1 | SM1, SM2, SM3 | CDC N2 (nominated result) | Forward: TTA CAA ACA TTG GCC GCA AA<br>Reverse: GCG CGA CAT TCC GAA GAA<br>Probe: FAM-ACA ATT TGC CCC CAG CGC TTC AG-QSY | Primers 900 nM<br>Probe 250 nM | 67 | Thermo Fisher Scientific |
| | SM1, SM2, SM3 | CDC N1 (supplementary result) | Forward: GAC CCC AAA ATC AGC GAA AT<br>Reverse: TCT GGT TAC TGC CAG TTG AAT CTG<br>Probe: FAM-ACC CCG CAT TAC GTT TGG TGG ACC-QSY | Primers 900 nM<br>Probe 250 nM | 72 | Thermo Fisher Scientific |
| 2 | SM1, SM2, SM3 | CDC N3 (nominated result) | Forward: GGG AGC CTT GAA TAC ACC AAA A<br>Reverse: TGT AGC ACG ATT GCA GCA TTG<br>Probe: FAM-AYC ACA TTG GCA CCC GCA ATC CTG-BHQ1 | Primers 500 nM<br>Probe 125 nM | 72 | Primers Invitrogen, Probes Eurofins Genomic |
| | SM1, SM2, SM3 | CDC N2 (supplementary result) | Forward: TTA CAA ACA TTG GCC GCA AA<br>Reverse: GCG CGA CAT TCC GAA GAA<br>Probe: FAM-ACA ATT TGC CCC CAG CGC TTC AG-BHQ1 | Primers 500 nM<br>Probe 125 nM | 67 | Primers Invitrogen, Probes Eurofins Genomic |
| 3 | SM1, SM2, SM3 | CDC N2 | Forward: TTA CAA ACA TTG GCC GCA AA<br>Reverse: GCG CGA CAT TCC GAA GAA<br>Probe: FAM-ACA ATT TGC /ZEN/ CCC CAG CGC TTC AG-BHQ1 | Primers 500 nM<br>Probe 125 nM | 67 | IDT |
| | SM1, SM2 | Sarbeco E | Forward: ACAGGTACGTTAATAGTTAATAGCGT<br>Reverse: ATATTGCAGCAGTACGCACACA<br>Probe: FAM-ACACTAGCCATCCTTACTGCGCTTCG- ZEN/ IowaBlack | Primers 400 nM<br>Probe 200 nM | 113 | IDT |
| 4 | SM1, SM2 | China CDC N / Sarbeco E (Duplex) | Forward: GGGGAACTTCTCCTGCTAGAAT<br>Reverse: CAGACATTTTGCTCTCAAGCTG<br>Probe: FAM-TTGCTGCTGCTTGACAGATT-BHQ1<br><br>Forward: ACAGGTACGTTAATAGTTAATAGCGT<br>Reverse: ATATTGCAGCAGTACGCACACA<br>Probe: HEX-ACACTAGCCATCCTTACTGCGCTTCG- BHQ1 | Primers 400 nM<br>Probe 200 nM | 99 / 113 | Eurofins genomics |
| | SM3 | China CDC N | Forward: GGGGAACTTCTCCTGCTAGAAT<br>Reverse: CAGACATTTTGCTCTCAAGCTG<br>Probe: FAM-TTGCTGCTGCTTGACAGATT-BHQ1 | Primers 400 nM<br>Probe 200 nM | 99 | Eurofins genomics |

| Lab. ID (Institute code) | Study Material | Assay name/ abbreviation Duplex if performed | Oligonucleotide sequences (5’ → 3’)† | [Oligonucleotide final] (nM[1]) | Amplicon size (bp) | Supplier and purification |
|---|---|---|---|---|---|---|
| 5 | SM1, SM2 | CDC N1 / Sarbeco E - NMIA modified† (duplex, SM2 only) | Forward: GAC CCC AAA ATC AGC GAA AT<br>Reverse: TCT GGT TAC TGC CAG TTG AAT CTG<br>Probe: FAM-ACC CCG CAT TAC GTT TGG TGG ACC-BHQ1<br><br>Forward: ACAGGTACGTAAATAGTTAATAGCGT<br>Reverse: ATATTGCAGCACTACGCACACA<br>Probe: HEX-ACACTAGCCATCCTTACTGCGCTTCG-BHQ1 | Primers: 900 nM<br>Probe 250 nM | 72 / 113 | Merck |
| | SM3, SM4 | CDC N2 / China CDC N (Duplex) | Forward: TTA CAA ACA TTG GCC GCA AA<br>Reverse: GCG CGA CAT TCC GAA GAA<br>Probe: FAM-ACA ATT TGC CCC CAG CGC TTC AG-BHQ1<br><br>Forward: GGGGAACTTCTCCTGCTAGAAT<br>Reverse: CAGACATTTTGCTCTCAAGCTG<br>Probe: HEX-TTGCTGCTGCTTGACAGATT-BHQ1 | Primers: 900 nM<br>Probe 250 nM | 67 / 99 | Merck |
| 6 | SM1, SM2, SM3, SM4 | China CDC N | Forward: GGGGAACTTCTCCTGCTAGAAT<br>Reverse: CAGACATTTTGCTCTCAAGCTG<br>Probe: FAM-TTGCTGCTGCTTGACAGATT-BHQ1 | Primers: 600 nM<br>Probe: 200 nM | 99 | Primers: Takara<br>Probe: BGI (Beijing, China) |
| | SM1 | Sarbeco E/China CDC Orf1ab (Duplex) | Forward: ACAGGTACGTTAATAGTTAATAGCGT<br>Reverse: ATATTGCAGCAGTACGCACACA<br>Probe: FAM-ACACTAGCCATCCTTACTGCGCTTCG-BBQ<br>Forward: CCCTGTGGGTTTTACACTTAA<br>Reverser:ACGATTGTGCATCAGCTGA<br>Probe:5'-FAM-CCGTCTGCGGTATGTGGAAAGGTTATGG-BHQ1-3' | Primers: 600 nM<br>Probe: 50 nM | 113 | Primers: Invitrogen<br>Probe: BGI (Beijing, China) |
| | SM2 | Sarbeco E (modified)† | Forward: ACAGGTACGTAAATAGTTAATAGCGT<br>Reverse: ATATTGCAGCACTACGCACACA<br>Probe: FAM-ACACTAGCCATCCTTACTGCGCTTCG-BBQ | Primers: 600 nM<br>Probe: 50 nM | 113 | Primers: Invitrogen<br>Probe: BGI (Beijing, China) |
| 7 | SM1, SM2, SM3 | China CDC N | Forward: GGGGAACTTCTCCTGCTAGAAT<br>Reverse: CAGACATTTTGCTCTCAAGCTG<br>Probe: FAM-TTGCTGCTGCTTGACAGATT-MGB-BHQ1 | Primers: 900 nM<br>Probe: 250 nM | 99 | Thermo Fisher Scientific |
| 8 | SM1 | In-house N ‘UME Set2’ | Forward:GTGATGCTGCTCTTGCTTTG<br>Reverse:GTGACAGTTTGGCCTTGTTG<br>Probe: FAM-TGACAGATT-ZEN-GAACCAGCTTGAGAGCA-IBFQ | Primers: 900 nM<br>Probe: 250 nM | 97 | Primers: Oligomer Biyoteknoloji (Turkey)<br>Probe: IDT, Integrated DNA Technologies |
| | SM3 | CDC N1 | Forward: GAC CCC AAA ATC AGC GAA AT<br>Reverse: TCT GGT TAC TGC CAG TTG AAT CTG<br>Probe: FAM-ACCCCGCAT-(ZEN)-TACGTTTGGTGGACC-IBFQ | Primers: 900 nM<br>Probe: 250 nM | 72 | Primers: Oligomer Biyoteknoloji (Turkey)<br>Probe: IDT, Integrated DNA Technologies |

| Lab. ID (Institute code) | Study Material | Assay name/ abbreviation Duplex if performed | Oligonucleotide sequences (5' → 3')† | [Oligonucleotide final] (nM[1]) | Amplicon size (bp) | Supplier and purification |
|---|---|---|---|---|---|---|
| | SM1 | Sarbeco E | Forward: ACAGGTACGTTAATAGTTAATAGCGT<br>Reverse: ATATTGCAGCAGTACGCACACA<br>Probe: FAM-ACACTAGCC-ZEN-ATCCTTACTGCGCTTCG-IBFQ | Primers: 900 nM<br>Probe: 250 nM | 113 | Primers: Oligomer Biyoteknoloji (Turkey)<br>Probe: IDT, Integrated DNA Technologies |
| 9 | SM1, SM2, SM3 | CDC N2 | Forward: TTA CAA ACA TTG GCC GCA AA<br>Reverse: GCG CGA CAT TCC GAA GAA<br>Probe: FAM-ACA ATT TGC CCC CAG CGC TTC AG-MGB-NFQ | Primers and probe: 250 nM | 67 | Primers: Eurofins<br>Probe: ABI |
| | SM1 | Sarbeco E | Forward: ACAGGTACGTTAATAGTTAATAGCGT<br>Reverse: ATATTGCAGCAGTACGCACACA<br>Probe: FAM-ACACTAGCCATCCTTACTGCGCTTCG- MGB-NFQ | Primers: 400 nM<br>Probe: 200 nM | 113 | Primers: Eurofins<br>Probe: ABI |
| | SM2 | Sarbeco E (modified)† | Forward: ACAGGTACGTAAATAGTTAATAGCGT<br>Reverse: ATATTGCAGCACTACGCACACA<br>Probe: FAM-ACACTAGCCATCCTTACTGCGCTTCG- MGB-NFQ | Primers: 400 nM<br>Probe: 200 nM | 113 | Primers: Eurofins<br>Probe: ABI |
| 10 | SM1 | ITA_INRiM_N / ITA_INRiM_E (Duplex) | Forward: CGATCAAAACAACGTCGGCC<br>Reverse: GGAACGCCTTGTCCTCGA<br>Probe: FAM-CACCGCTCTCACTCAACATGGC-BHQ1<br><br>Forward: CGTTTCGGAAGAGACAGGTACG<br>Reverse: AGCGCAGTAAGGATGGCTAGT<br>Probe: FAM-CTTGCTTTCGTGGTATTCTTGCT- BHQ1 | ITA_INRiM_N<br>Primers: 1350 nM<br>Probe: 375 nM<br><br>ITA_INRiM_E<br>Primers: 900 nM<br>Probe: 250 nM | 113 / 99 | Metabion International AG |
| | SM2 | China CDC N / ITA_INRiM_E (Duplex) | Forward: GGGGAACTTCTCCTGCTAGAAT<br>Reverse: CAGACATTTTGCTCTCAAGCTG<br>Probe: FAM-TTGCTGCTGCTTGACAGATT- BHQ1<br><br>Forward: CGTTTCGGAAGAGACAGGTACG<br>Reverse: AGCGCAGTAAGGATGGCTAGT<br>Probe: FAM-CTTGCTTTCGTGGTATTCTTGCT- BHQ1 | China CDC N<br>Primers: 1350 nM<br>Probe: 375 nM<br><br>ITA_INRiM_E<br>Primers: 900 nM<br>Probe: 250 nM | 99 / 99 | Metabion International AG |
| | SM3 | ITA_INRiM_N | Forward: CGATCAAAACAACGTCGGCC<br>Reverse: GGAACGCCTTGTCCTCGA<br>Probe: FAM-CACCGCTCTCACTCAACATGGC-BHQ1 | Primers: 1350 nM<br>Probe: 375 nM | 113 | Metabion International AG |
| 11 | SM1, SM2, SM3 | DNA-Technology N-gene | Exact sequences proprietary | Primers: 900 nM<br>Probe: 250 nM | 200 | DNA-Technology |
| | SM1, SM2, SM3 | China CDC N | Forward: GGGGAACTTCTCCTGCTAGAAT<br>Reverse: CAGACATTTTGCTCTCAAGCTG<br>Probe: FAM-TTGCTGCTGCTTGACAGATT- BHQ1 | Primers: 900 nM<br>Probe: 250 nM | 99 | DNA-Technology |

| Lab. ID (Institute code) | Study Material | Assay name/ abbreviation Duplex if performed | Oligonucleotide sequences (5’ → 3’)† | [Oligonucleotide final] (nM[1]) | Amplicon size (bp) | Supplier and purification |
|---|---|---|---|---|---|---|
| | SM1, SM2 | DNA-Technology E-gene | Exact sequences proprietary | Primers: 900 nM<br>Probe: 250 nM | 200 | DNA-Technology |
| | SM1, SM2 | Sarbeco E | Forward: ACAGGTACGTTAATAGTTAATAGCGT<br>Reverse: ATATTGCAGCAGTACGCACACA<br>Probe: FAM-ACACTAGCCATCCTTACTGCGCTTCG- BHQ1 | Primers: 900 nM<br>Probe: 250 nM | 113 | DNA-Technology |
| 12 | SM2 | China CDC N | Forward: GGGGAACTTCTCCTGCTAGAAT<br>Reverse: CAGACATTTTGCTCTCAAGCTG<br>Probe: FAM-TTGCTGCTGCTTGACAGATT- BHQ1 | 800 nM Forward primer, 900 nM Reverse primer, 300 nM Probe | 99 | LGC Biosearch technologies. Primers purification with RPC and probes purification with Dual HPLC. |
| | SM2, SM3 | CDC N2 | Forward: TTA CAA ACA TTG GCC GCA AA<br>Reverse: GCG CGA CAT TCC GAA GAA<br>Probe: FAM-ACA ATT TGC CCC CAG CGC TTC AG-BHQ1 | 800 nM Forward primer, 900 nM Reverse primer, 300 nM Probe | 67 | LGC Biosearch technologies. Primers purification with RPC and probes purification with Dual HPLC. |
| | SM2 | E_INM | Forward: CTTGCTTTCGTGGTATTCTTG<br>Reverse: ACGTTAACAATATTGCAGCA<br>Probe: FAM-CCTTACTGCGCTTCGATTGTGTGCGT-BHQ | 800 nM Forward primer, 900 nM Reverse primer, 300 nM Probe | 86 | LGC Biosearch technologies. Primers purification with RPC and probes purification with Dual HPLC. |
| | SM2 | Sarbeco E (modified)† | Forward: ACAGGTACGTAAATAGTTAATAGCGT<br>Reverse: ATATTGCAGCACTACGCACACA<br>Probe: FAM-ACACTAGCCATCCTTACTGCGCTTCG-BHQ | 800 nM Forward primer, 900 nM Reverse primer, 300 nM Probe | 113 | LGC Biosearch technologies. Primers purification with RPC and probes purification with Dual HPLC. |
| 13 | SM1, SM2, SM3 | In-house N assay | Forward: CAGCAGTAGGGGAACTTCTC<br>Reverse: GCTGGTTCAATCTGTCAAGC<br>Probe: FAM-TGATGCTGCTCTTGCTTTGCT - SFCQ2 | Primers: 1000 nM<br>Probe: 250 nM | 88 | IDT |
| | SM1 | In-house E assay | Forward: CGGAAGAGACAGGTACGTTAA Reverse: GCAGTAAGGATGGCTAGTGT<br>Probe: FAM-TCTTGCTTTCGTGGTATTCTTGCT-SFCQ2 | Primers: 1000 nM<br>Probe: 250 nM | 91 | IDT |

| Lab. ID (Institute code) | Study Material | Assay name/ abbreviation Duplex if performed | Oligonucleotide sequences (5’ → 3’)† | [Oligonucleotide final] (nM[1]) | Amplicon size (bp) | Supplier and purification |
|---|---|---|---|---|---|---|
| | SM2 | In-house E assay (modified) | Forward: TTCGGAAGAGACAGGTACGT<br>Reverse: GCAGTAAGGATGGCTAGTGT<br>Probe: FAM-TCTTGCTTTCGTGGTATTCTTGCT- SFCQ2 | Primers: 1000 nM<br>Probe: 250 nM | 91 | IDT |
| 14 | SM1, SM2, SM3 | CDC N1 | Forward: GAC CCC AAA ATC AGC GAA AT<br>Reverse: TCT GGT TAC TGC CAG TTG AAT CTG<br>Probe: FAM-ACC CCG CAT TAC GTT TGG TGG ACC-BHQ1 | Primers: 900 nM<br>Probe: 250 nM | 72 | Macrogen, Korea |
| | SM1, SM2 | Sarbeco E | Forward: ACAGGTACGTTAATAGTTAATAGCGT<br>Reverse: ATATTGCAGCAGTACGCACACA<br>Probe: HEX-ACACTAGCCATCCTTACTGCGCTTCG-BHQ1 | Primers: 900 nM<br>Probe: 250 nM | 113 | Macrogen, Korea |
| 15 | SM1, SM2, SM3, SM4 | CDC N2 | Forward: TTA CAA ACA TTG GCC GCA AA<br>Reverse: GCG CGA CAT TCC GAA GAA<br>Probe: FAM-ACA ATT TGC CCC CAG CGC TTC AG-BHQ1 | Primers: 500 nM<br>Probe: 125 nM | 67 | IDT both HPLC purified |
| | SM1, SM2 | Sarbeco E | Forward: ACAGGTACGTTAATAGTTAATAGCGT<br>Reverse: ATATTGCAGCAGTACGCACACA<br>Probe: FAM-ACACTAGCCATCCTTACTGCGCTTCG-BHQ1 | Primers: 500 nM<br>Probe: 125 nM | 113 | IDT both HPLC purified |
| 16 | SM1, SM2, SM3 | China CDC N | Forward: GGGGAACTTCTCCTGCTAGAAT<br>Reverse: CAGACATTTTGCTCTCAAGCTG<br>Probe: FAM-TTGCTGCTGCTTGACAGATT- BHQ1 | Primers: 900 nM<br>Probe: 250 nM | 99 | T4 Oligo |
| | SM1, SM2 | Sarbeco E | Forward: ACAGGTACGTTAATAGTTAATAGCGT<br>Reverse: ATATTGCAGCAGTACGCACACA<br>Probe: FAM-ACACTAGCCATCCTTACTGCGCTTCG-BHQ1 | Primers: 900 nM<br>Probe: 250 nM | 113 | T4 Oligo |
| 17 | SM1, SM2, SM3 | CDC N1 | Forward: GAC CCC AAA ATC AGC GAA AT<br>Reverse: TCT GGT TAC TGC CAG TTG AAT CTG<br>Probe: FAM-ACC CCG CAT TAC GTT TGG TGG ACC-ZEN/Iowa Black | Primers: 900 nM<br>Probe: 250 nM | 72 | Primers: CBER Core Facility<br>Probe: IDT |
| 18 | SM1, SM2, SM3 | CDC N2 | Forward: TTA CAA ACA TTG GCC GCA AA<br>Reverse: GCG CGA CAT TCC GAA GAA<br>Probe: FAM-ACA ATT TGC CCC CAG CGC TTC AG- ZEN/Iowa Black | Primers: 900 nM<br>Probe: 250 nM | 67 | IDT |

| Lab. ID (Institute code) | Study Material | Assay name/ abbreviation Duplex if performed | Oligonucleotide sequences (5' → 3')† | [Oligonucleotide final] (nM[1]) | Amplicon size (bp) | Supplier and purification |
|---|---|---|---|---|---|---|
| | SM1, SM2 | Sarbeco E | Forward: ACAGGTACGTTAATAGTTAATAGCGT<br>Reverse: ATATTGCAGCAGTACGCACACA<br>Probe: FAM-ACACTAGCCATCCTTACTGCGCTTCG-BBQ | Primers: 900 nM<br>Probe: 250 nM | 113 | IDT |
| 19 | SM1, SM2, SM3 | CDC N2 | Forward: TTA CAA ACA TTG GCC GCA AA<br>Reverse: GCG CGA CAT TCC GAA GAA<br>Probe: FAM-ACA ATT TGC CCC CAG CGC TTC AG-BHQnova1 | Primers: 900 nM<br>Probe: 250 nM | 67 | LGC Biosearch Technologies RP-HPLC |
| | SM1 | Sarbeco E | Forward: ACAGGTACGTTAATAGTTAATAGCGT<br>Reverse: ATATTGCAGCAGTACGCACACA<br>Probe: FAM-ACACTAGCCATCCTTACTGCGCTTCG- BHQnova1 | Primers: 900 nM<br>Probe: 250 nM | 113 | LGC Biosearch Technologies RP-HPLC |
| | SM2 | Sarbeco E (modified)† | Forward: ACAGGTACGTAAATAGTTAATAGCGT<br>Reverse: ATATTGCAGCACTACGCACACA<br>Probe: FAM-ACACTAGCCATCCTTACTGCGCTTCG-BHQnova1 | Primers: 900 nM<br>Probe: 250 nM | 113 | LGC Biosearch Technologies RP-HPLC |
| 20 | SM1, SM2, SM3 | China CDC N | Forward: GGGGAACTTCTCCTGCTAGAAT<br>Reverse: CAGACATTTTGCTCTCAAGCTG<br>Probe: FAM-TTGCTGCTGCTTGACAGATT- BHQ1 | Primers: 1200 nM<br>Probe: 300 nM | 99 | Syntol Ltd., Moscow, Russia. HPLC purified |
| | SM1, SM2 | Sarbeco E | Forward: ACAGGTACGTTAATAGTTAATAGCGT<br>Reverse: ATATTGCAGCAGTACGCACACA<br>Probe: FAM-ACACTAGCCATCCTTACTGCGCTTCG- BHQ1 | Primers: 1200 nM<br>Probe: 300 nM | 113 | Syntol Ltd., Moscow, Russia. HPLC purified |
| 21 | SM1, SM2, SM3 | HKU N | Forward: TAATCAGACAAGGAACTGATTA<br>Reverse: CGAAGGTGTGACTTCCATG<br>Probe: FAM-GCAAATTGTGCAATTTGCGG-TAMRA | Primers: 500 nM<br>Probe: 250 nM | 110 | Biolabmix, Russia |
| | SM1, SM2 | Sarbeco E | Forward: ACAGGTACGTTAATAGTTAATAGCGT<br>Reverse: ATATTGCAGCAGTACGCACACA<br>Probe: FAM-ACACTAGCCATCCTTACTGCGCTTCG- BBQ | Primers: 400 nM<br>Probe: 200 nM | 113 | TIB MolBio, Germany |

†Nucleotides modified to match SM2 E gene sequence are indicated in red.

[1] The molecular biology community express amount of substance concentration with units of molarity (M) (SI units: mmol/L).

Table H-4: Technique/method- specific parameters for CCQM P199b RT-dPCR

| Laboratory ID (Institute code) | Partition volume (nL) | Partition volume standard uncertainty (nL) (if considered†) | Partition volume basis* | Study Material (if given) | Target (if given – refer to Table H-3 for specific assay used) | Mean accepted partition (droplet) number | SD in partition number (metric not SD) | Software version |
|---|---|---|---|---|---|---|---|---|
| 1 | 0.85 | NA‡ | 3 | SM1 | N1<br>N2 | 15,380<br>14,102 | 1,435<br>1,547 | Bio-Rad QuantaSoft version 1.7.4.0917 |
| | | | | SM2 | N1<br>N2 | 16,075<br>14,562 | 1,559<br>2,016 | |
| | | | | SM3 | N1<br>N2 | 15,225<br>16,406 | 1,662<br>1,566 | |
| 2 | 0.735 | 0.022 | 4# | | N2<br>N3 | 16,324<br>15,880 | 1,233<br>1,564 | Bio-Rad QuantaSoft version 1.7.4.0917 |
| 3 | 0.85 | NA | 3 | | N2<br>E | 13,206<br>12,860 | 1,440<br>1,347 | Bio-Rad QuantaSoft version 1.7.4.0917 |
| 4 | 0.85 | NA | 3 | | China N<br>Sarbeco E | 13,737<br>13,053 | 1,196<br>1,447 | Bio-Rad QuantaSoft version 1.7.4.09171 |
| 5 | 0.76 | NA | 1 | | | 10,959 | 776 | Bio-Rad QuantaSoft version 1.7.4.0917 |
| 6 | 0.85 | 0.8% (0.0068 nL) | 3 | | | 13,634 | 993 | Bio-Rad QuantaSoft version 1.7.4.0917 |
| 7 | 0.7532 | 0.0048 | 1 | SM1 | | 17,052 | 562 | QuantStudio® AnalysisSuite version 3.1.6-PCR-build2 |
| | | | | SM2 | | 17,141 | 604 | |
| | | | | SM3 | | 17,063 | 693 | |
| 8 | 0.749 | NA | 2 (1-3) | SM1 | N<br>E | 14,113<br>11,077 | 2,206<br>3,598 | Bio-Rad QuantaSoft Analysis Pro version 1.0.596 |
| | | | | SM3 | N | 16,087 | 1,480 | |
| 9 | 0.7472 | 0.013083 | 1+4## | SM1 | | 10,350 | 2,514 | Bio-Rad QuantaSoft version 1.7.4.0917 |
| | | | | SM2 | | 10,897 | 2,149 | |
| | | | | SM3 | | 10,721 | 1,765 | |
| 10 | 0.76 | 0.05 | 1 | | | 11,508 | 1,587<br>(CV) 14% | Bio-Rad QuantaSoft Analysis Pro version 1.0.596 |

| Laboratory ID (Institute code) | Partition volume (nL) | Partition volume standard uncertainty (nL) (if considered†) | Partition volume basis* | Study Material (if given) | Target (if given – refer to Table H-3 for specific assay used) | Mean accepted partition (droplet) number | SD in partition number (metric not SD) | Software version |
|---|---|---|---|---|---|---|---|---|
| 11 | 0.85 | NA | 3 | | | 17,262 | 952 | Bio-Rad QuantaSoft version 1.7.4.0917 |
| 12 | 0.782 | 0.000025 | 1 (microscopy) | SM2 | E (INM)<br>E<br>N (China)<br>N2 | 12,832<br>11,310<br>14,907<br>15,528 | 2,507<br>1,209<br>1,705<br>951 | Bio-Rad QuantaSoft version 1.7.4.0917 |
| | | | | SM3 | N2 | 14,185 | 1,464 | |
| 13 | 0.868 | NA | 3 | | | 16,814.4 | 1,417.52 | Bio-Rad QuantaSoft version 1.7.4.0917 |
| 14 | 0.85 | 0.01235 | 3 | | | 18,690 | | Bio-Rad QuantaSoft version 1.7.4.0917 |
| 15 | 0.762 | 0.0300 | 2 (quadratic average from five values in literature (1, 2, 4-6), ponderated by uncertainty<br><br>as in ISO Guide 35:2017 eq A.2). | | | 16,480 | 1,830 | QuantaSoft Analysis Pro version 1.0.596 (data analysis);<br>Bio-Rad QuantaSoft version 1.7.4.0917 (data collection) |
| 16 | 0.85 | NA | 3 | | | 12,847 | 1,799 | Bio-Rad QuantaSoft version 1.7.4.0917 |
| 17 | 0.85 | NA | 3 | SM1 | | 14,057 | 1,613 | Bio-Rad QuantaSoft Analysis Pro version 1.0.596.0525 |
| | | | | SM2 | | 13,914 | 1,855 | |
| | | | | SM3 | | 12,721 | 1,657 | |

| Laboratory ID (Institute code) | Partition volume (nL) | Partition volume standard uncertainty (nL) (if considered[†]) | Partition volume basis* | Study Material (if given) | Target (if given – refer to Table H-3 for specific assay used) | Mean accepted partition (droplet) number | SD in partition number (metric not SD) | Software version |
|---|---|---|---|---|---|---|---|---|
| 18 | 0.85 | NA | 3 | SM1 | | 16,770.75 | 1,220.478677 | QuantaSoft version 1.7.4.0917 and QuantaSoft Analysis Pro 1.0.596 |
| | | | | SM2 | | 17,328.1875 | (SD) 1,311.754104 | |
| | | | | SM3 | | 16,777.4375 | (SD) 1,510.913277 | |
| 19 | 0.776 | 0.0403 | 2 (1, 2, 7) + 4[##] | SM1 | N2<br>E Sarbeco<br>All assays | 14,832<br>15,803<br>15,317 | ± 1,311<br>± 1,430<br>± 1,439 | Bio-Rad QuantaSoft version 1.7.4.0917 |
| | | | | SM2 | N2<br>E Sarbeco (mod)<br>All assays | 14,247<br>14,945<br>14,596 | ± 1,155<br>± 1,087<br>± 1,161 | |
| | | | | SM3 | N2 | 14,879 | ± 1,303 | |
| 20 | 0.793 | NA | 1 | | | 9,573 | ± 2,434 | Bio-Rad QuantaSoft version 1.7.4.0917 |

† Uncertainty values given as reported by participants

‡ NA – not applicable

*Basis for partition volume value: 1 = in-house measurement; 2 = Literature; 3 = Bio-Rad value; 4 = other. [#]personal communication [##]reagent comparison.

Table H-5: <u>Reverse transcription and thermal cycling parameters for </u>CCQM P199b RT-dPCR/RT-qPCR

| Lab. ID | Assay summary | RT temp (°C) | RT time (min) | PCR intial step temp | PCR intial step time (min) | PCR cycling temp 1 (°C) | PCR cycling time 1 (s) | PCR cycling temp 2 (°C) | PCR cycling time 2 (s) | Cycle number | PCR final incubation (Hold) | Ramp rate (ddPCR only) |
|---|---|---|---|---|---|---|---|---|---|---|---|---|
| 1 | CDC N1, N2 | 45 | 60 | 95 °C | 10 | 95 | 30 | 60 | 60 | 40 | 4 °C ≥ 40 min | |
| 2 | CDC N2, N3 | 50 | 60 | 95 °C | 10 | 95 | 30 | 58 | 60 | 50 | 4 °C | |
| 3 | CDC N2<br>Sarbeco E | 50 | 60 | 95 | 10 | 95 | 30 | 55 | 60 | 45 | 4 °C | 2 °C/s |
| 4 | China CDC N<br>Sarbeco E | 50 | 15 | 95 °C | 5 | 95 | 30 | 55 | 30 | 45 | 4 °C | 2 °C/s |
| 5 | CDC N1/Sarbeco E<br>China CDC N/CDC N2 | 45 | 60 | 95 °C | 10 | 95 | 30 | 57 | 60 | 50 | | |
| 6 | China CDC N<br>Sarbeco E | 45 | 10 | 95 °C | 5 | 95 | 15 | 58 | 30 | 40 | | |
| 7 | China CDC N | 55 | 10 | 96 ºC | 10 | 98 | 30 | 56 | 120 | 40 | 10 °C | |
| 8 | In house N / CDC N1<br>Sarbeco E | 42 | 15 | 95 °C | 10 | 94 | 30 | 61 | 60 | 40 | 4 °C | |
| 9 | CDC N2<br>Sarbeco E | 50 | 60 | 95 °C | 10 | 95 | 30 | 55 | 60 | 60 | 4 °C | |
| 10 | In house N/China CDC N<br>In house E | 45 | 60 | 95°C | 5 | 95 | 30 | 56 | 60 | 40 | 4 °C | 2 °C/s |
| 11 | China CDC N | 40 | 30 | 95 °C | 10 | 94 | 30 | 58 | 60 | 40 | 10 °C | |
| | Commercial N / E | | | | 10 | | | 62 | | | | |
| 12 | China CDC N/ CDC N2 | 50 | 60 | 95 °C | 10 | 95 | 15 | 59 (N) | 30 | 45 | 4 °C | 0.5 °C/s |
| | In house E / Sarbeco E | | | | | | | 56 (E) | | | | |
| 13 | In house N<br>In house E | 42 | 60 | 95 °C | 10 | 95 | 30 | 59 | 150 | 70 | 4 °C | 2 °C/s |
| 14 | CDC N1<br>Sarbeco E | 45 | 60 | 95 °C | 10 | 94 | 30 | 63 | 60 | 40 | 4 °C | |
| 15 | CDC N2<br>Sarbeco E | 50 | 60 | 95 °C | 2 | 95 | 30 | 55 | 30 | 35 | 16 °C | |
| 16 | China CDC N | 50 | 60 | 95 °C | 10 | 95 | 30 | 58 (N) | 60 | 45 | | |
| | Sarbeco E | 45 | 10 | 95 °C | 5 | 95 | 15 | 57 (E) | 30 | 45 | | |
| 17 | CDC N1 | 50 | 60 | 95 °C | 10 | 95 | 30 | 55 | 60 | 40 | | |
| 18 | CDC N2 | 43 | 60 | 98 °C | 10 | 95 | 30 | 55 (N) | 60 | 40 | 12 °C | |

| Lab. ID | Assay summary | RT temp (°C) | RT time (min) | PCR intial step temp | PCR intial step time (min) | PCR cycling temp 1 (°C) | PCR cycling time 1 (s) | PCR cycling temp 2 (°C) | PCR cycling time 2 (s) | Cycle number | PCR final incubation (Hold) | Ramp rate (ddPCR only) |
|---|---|---|---|---|---|---|---|---|---|---|---|---|
| | Sarbeco E | | | | | | | 58 (E) | | | | |
| 19 | CDC N2<br>Sarbeco E | 47.5 | 60 | 95 °C | 10 | 95 | 30 | 55 | 60 | 40 | 4 °C ≥ 1 h | 2°C/s |
| 20 | China CDC N<br>Sarbeco E | 50 | 60 | 95 °C | 10 | 95 | 30 | 57 | 60 | 60 | 4 °C | 2°C/s |
| 21 | HKU N | 50 | 5 | 95 °C | 20 s | 95 | 5 | 60 | 30 | 40 | | |
| | Sarbeco E | 55 | 10 | 95 °C | 3 | 95 | 15 | 58 | 30 | 45 | N/A | N/A |

Table H-6: Additional Comments for CCQM-P199b

| Laboratory ID | Additional Comments |
|---|---|
| 5 | Two assays were applied for the analysis of SM3 and SM4: CDC N2 and China CDC N, in duplex. The submitted result is an average of the values obtained using each of these assays.<br>Reported RT-dPCR values for all Study Materials were corrected for reverse transcription (RT) efficiency calculation using (i) synthetic short oligonucleotides analysed by ID-MS and RT-dPCR (Study Materials 1 and 2 results) (ii) Study Material 4 (for Study Material 3 and 4 results). |
| 11 | Two assays were employed for analysis of Measurand 1 (SM1, SM2, SM3) and Measurand 2 (SM1, SM2). Submitted results were an average of the values obtained for each of the assays per measurand. |
| 12 | For SM2, two assays were employed for analysis of each of Measurand 1 and 2. Submitted results were an average of the values obtained for each of the assays per measurand. |
| 19 | Prior to analysis by RT-dPCR, all nucleic acid templates (SM1, SM2, SM3) were heat denatured for 5 minutes at 65°C, then quenched on ice prior to loading for at least 1 min. |

Table H-7: preparation of SM4 for laboratories performing RT-dPCR analysis

| Laboratory ID | Dilution | Temperature | Diluent | Number of tubes analysed | Additional information |
|---|---|---|---|---|---|
| 5 | Gravimetric | Ambient (22°C) | Proprietary based on 1 mM citrate pH 6.5 with yeast total RNA at 5 ng/microliter | 4 | All tubes of SM4 supplied were pooled, then a 100 microlitre subsample was gravimetrically diluted for dPCR analysis. |
| 6 | Gravimetric | Ambient | RNA storage solution from Thermo Fisher,AM7001 | 1 | |
| 15 | Gravimetric | Ambient (23 +/- 1 °C) | Molecular biology grade water | 1 | gravimetric dilution in three steps: ~1:400 ; ~1:400 and then ~1:60 |

Table H-8: Partition volume (Vp) values measured in the study

| Lab. ID | Vp (nL) |
|---|---|
| 5 | 0.76 |
| 10 | 0.76 |
| 12 | 0.782 |
| 20 | 0.793 |
| Mean | 0.774 |
| SD | 0.0165 |

### REFERENCES


1. Kosir AB, Divieto C, Pavsic J, Pavarelli S, Dobnik D, Dreo T, Bellotti R, Sassi MP, Zel J. Droplet volume variability as a critical factor for accuracy of absolute quantification using droplet digital PCR. Anal Bioanal Chem. 2017;409(28):6689-97.
2. Dagata JA, Farkas N, Kramar JA. Method for Measuring the Volume of Nominally 100 µm Diameter Spherical Water-in-Oil Emulsion Droplets. NIST Special Publication 260-184. 2016.
3. Pinheiro LB, Coleman VA, Hindson CM, Herrmann J, Hindson BJ, Bhat S, Emslie KR. Evaluation of a droplet digital polymerase chain reaction format for DNA copy number quantification. Anal Chem. 2012;84(2):1003-11.
4. Corbisier P, Pinheiro L, Mazoua S, Kortekaas AM, Chung PY, Gerganova T, Roebben G, Emons H, Emslie K. DNA copy number concentration measured by digital and droplet digital quantitative PCR using certified reference materials. Anal Bioanal Chem. 2015;407(7):1831-40.
5. Mehle N, Gregur L, Bogožalec Košir A, Dobnik D. One-Step Reverse-Transcription Digital PCR for Reliable Quantification of Different Pepino Mosaic Virus Genotypes. Plants (Basel, Switzerland). 2020;9(3).
6. Emslie KR, JL HM, Griffiths K, Forbes-Smith M, Pinheiro LB, Burke DG. Droplet Volume Variability and Impact on Digital PCR Copy Number Concentration Measurements. Anal Chem. 2019;91(6):4124-31.
7. Pinheiro LB, O'Brien H, Druce J, Do H, Kay P, Daniels M, You J, Burke D, Griffiths K, Emslie KR. Interlaboratory Reproducibility of Droplet Digital Polymerase Chain Reaction Using a New DNA Reference Material Format. Anal Chem. 2017;89(21):11243-51.
