## Supplementary material for "CCQM-P199b: Interlaboratory comparability study of SARS-CoV-2 RNA copy number quantification": CCQM P199b supplementary file Appendix I

### APPENDIX I: Summary of Participants’ Uncertainty Estimation Approaches

Table I1 summarises factors included in the uncertainties reported by laboratories for Study Materials 1-3 (RT-dPCR and RT-qPCR (laboratory 21)) and those using RT-dPCR for Study Material 4. Table I2 summarises factors included in the uncertainties reported by laboratories using orthogonal methods for the analysis of Study Material 4.

Table I-1A: Summary of measurement uncertainty sources considered for measurement of Study Materials 1-3 and Study Material 4 (RT-dPCR)

| **MU Type** | **A** | **A/B*** | **A** | **A/B** | **A/B** | **A/B** | **B** | **B** | **B** | **B** | **B** | **B** | **B** | **B** | **B** |
|---|---|---|---|---|---|---|---|---|---|---|---|---|---|---|---|
| **Factor** / Laboratory ID | **RT-d/qPCR Method repeatbility** | **RT-d/qPCR intermediate precision** | **Between vial** | **Assay** | **Threshold setting** | **Sample dilution (volumetric)** | **Sample dilution (gravimetric)** | **Reaction preparation (volumetric)** | **Reaction preparation (gravimetric)** | **Homogeneity** | **Partition volume** | **RT efficiency** | **Poisson error** | **Material integrity** | **Nanodrop calibration** |
| 1 | ✓ | | ✓ | | | | | | | | | | | | |
| 2 | ✓ | ✓ | | | | | | | | ✓ | ✓ | | | | |
| 3 | ✓ | ✓ | | | | ✓ (B) | | | | ✓ | ✓ | | | | |
| 4 | ✓ | ✓ (pooled SD) | | | | | | | | | | | | | |
| 5 | ✓ | ✓ | | ✓ (A, SM3) | | | ✓ (SM1, 3) | | ✓ | ✓ | ✓ | ✓ | | | |
| 5 SM4 | ✓ | ✓ | | ✓ | | | ✓ | | ✓ | ✓ | ✓ | ✓ | | | |
| 6 | ✓ | ✓ | ✓ | | | | | | | | ✓ | | | | |
| 6 SM4 | ✓ | ✓ | | | | | ✓ | ✓ | | | ✓ | | | | |
| 7 | ✓ | ✓ | | | | | | | ✓ | | ✓ | | | | |
| 8 | ✓ | ✓ | | | | | | | | | | | ✓ | | |
| 9 [1] | ✓ | ✓ (pooled SD) | | | | | | | | | ✓ | | | | |
| 10 | ✓ | ✓ | | | | ✓ (B) | | | | | ✓ | | | | |

**KEY**: *RT-dPCR intermediate precision: If not stated in brackets, Type A approach.

[1] Uncertainties combined using NIST uncertainty machine (https://uncertainty.nist.gov/)

Table I-1B: Summary of measurement uncertainty sources *continued*

| **MU type** | **A** | **A/B*** | **A** | **A/B** | **A/B** | **A/B** | **B** | **B** | **B** | **B** | **B** | **B** | **B** | **B** | **B** |
|---|---|---|---|---|---|---|---|---|---|---|---|---|---|---|---|
| **Factor**<br>Laboratory ID | **RT-d/qPCR repeatbility** | **RT-d/qPCR intermediate precision** | **Between vial** | **Assay** | **Threshold setting** | **Sample dilution (volumetric)** | **Sample dilution (gravimetric)** | **Reaction preparation (volumetric)** | **Reaction preparation (gravimetric)** | **Homogeneity** | **Partition volume** | **RT efficiency** | **Poisson error** | **Material integrity** | **Nanodrop calibration** |
| 11 | ✓ | ✓ | | ✓ (A) | | ✓ (B) | | | | ✓ | | | | | ✓[2] |
| 12 | ✓ | ✓ | | ✓ (A, SM2) | | ✓ (B) | | | | | ✓ | | ✓ | | |
| 13 | ✓ | ✓ | | | ✓ (B) | | | | | ✓ | ✓ | | | | |
| 14 | ✓ | ✓ | | | | | | | | | ✓ | | | | |
| 15 | ✓ | | | | ✓ (A) | | | | ✓ | | ✓ | | | | |
| 15 (SM4) | ✓ | ✓ (B) | | | ✓ (A) | | ✓ | | ✓ | | ✓ | | | | |
| 16 | ✓ | ✓ | ✓ | | | ✓ (SM1, 3) (B) | | | | | | | | | |
| 17 | ✓ | ✓ (pooled SD) | | | | | | | | | | | | | |
| 18 | ✓ | | ✓ (pooled SD) | | | | | | | | | | | | |
| 19 | ✓ | ✓ | ✓ | ✓ (B) | | | | | | | ✓ | | | | |
| 20 | ✓ | ✓ | | | | ✓ (B) | | | | ✓ | ✓ | | | | |
| 21 | | ✓ | | | | | | | | | N/A | | | | |

**KEY**: *If not stated in brackets, Type A approach.

[2] Laboratory 11: Initial evaluation of RNA concentration measured using NanoDrop.

Table I-2 Summary of Participants' Uncertainty Estimation for SM4 (ID-MS and single molecule flow cytometry)

| **Laboratory ID** | **Factors included in uncertainty calculations** |
|---|---|
| 5 | IDMS -Between-subsamples, IDMS -Between NMPs, QNMR-molar abs coefficient –GMP, QNMR-molar abs coefficient –CMP, QNMR-molar abs coefficient –AMP, QNMR-molar abs coefficient –UMP, Purity |
| 6 | IDMS -Between-subsamples, IDMS -Between NMPs, preparation of standard and sample, uncertainty of NMP calibrators |
| 7 | Acid hydrolysis-IDMS, enzymatic hydrolysis-IDMS, between methods, preparation of standard and sample, density |
| 13 | Single molecule flow cytometry – method repeatability and day-to-day reproducibility |
