## Supplementary material for "CCQM-P199b: Interlaboratory comparability study of SARS-CoV-2 RNA copy number quantification": CCQM P199b supplementary file Appendix J

### APPENDIX J: Statistical Analysis

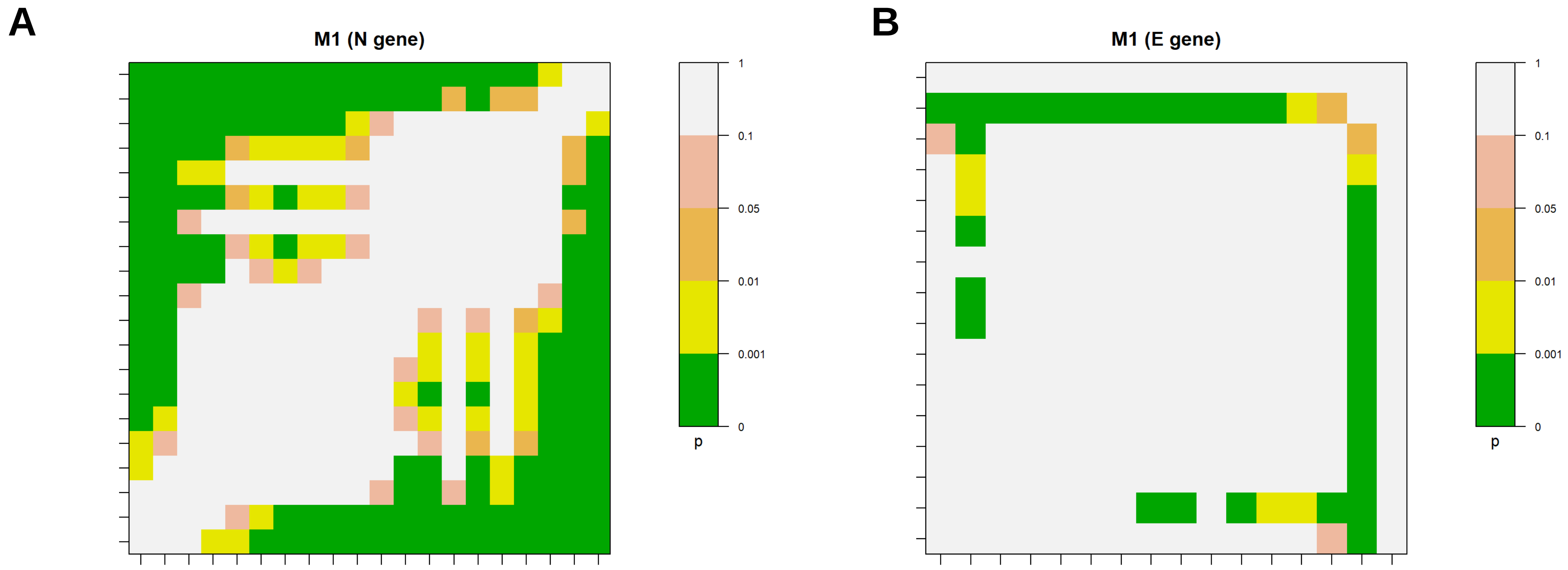


**Figure J-1: Study Material 1: Pairwise comparison consistency plots**.
Consistency plots for Measurand 1 (N gene) and 2 (E gene) indicate the result of pairwise comparison of the difference between two laboratories' results ordered from low to high (x-axis, left to right; y-axis, top to bottom; each dash represents a laboratory as per main manuscript **Figure 13A-B**) compared to the uncertainty in the difference (based on the laboratories' reported uncertainties). Significance levels of inconsistent results are indicated by colours, with the scale of p-values shown in the key.

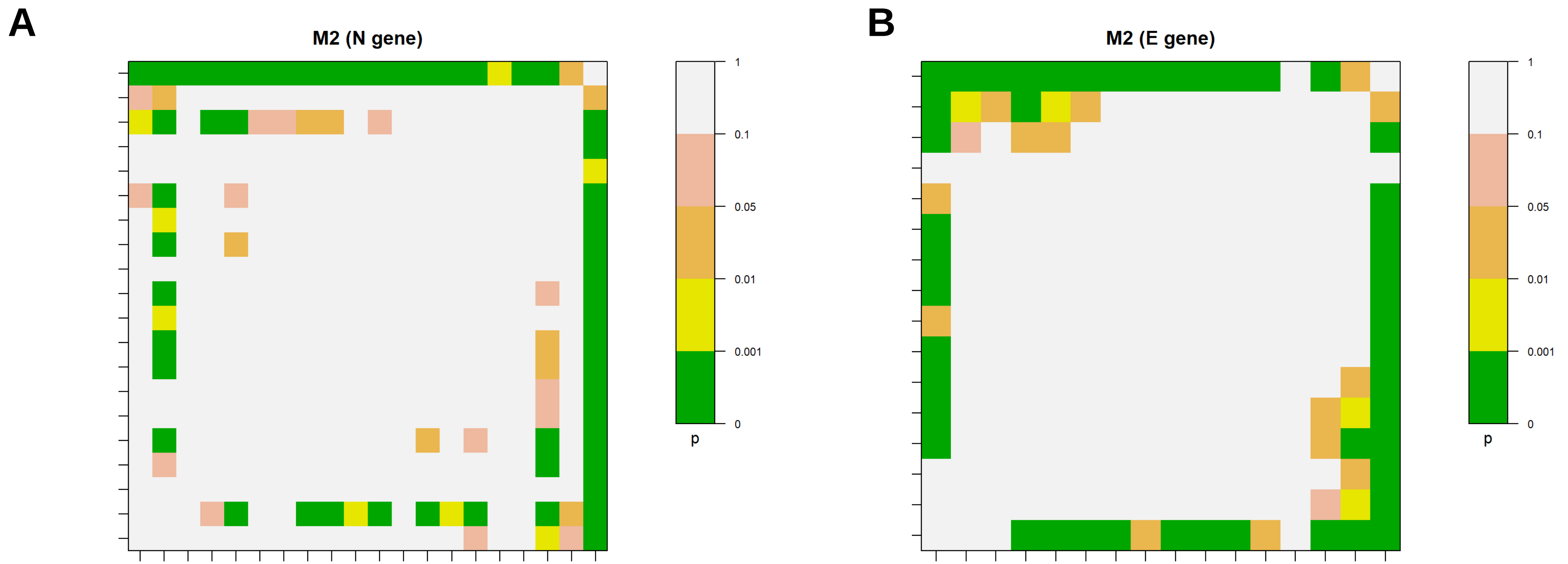


**Figure J-2: Study Material 2: Pairwise comparison consistency plots.**
Consistency plots for Measurand 1 (N gene) and 2 (E gene) indicate the result of pairwise comparison of the difference between two laboratories' results ordered from low to high (x-axis, left to right; y-axis, top to bottom; each dash represents a laboratory as per main manuscript **Figure 14A-B**) compared to the uncertainty in the difference (based on the laboratories' reported uncertainties). Significance levels of inconsistent results are indicated by colours, with the scale of [illegible]

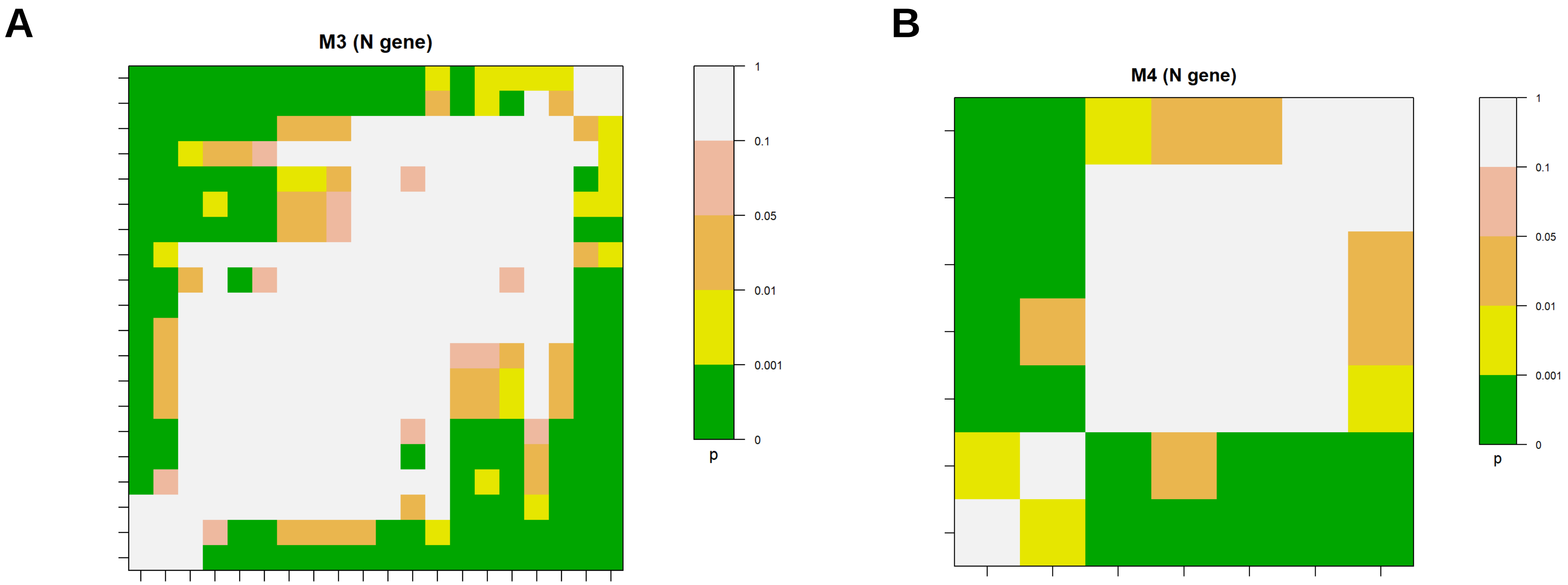


**Figure J-3: Study Materials 3-4: Pairwise comparison consistency plots.**
Consistency plots for Measurand 1 (N gene) indicate the result of pairwise comparison of the difference between two laboratories' results ordered from low to high (x-axis, left to right; y-axis, top to bottom; each dash represents a laboratory as per main manuscript **Figure 15A-B**) compared to the uncertainty in the difference (based on the laboratories' reported uncertainties). Significance levels of inconsistent results are indicated by colours, with the scale of p-values shown in the key.
