## Supplementary material for "CCQM-P199b: Interlaboratory comparability study of SARS-CoV-2 RNA copy number quantification": CCQM P199b supplementary file Appendix K

### APPENDIX K: Supplementary and follow-up data

#### Additional results for Study Material 4 (Laboratory 6)

Additional units of P199b Study Material 4 were analysed to evaluate the impact of different dilution approaches, and to assess unit homogeneity. Different concentrations of yeast RNA were added to RNA storage solution buffer (SS) to dilute the samples. Three units were analysed using the China CDC N, and US CDC N2 and N3 assays, with each unit being analysed on a different day. RNA copy number concentration was found to increase with the concentration of yeast RNA added.

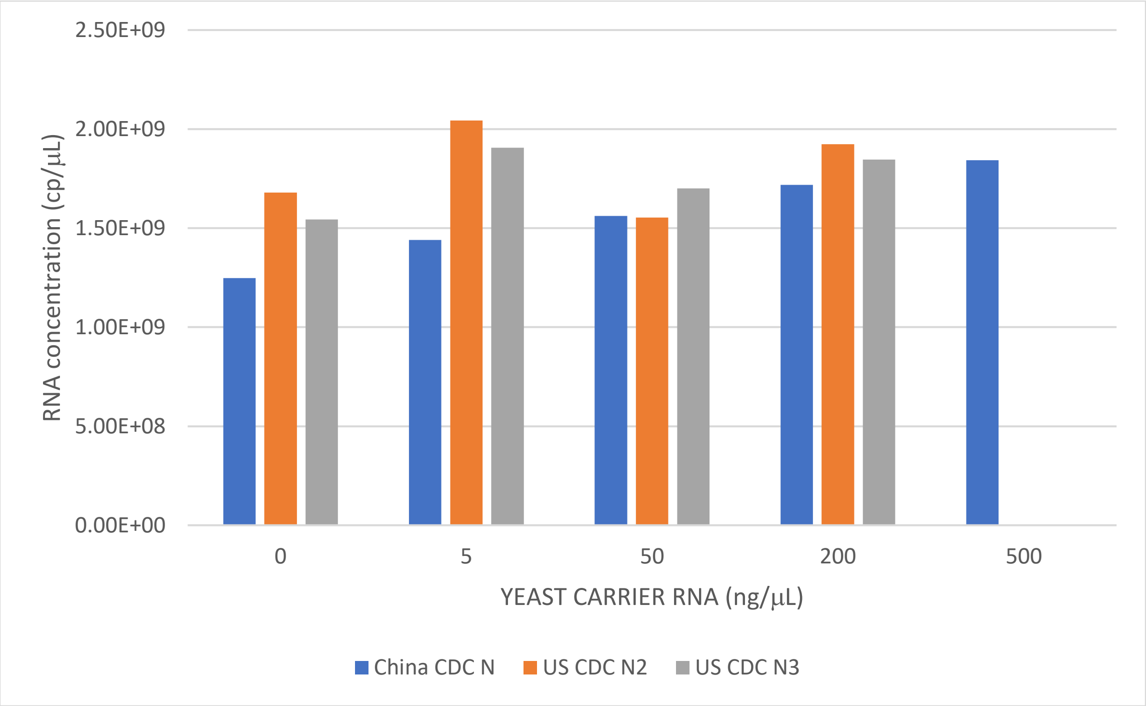

Figure K-1: Follow-up results for Study Material 4

Table K-1. Study Material 4 Follow-up Results (China CDC N assay)

| Concentration of yeast RNA added to SS (ng/µL) | Results of the three tubes (/µL) | | |
|---|---|---|---|
| | Tube 1-014 | Tube 2-037 | Tube 3-040 |
| 0 | 1.35E+09 | 1.28E+09 | 1.11E+09 |
| | 1.33E+09 | 1.30E+09 | 1.12E+09 |
| | 1.38E+09 | 1.25E+09 | 1.11E+09 |
| 5 | / | / | / |
| | 1.47E+09 | / | / |

| | | | |
|---|---|---|---|
| | 1.41E+09 | / | / |
| 50 | 1.45E+09 | 1.77E+09 | 1.49E+09 |
| | 1.38E+09 | 1.64E+09 | 1.52E+09 |
| | 1.52E+09 | 1.70E+09 | 1.59E+09 |
| 200 | 1.80E+09 | 1.72E+09 | 1.67E+09 |
| | 1.72E+09 | 1.74E+09 | 1.63E+09 |
| | 1.73E+09 | 1.82E+09 | 1.64E+09 |
| 500 | / | 1.86E+09 | 1.79E+09 |
| | / | 1.96E+09 | 1.79E+09 |
| | / | 1.92E+09 | 1.74E+09 |

Table K-2. Results of US CDC N2 assay

| Concentration of yeast RNA added to SS (ng/µL) | Results of the three tubes (/µL) | | |
|---|---|---|---|
| | Tube 1-014 | Tube 2-037 | Tube 3-040 |
| 0 | 1.83E+09 | 1.63E+09 | 1.58E+09 |
| | 1.84E+09 | 1.64E+09 | 1.60E+09 |
| | 1.83E+09 | 1.61E+09 | 1.55E+09 |
| 5 | 2.07E+09 | / | / |
| | 2.04E+09 | / | / |
| | 2.02E+09 | / | / |
| 50 | 1.53E+09 | / | / |
| | 1.57E+09 | / | / |
| | 1.56E+09 | / | / |
| 200 | 1.85E+09 | 1.99E+09 | 1.86E+09 |
| | 1.81E+09 | 1.96E+09 | 2.04E+09 |
| | 1.81E+09 | 1.98E+09 | 2.01E+09 |

Table K-3. Results of US CDC N3 assay

| Concentration of yeast RNA added to SS (ng/µL) | Results of the three tubes (/µL) | | |
|---|---|---|---|
| | Tube 1-014 | Tube 2-037 | Tube 3-040 |
| 0 | 1.80E+09 | 1.40E+09 | 1.42E+09 |
| | 1.81E+09 | 1.34E+09 | 1.44E+09 |
| | 1.83E+09 | 1.42E+09 | 1.44E+09 |
| 5 | 1.90E+09 | / | / |
| | 1.89E+09 | / | / |
| | 1.93E+09 | / | / |
| 50 | 1.72E+09 | / | / |
| | 1.69E+09 | / | / |
| | 1.69E+09 | / | / |
| 200 | 1.88E+09 | 1.84E+09 | 1.86E+09 |

| | | | |
|---|---|---|---|
| | 1.92E+09 | 1.84E+09 | 1.80E+09 |
| | 1.87E+09 | 1.79E+09 | 1.82E+09 |

Table K-4. Results of SM4

| Study Material 4 | Measurand 1 (*N* gene) (/μL) (dPCR) |
|---|---|
| Value (*x*) | $1.81 \times 10^9$ |
| Standard uncertainty (*u*) | $0.09 \times 10^9$ |
| Coverage factor (*k*) | 2 |
| Expanded uncertainty (*U*) | $0.18 \times 10^9$ |
| Relative expanded uncertainty (Rel *U*) | 9.7 % |

**Supplementary one-step RT-dPCR results (Laboratory 8)**

All four Study Materials were analysed by one-step RT-dPCR using two assays targeting Measurand 1 (*N* gene). In addition, Study Materials 1 and 2 were analysed using assays targeting Measurand 2 (*E* gene). Oligonucleotide sequences are described in Table H-3. For Study Material 2, modified primers were applied to account for polymorphisms in the nucleotide sequence of this material (Forward: ACAGGTACGTAAATAGTTAATAGCGT, Reverse: ATATTGCAGCACTACGCACACA). Two units of each material were analysed across two days in duplicate per day. Prior to analysis, Study Material 4 was diluted gravimetrically in water to a total dilution factor of 2.02 x $10^5$. RT-dPCR was performed using a One-Step RT-ddPCR Advanced Kit for Probes (Bio-Rad, #1864022) on the QX200 dPCR platform. 5.5 µL of RNA template was added per reaction. Results are presented in Table K-5.

Table K-5. Supplemental one-step RT-dPCR Results

| | |
|---|---|
| **STUDY MATERIAL 1** | |
| **Measurand 1 (*N* gene with in house Assay –CDC-N1)** (/µL) | |
| Value (*x*) | 1216 |
| Standard uncertainty (*u*) | 81 |
| Coverage factor (*k*) | 2 |
| Expanded uncertainty (*U*) | 162 |
| Relative expanded uncertainty (Rel *U*) | 14 |
| **Measurand 1 (*N* gene measured with Set2 Assay)** (/µL) | |
| Value (*x*) | 1608 |
| Standard uncertainty (*u*) | 127 |
| Coverage factor (*k*) | 2 |
| Expanded uncertainty (*U*) | 254 |
| Relative expanded uncertainty (Rel *U*) | 16 |
| **Measurand 2 (*E* gene measured with E Sarbeco Assay)** (/µL) | |
| Value (*x*) | 842 |
| Standard uncertainty (*u*) | 58 |
| Coverage factor (*k*) | 2 |
| Expanded uncertainty (*U*) | 115 |
| Relative expanded uncertainty (Rel *U*) | 14 |
| **STUDY MATERIAL 2** | |
| **Measurand 1 (*N* gene with in house Assay –CDC-N1)** (/µL) | |
| Value (*x*) | 14 |
| Standard uncertainty (*u*) | 2 |

| | |
|---|---|
| Coverage factor (*k*) | 2 |
| Expanded uncertainty (*U*) | 3 |
| Relative expanded uncertainty (Rel *U*) | 23 |
| **Measurand 1 (*N* gene measured with Set2 Assay)** (/μL) | |
| Value (*x*) | 14 |
| Standard uncertainty (*u*) | 1 |
| Coverage factor (*k*) | 2 |
| Expanded uncertainty (*U*) | 2 |
| Relative expanded uncertainty (Rel *U*) | 16 |
| **Measurand 2 (*E* gene measured with E Sarbeco Assay)** (/μL) | |
| Value (*x*) | 14 |
| Standard uncertainty (*u*) | 2 |
| Coverage factor (*k*) | 2 |
| Expanded uncertainty (*U*) | 4 |
| Relative expanded uncertainty (Rel *U*) | 30 |
| **STUDY MATERIAL 3** | |
| **Measurand 1 (*N* gene with in house Assay –CDC-N1)** (/μL) | |
| Value (*x*) | 1950 |
| Standard uncertainty (*u*) | 39 |
| Coverage factor (*k*) | 2 |
| Expanded uncertainty (*U*) | 76 |
| Relative expanded uncertainty (Rel *U*) | 4 |
| **Measurand 1 (*N* gene measured with Set2 Assay)** (/μL) | |
| Value (*x*) | 2202 |
| Standard uncertainty (*u*) | 50 |
| Coverage factor (*k*) | 2 |
| Expanded uncertainty (*U*) | 100 |
| Relative expanded uncertainty (Rel *U*) | 5 |
| **STUDY MATERIAL 4** | **Measurand 1 (*N* gene with in house Assay – Set2)** (/μL) |
| Value (*x*) | 1.18E+09 |

| | |
|---|---|
| Standard uncertainty (*u*) | 1.58E+08 |
| Coverage factor (*k*) | 2 |
| Expanded uncertainty (*U*) | 3.16E+08 |
| Relative expanded uncertainty (Rel *U*) | 27 |
| **STUDY MATERIAL 4** | **Measurand 1 (*N* gene measured with CDC-N1 Assay)** (/μL) |
| Value (*x*) | 1.00E+09 |
| Standard uncertainty (*u*) | 1.09E+08 |
| Coverage factor (*k*) | 2 |
| Expanded uncertainty (*U*) | 2.17E+08 |
| Relative expanded uncertainty (Rel *U*) | 22 |

**Supplementary two-step RT-dPCR results (Laboratory 20)**

Study Materials 1, 2 and 3 were analysed by two-step RT-dPCR using assays targeting Measurand 1 (*N* gene) and Measurand 2 (*E* gene). Assay details are described in Table H-3. Each material was analysed in quadruplicate using 10 µL RNA per reverse transcription (RT) reaction. cDNA generation was performed using gene-specific priming (800 nM[1] primer per reaction) and RevertAid H Minus First Strand cDNA Synthesis Kit (ThermoScientific, #K1632). RNA templates were pre-heating with gene-specific primers at 99°C for 1 min with subsequent quenching on ice. Analysis of cDNA by dPCR was performed using ddPCR Supermix for Probes (Bio-Rad, #1863026) and analysed on the Bio-Rad QX200 platform. Results are presented in Table K-6.

Table K-6. Two-step RT-dPCR Results

| Study Material | Measurand | Two-step Result (/µL) |
|---|---|---|
| 1 | 1 | 2250 |
| | 2 | 1260 |
| 2 | 1 | 25.9 |
| | 2 | 24.6 |
| 3 | 1 | 2684 |

[1] The molecular biology community express amount of substance concentration with units of molarity (M) (SI units: mmol/L).
