## Supplementary material for "CCQM-P199b: Interlaboratory comparability study of SARS-CoV-2 RNA copy number quantification": CCQM P199b supplementary file Appendix L

### APPENDIX L: RNA secondary structure analysis

Predicted secondary structure of the SARS-CoV-2 *N* gene was investigated in silico as part of comparison of study results for alternative assays.

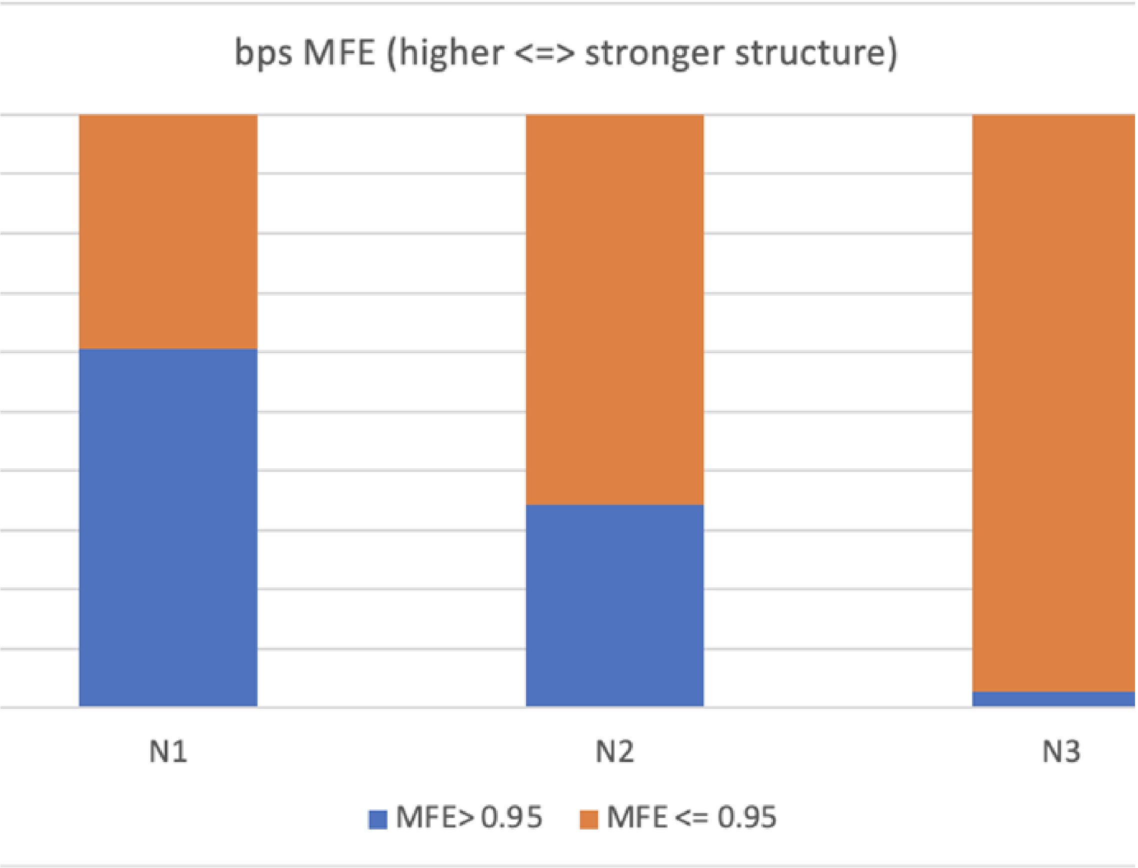


Figure L-1: Minimum free energy of intramolecular structure: CDC N1, N2, N3 assay amplicon regions. Minimum free energy (MFE) was calculated to be higher for CDC N1 amplicon region compared to CDC N2 and N3. RNAfold analysis (1) was performed by Laboratory 2.

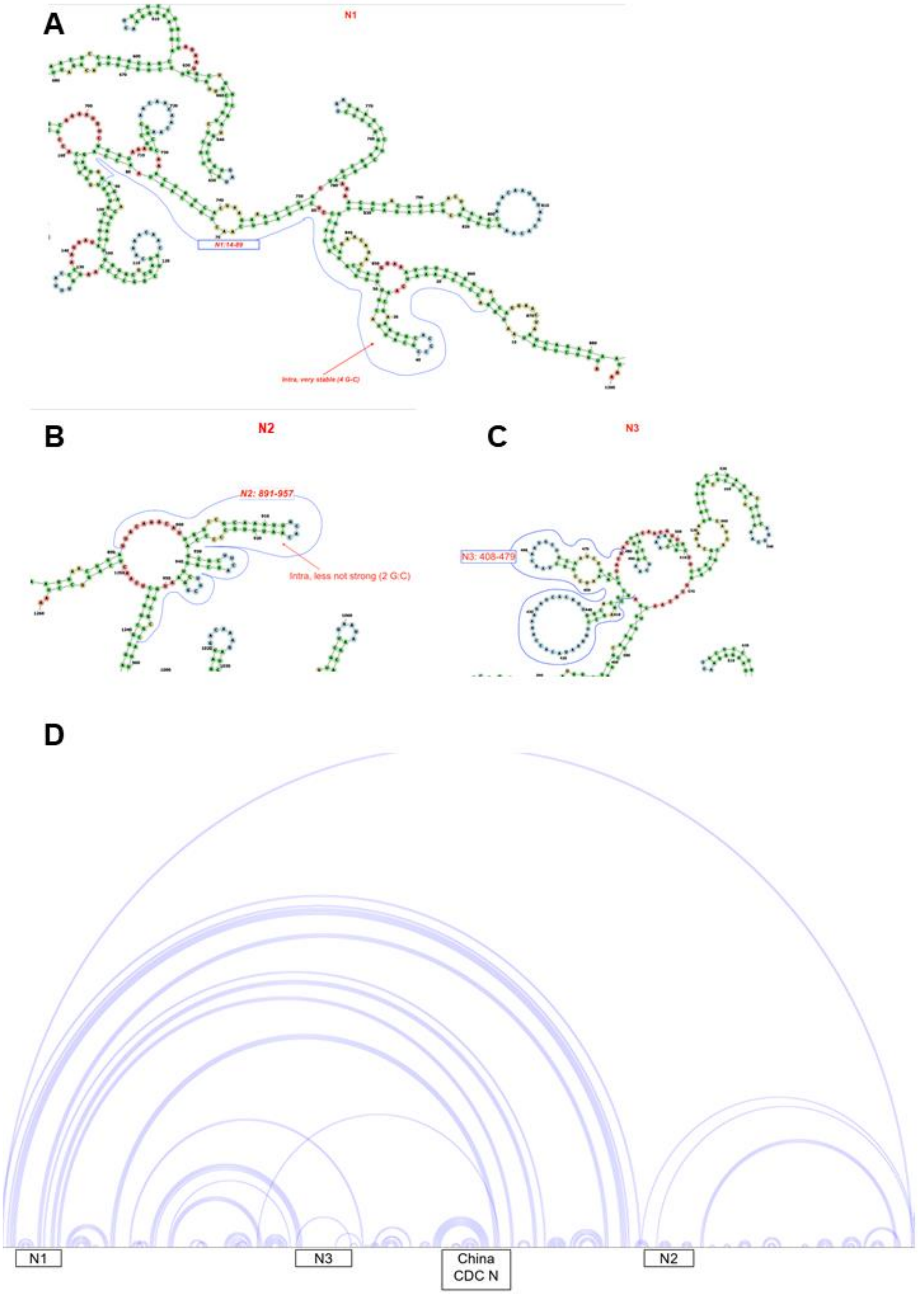


Figure L-2: Secondary structure prediction: *N* gene. Intramolecular interactions shown for CDC N1, N2 and N3 amplicon-containing regions (A-C respectively) using RNAFold by JRC-Geel (as Figure L1) and full N gene (D) using IPknot (2) and VARNA version 3.8 (3) performed by Laboratory 19. Location of CDC N1, N2 and N3 and China CDC N assays are shown.

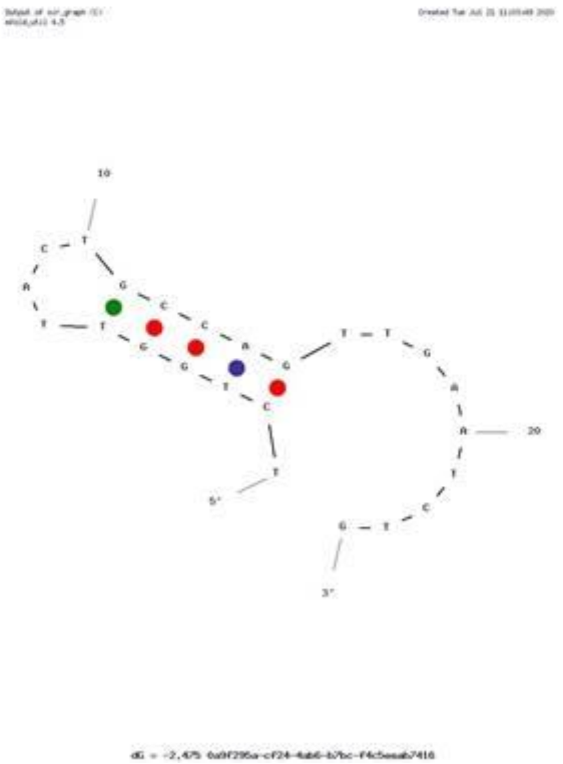


Figure L-3: Intramolecular hairpin prediction in CDC N1 assay reverse primer. Intramolecular interaction was predicted using mfold (4) and was confirmed using the Oligo Analyzer web tool from Integrated DNA Technologies (IDT) (5).

### REFERENCES

1. Gruber AR, Lorenz R, Bernhart SH, Neuböck R, Hofacker IL. The Vienna RNA Websuite. Nucleic Acids Res. 2008;36(suppl_2):W70-W4.
2. Sato K, Kato Y, Hamada M, Akutsu T, Asai K. IPknot: fast and accurate prediction of RNA secondary structures with pseudoknots using integer programming. Bioinformatics (Oxford, England). 2011;27(13):i85-i93.
3. Darty K, Denise A, Ponty Y. VARNA: Interactive drawing and editing of the RNA secondary structure. Bioinformatics (Oxford, England). 2009;25(15):1974-5.
4. Zuker M. Mfold web server for nucleic acid folding and hybridization prediction. Nucleic Acids Res. 2003;31(13):3406-15.
5. Owczarzy R, Tataurov AV, Wu Y, Manthey JA, McQuisten KA, Almabrazi HG, Pedersen KF, Lin Y, Garretson J, McEntaggart NO, *et al.* IDT SciTools: a suite for analysis and design of nucleic acid oligomers. Nucleic Acids Res. 2008;36(Web Server issue):W163-9.
